# The RNA-binding protein landscapes differ between mammalian organs and cultured cells

**DOI:** 10.1101/2022.02.10.479897

**Authors:** Joel I. Perez-Perri, Dunja Ferring-Appel, Ina Huppertz, Thomas Schwarzl, Frank Stein, Mandy Rettel, Bruno Galy, Matthias W. Hentze

## Abstract

System-wide approaches have unveiled an unexpected breadth of the RNA-bound proteomes of cultured cells. Corresponding information regarding RNA-binding proteins (RBPs) of mammalian organs is still missing, largely due to technical challenges. Here, we describe *ex vivo* eRIC (enhanced RNA interactome capture) to characterize the poly(A)RNA-bound proteomes of three different mouse organs. The resulting organ atlases encompass more than 1300 RBPs active in brain, kidney or liver. Nearly a quarter (291) of these had formerly not been identified in cultured cells, with more than 100 being metabolic enzymes. Remarkably, RBP activity differs between organs independent of RBP abundance, suggesting organ-specific levels of control. Similarly, we identify systematic differences in RNA binding between animal organs and cultured cells. The pervasive RNA binding of enzymes of intermediary metabolism in organs points to tightly knit connections between gene expression and metabolism, and displays a particular enrichment for enzymes that use nucleotide cofactors. We describe a generically applicable refinement of the eRIC technology and provide an instructive resource of RBPs active in intact mammalian organs, including the brain.

## Introduction

RNA-binding proteins (RBPs) constitute a versatile ensemble of proteins that play key roles in fundamental biological processes. They orchestrate the life cycle of messenger RNAs, from their synthesis in the nucleus to their translation and decay in the cytoplasm, and are thus essential for shaping cellular proteomes. They are also essential for the processing, function and decay of all other classes of RNA. There is growing evidence that protein activity can conversely be regulated by RNA (Hentze et al., 2018). Illustrating the importance of RBPs for cellular homeostasis, numerous diseases including neurological disorders and cancer have been linked to RBP misfunction (Gebauer et al., 2021).

Unbiased, system-wide approaches have paved the way for the determination of the composition, subcellular distribution and dynamics of RNA-bound proteomes (Baltz et al., 2012; Castello et al., 2012; Perez-Perri et al., 2018; Queiroz et al., 2019; Trendel et al., 2019; Urdaneta et al., 2019). These methods start with the crosslinking of RBPs to RNA *in cellulo* to stabilize RNA-protein interactions that occur within the native cellular environment. Irradiation of cells with ultraviolet (UV) light has been widely used because, unlike chemicals such as formaldehyde, UV exposure only exceptionally promotes protein-protein crosslinking and hence selects for direct RNA-protein interactions. The crosslinked RNA-protein complexes are subsequently isolated under highly stringent, denaturing conditions. In RNA interactome capture (RIC), oligo(dT)-coated magnetic beads are used to select polyadenylated transcripts together with their crosslinked RBP partners (Baltz et al., 2012; Castello et al., 2012). In an enhanced version called eRIC, the capture probe is modified with locked nucleic acids (LNA) improving specificity and the signal-to-noise ratio (Perez-Perri et al., 2021; Perez-Perri et al., 2018). Methods to isolate the whole RNA-bound proteome regardless of RNA biotype have also been developed (Bao et al., 2018; Huang et al., 2018; Queiroz et al., 2019; Trendel et al., 2019; Urdaneta et al., 2019). After the different capture strategies, the RNA-bound polypeptides are retrieved and analysed by mass spectrometry.

With only few exceptions from non-mammalian model organisms such as *Drosophila* embryos (Sysoev et al., 2016; Wessels et al., 2016), *Caenorhabditis elegans* (Matia-Gonzalez et al., 2015), zebrafish (Despic et al., 2017), and plants (Marondedze et al., 2016; Reichel et al., 2016; Zhang et al., 2016), RBP profiling methods have principally been used to study the RNA-bound proteomes of unicellular organisms or cultured cells. The lack of knowledge regarding mammalian organs and tissues is owed to technical limitations. The low penetration depth of UV light into biological specimen (Duck, 1990) limits the applicability of UV crosslinking in large multi-cellular organisms. A first approach to characterize the RNA-bound proteome of a mammalian organ was recently reported, which identified a relatively narrow set of 119 RBPs active in mouse liver (Na et al., 2021). This study used formaldehyde crosslinking and hence required measures to reduce contamination from protein-protein crosslinking. A method for the sensitive and specific detection of the RNA-bound proteomes of mammalian organs is thus still missing and the subject of this report.

We adapted the stringent eRIC protocol by using cryosectioning to comprehensively characterize the poly(A) RNA-bound proteomes of brain, liver, and kidneys from the house mouse *Mus musculus*. Our work represents the first in-depth profiling of RBPs from mammalian organs, refining the scope of RBPs, and revealing remarkable differences in RBP activity between organs as well as between organs and cultured cells.

## Results

### Refinement of eRIC to characterize the poly(A) RNA-bound proteomes of mammalian organs

The poor penetration of UV light through biological material (Duck, 1990) restricts UV crosslinking of RNA-protein contacts to the surface of intact tissues and organs. To overcome this limitation and to determine the RNA-bound proteomes of murine liver, brain and kidneys, we flash froze intact dissected organs in liquid nitrogen. The frozen organs were sectioned in a cryostat (thickness 30 µm) at low temperature, and the sections were transferred onto glass slides placed on a metal surface in direct contact with dry ice (Figure 1A). Subsequently, the slices were exposed to UV light (λ =254 nm) at a dose of 1 J/cm^2^, following titration experiments to maximise RNA-RBP crosslinking efficiency while minimizing sample heating and preserving RNA integrity (not shown). Immediately after UV irradiation, the samples were directly scraped into denaturing lysis buffer. To control for (crosslink-independent) background, every other organ slice was lysed without prior exposure to UV. Subsequently, the highly performant eRIC protocol for cultured cells was applied, starting with the isolation of poly(A) RNAs with magnetic beads conjugated to LNA-modified oligo(dT) probes, followed by extensive washes to remove non-crosslinked proteins, RBP elution by Rnase treatment, and finally RBP identification by mass spectrometry (Perez-Perri et al., 2021; Perez-Perri et al., 2018). A small fraction of the washed beads was subjected in parallel to heat treatment to recover and analyse the captured RNA (Figure 1A).

**Figure 1.**
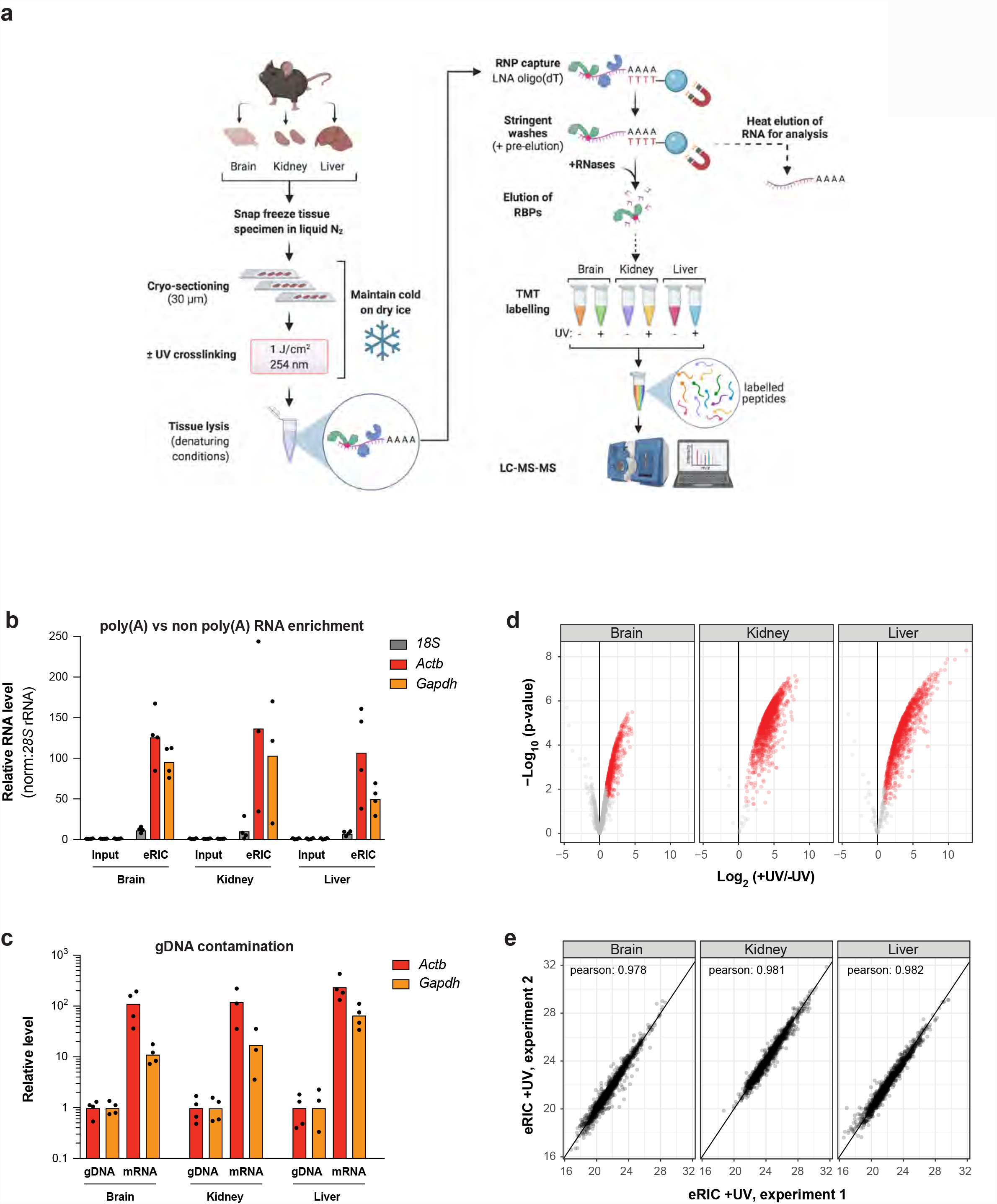
A method for the specific determination of the RNA-bound proteomes of mammalian organs. (A) Schematic representation of *ex vivo* eRIC (enhanced RNA interactome capture) applied to organs. Intact flash-frozen organs are sectioned into 30 µm slices amenable for UV irradiation. Following UV cross-linking (indicated by a red dot), tissue sections are lysed under denaturing conditions. RNA-binding proteins (RBPs) bound to polyadenylated RNA are subsequently isolated under highly stringent conditions using an LNA-modified oligo(dT) probe coupled to magnetic beads (Perez-Perri et al., 2021; Perez-Perri et al., 2018). A fraction of the isolated material is used for RNA analysis. The rest is subjected to RNase digestion to retrieve RBPs. Following solid-phase-enhanced sample-preparation (SP3) (Hughes et al., 2014; Hughes et al., 2019), peptides subjected to tandem mass tag (TMT) labelling are multiplexed and analyzed using LC-MS/MS (liquid chromatography/tandem mass spectrometry). Created with BioRender.com. (B-E) *ex vivo* eRIC was used to characterize the RNA-bound proteomes of brain, kidney and liver from adult C57BL6/J mice. (B) RT-qPCR analysis of *18S* rRNA as well as *Actb* and *Gapdh* mRNA abundance in eRIC eluates versus input, demonstrating enrichment of mRNA. Values are expressed relative to the respective input (input mean corresponds to 1.0). (C) qPCR analysis of mRNA versus genomic DNA (gDNA) for the housekeeping genes *Actb* and *Gapdh*, showing that gDNA contamination is minor. (D) Volcano plots showing significant enrichment (red dots) of RBPs in UV crosslinked over non-irradiated samples. The combined *ex vivo eRIC* data from brain, kidney and liver reveal more than 1300 RBPs (see Table S1). (E) Scatter plots comparing the normalized signal sums in ex vivo eRIC eluates obtained from independent experiments performed with distinct animals.

One key benefit of the LNA-based eRIC procedure for cultured cells that drove our choice for this enrichment method for organ samples is the high specificity and low background with the selective capture of poly(A)-transcripts and minimal genomic DNA (gDNA) contamination (Backlund et al., 2020; Perez-Perri et al., 2018). Capillary electrophoresis analysis of the RNA captured from the three tissues confirms the profound enrichment of mRNA, combined with a strong depletion of the highly abundant non-poly(A) RNAs that otherwise prevail, such as rRNA and tRNA (Figure S1A). As previously observed with cultured cells (Backlund et al., 2020; Perez-Perri et al., 2018), 18S rRNA is profoundly reduced but not completely eliminated, possibly due to the presence of a sufficiently long poly(A) stretch recognized by the capture probe, or by the interaction of the 18S rRNA with complementary sequences within poly(A) RNA. The capillary electrophoresis data were further corroborated by quantitative reverse transcription PCR (qRT-PCR) analyses performed with equal amounts of RNA from eRIC eluates and inputs, showing a 50- to 100-fold enrichment of two housekeeping mRNAs (*Actb, Gapdh*) over 18S rRNA (Figure 1B). Importantly, the direct qPCR analysis of eRIC eluates without prior reverse transcription step revealed that *Actb* and *Gapdh* cDNAs are at least 10-100 times more abundant than DNA from the same loci, indicating that gDNA contamination is minimal (Figure 1C).

To obtain sufficient material for the proteomics analyses, we combined the eRIC eluates obtained from the respective organs of two mice. The analyses were conducted in duplicate for each organ studied. For the no crosslink controls, organ sections from four mice were pooled to generate one unique no crosslink eRIC sample per organ studied. To reduce technical variability, RBP peptides were tandem mass tag (TMT)-labeled, multiplexed, and analyzed in a single liquid chromatography/tandem mass spectrometry (LC-MS/MS) run (Figure 1A). Total proteomes were also determined from input fractions for cross comparison with the poly(A) RNA-binding proteins.

High confidence eRIC hits were defined as proteins significantly enriched in eRIC eluates from UV-irradiated samples compared to the corresponding no-UV controls (fold difference > 2, false discovery rate (FDR) < 0.05). A total of 622, 1345 and 1238 eRIC hits were identified, respectively, from brain, kidney and liver (Figure 1D; Table S1), with remarkable reproducibility between the two independent experiments performed (Figure 1E). Of note, total (input) versus RNA-bound (eRIC eluates) protein samples correlate poorly, indicating specific enrichment of RBPs (Figure S1B).

Overall, the eRIC hits identified in the three organs studied are enriched for protein domains typically associated with RNA binding such as RNA-recognition motif (RRM), nucleotide-binding alpha-beta plait, DEAD/DEAH box helicase, and K homology (KH) (Figure 2A), as expected for a comprehensive set of RBPs. A gene ontology (GO) analysis also shows that eRIC hits are associated with terms mainly related to RNA metabolism, including translation (GO biological process), ribonucleoprotein complex (GO molecular function) or mRNA binding (GO cellular component) (Figure 2B; Table S5). Moreover, most eRIC hits belong to the RNA-related protein groups “nucleic acid binding protein” and “translational protein” according to the PANTHER classification system (Mi et al., 2021) (Figure 2C). These enrichments indicate that eRIC captures the core mRNA-bound proteome from intact organs. However, and in agreement with previous RBP profiling studies from cultured cells, the mRNA-bound proteome of mouse organs also includes many proteins that lack direct relationship to RNA metabolism a priori, as discussed below.

**Figure 2.**
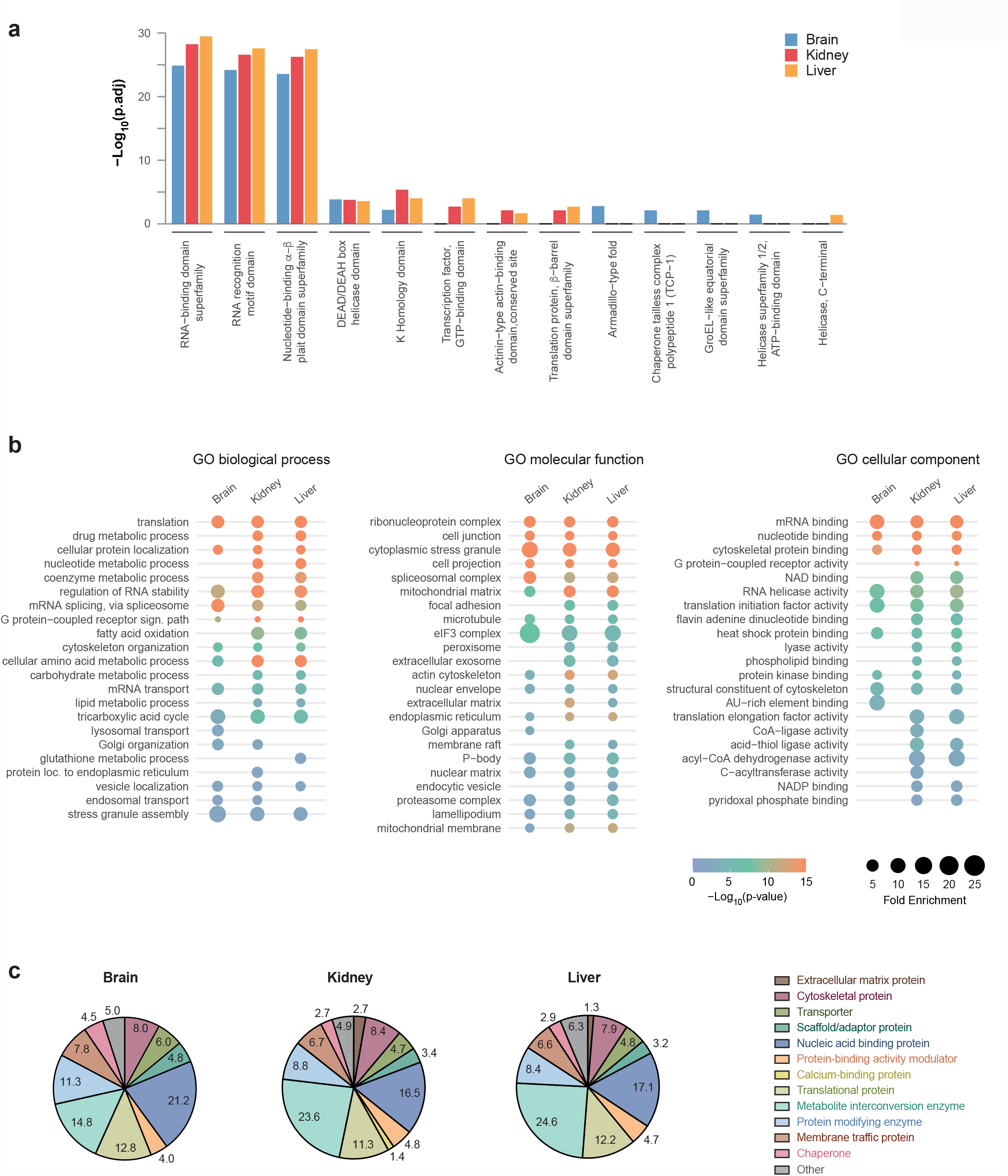
Characteristic features of the RBPs from mouse brain, kidney and liver. (A) Protein domains highly represented among *ex vivo* eRIC hits. (B) Gene Ontology (GO)-term enrichment analysis. Selected GO terms corresponding to biological process, molecular function, and cellular component are displayed (see Table S5 for the full list of GO terms). (C) Protein class distribution among the three tissues studied based on the PANTHER protein class ontology (Mi et al., 2021).

Taken together, *ex vivo* eRIC captures poly(A) RNA and crosslinked RBPs with high specificity and reproducibility from intact mouse brain, liver and kidneys, enabling the reliable, in depth characterization of whole organ poly(A) RNA-bound proteomes.

### Global control of RBPs in an organ-specific way

We next compared the poly(A) RNA-bound proteomes of brain, kidney, and liver tissues to each other. In total, 588 active RBPs are shared between the three organs, and an additional 648 active RBPs were identified both in kidney and liver (Figure 3A). 77 RBPs were solely detected in kidney, and, surprisingly, only very few RBPs were exclusively identified in brain or liver. This distribution suggests that the MS analysis may have been biased towards the detection of RBPs active in the kidney. Indeed, the averaged normalized TMT reporter ion signal (signal sum) in eRIC eluates of the proteins that scored as RBPs in at least one organ is 1.0e7, 5.3e7 and 1.4e7 in brain, kidney and liver, respectively (Figure 3B, top panel; Table S2), representing a mean eRIC intensity 5.3 and 3.8 times larger in kidney relative to brain and liver, respectively, and 1.4 larger in liver compared to brain. Hence, recovery of crosslinked RBPs differs substantially across organs, with kidney >> liver > brain. These marked differences in RBP binding across organs are not explained by differences in RBP expression. To the contrary: the mean RBP abundance is similar in kidney and liver, and only marginally lower in brain (Figure 3B, middle panel). Hierarchical clustering confirms that with few exceptions (Figure 3C, e.g. clusters 1 and 2), active RBPs are captured predominantly from kidney, irrespective of their relative expression levels across organs (Figure 3C, e.g. clusters 5, 7 and 10). The observed differences in RBP capture efficiency can neither be explained by disparities in the quantity (Figure 3B, bottom panel), integrity and/or purity of the captured RNA (Figures 1B,C and Figure S1A).

**Figure 3.**
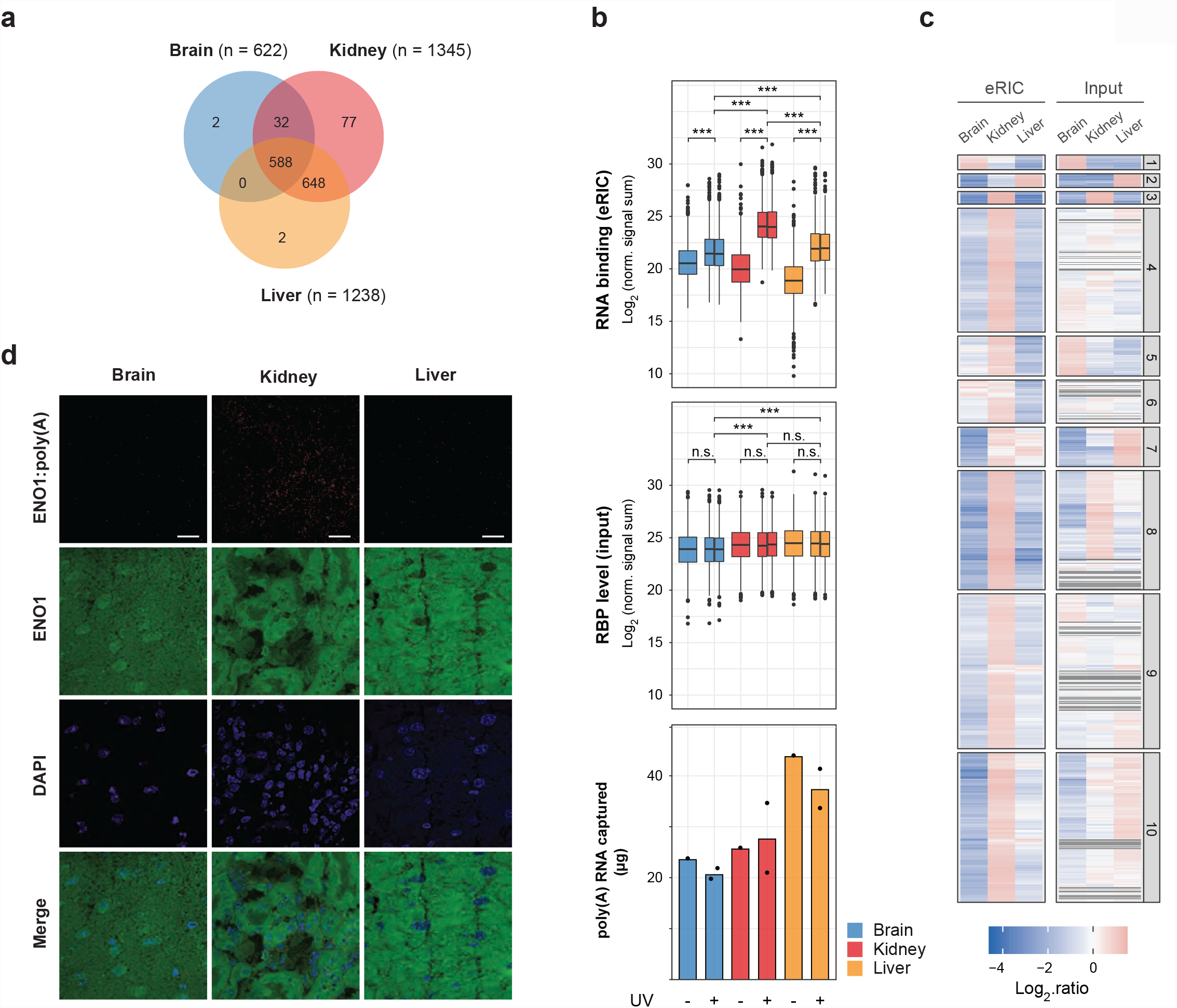
Comparative analysis of RBPs from mouse brain, kidney, and liver. (A) Venn diagram showing the number of shared and organ-specific RBPs identified. (B) Normalized signal sum of identified RBPs in *ex vivo* eRIC eluates (upper panel) and the corresponding input samples (middle panel) for each organ analyzed. Center lines indicate the median, box borders represent the interquartile range (IQR), and whiskers extend to ±1.5 time the IQR; outliers are shown as black dots (pairwise comparisons using t-test with FDR correction, ***p.adj < 2e-16; n.s.: not significant). Bottom panel: amount of poly(A) RNA isolated from each organ by eRIC. Note that the retrieved mass of protein (upper panel) differs extensively across the tissues analyzed (with kidney >> liver > brain) and does not necessarily correlate with the mass of RNA recovered (bottom panel) (see also Table S2). (C) Hierarchical clustering and heatmap of the RBPs identified in brain, kidney and liver, showing protein abundance in eRIC eluates (left columns) and inputs (right columns) across the three organs. (D) Representative images of the proximity ligation assay (PLA) for interactions of ENO1 and poly(A) RNA, nuclear staining (DAPI) and ENO1 immunofluorescence in brain, kidney and liver. Scale bar, 20 µM. See quantification in Figure S2B.

We realized that the disparities in RBP capture could result from organ-specific differences in UV crosslinking efficacy. To address this concern, we established an orthogonal, crosslinking-independent approach to assess the poly(A) RNA-binding activity of RBPs *in situ*. We used a proximity ligation assay (PLA) adapted to detect protein-RNA interactions using a specific antibody against the RBP of interest and a biotinylated DNA probe complementary to the interacting RNA (Roussis et al., 2016). To identify interacting poly(A) transcripts irrespective of their sequence, we generated biotinylated oligo(dT) probes anchored to the 3’UTR/poly(A) boundaries via two randomised nucleotides at its 3’end (Figure S2A).

For analysis, we strategically selected four RBPs that displayed higher RNA-binding activity in kidneys although their overall expression levels, judged by both proteomics and immunofluorescence, are similar or even lower in kidneys than in brain and liver (Figure S2B, left two panels). These include a classical RBP, the ATP-dependent RNA helicase DDX6, and three non-canonical RBPs: the glycolytic enzymes Enolase 1 (ENO1) and Pyruvate kinase (PKM), and the membrane-associated amino acid transporter SLC3A2 (a.k.a. 4F2 cell-surface antigen heavy chain). Confirming the *ex vivo* eRIC data (Figure S2B, third panel from the left), all four RBPs show stronger poly(A) RNA binding in kidneys than in liver and brain using the PLA (Figure 3D and Figure S2B, right panel). Thus, kidney RBPs appear to be highly active, and the widespread differences detected by eRIC are likely biologically determined rather than technical artifacts.

Thus, poly(A) RNA-protein interactions are globally controlled in an organ-specific manner.

### Surveying the organ-specific regulation of individual RBPs

Our study revealed widespread differences in the overall association of proteins with poly(A) RNA across brain, kidney and liver tissues. We next asked whether and how the binding of individual RBPs to poly(A) RNA differs between organs. For this, we analyzed the *ex vivo* eRIC data this time assuming equal mean signal intensity across samples (Figure S3A, Table S3). As before, we processed proteomics data from both eRIC eluates and total proteomic input to assess whether differences in eRIC signal intensity are due to differences in specific RNA-binding activity or in RBP levels. Hierarchical clustering shows similar patterns of protein abundance in eRIC eluates and total proteomic input for the majority of RBPs (Figure S3B). This suggests that differential RBP binding to RNA across mouse organs most commonly results from differential RBP expression. This standard pattern has however numerous exceptions, and dozens of RBPs exhibit differential RNA binding without commensurate changes in overall protein abundance (Figure S3C, Table S3).

These results reveal differences in the activity of individual RBPs between brain, kidney, and liver beyond the global effects addressed above.

### eRIC uncovers organ RBPs not previously detected in cultured cells

Previous RIC studies performed with cultured cells have systematically identified hundreds of RBPs that lack recognizable RNA-binding domains (RBDs) or RNA-related functions (reviewed in (Hentze et al., 2018). In mouse organs, less than half of the proteins captured by *ex vivo* eRIC have been formerly annotated as RBPs (Figure 4A, left panel), and not more than one fifth bear a known RBD (Figure 4A, second panel from the left). This is also consistent with the fact that many organ RBPs are associated with biological processes and molecular functions not directly linked to RNA biology (Figure 2B,C).

**Figure 4.**
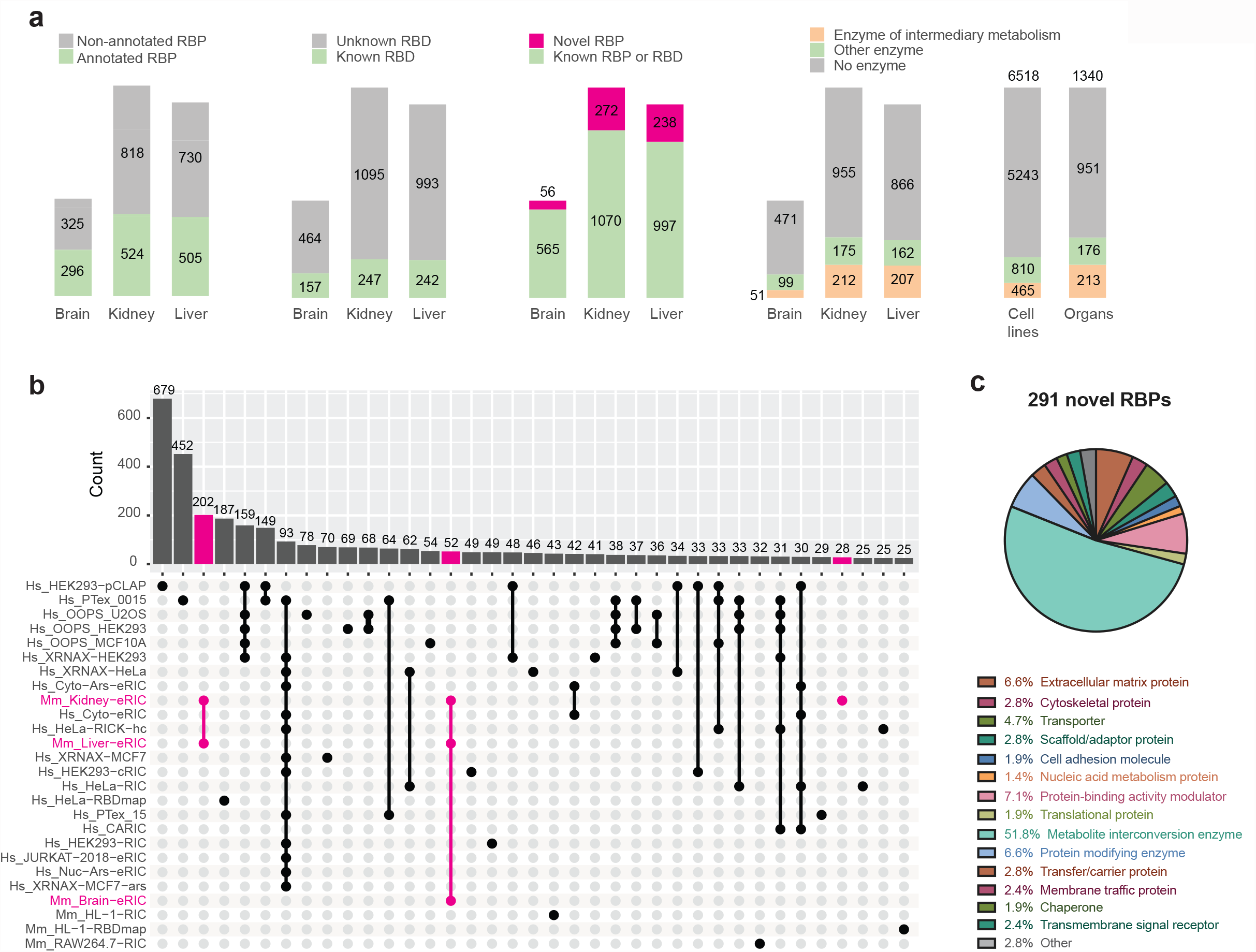
*Ex vivo* eRIC uncovers many novel RBPs. (A) For each organ, number of eRIC hits that are annotated RBPs, that bear a known RNA-binding domain (RBD), and that are novel RBPs (i.e. not previously reported as RBP or as bearing an RBD, and not identified in any published study). Right: Number of RBPs identified in organs or cell lines that are (metabolic) enzymes. (B) Upset plot representing the number of RBPs (y axis) shared between this study and published lists of mouse and human RBPs (only intersections comprising 25 or more RBPs are shown). (C) Protein class annotation of the novel RBPs identified in any of the three organs studied (based on PANTHER protein class ontology (Mi et al., 2021)). Note that compared to the overall RBP dataset (see Figure 2C), the class “translational protein” is under-represented, while the category “Metabolite interconversion enzyme” is over-represented.

We next compared the organ RBP dataset of 1349 proteins with the large integrated atlas of published RNA-binding proteomes derived from studies conducted in mouse or human cell lines (30 datasets encompassing a total of 6518 RBPs, based on the analysis of 12 cell lines and 10 different methods for the identification of poly(A)- or total RNA-binding proteins (Backlund et al., 2020; Baltz et al., 2012; Bao et al., 2018; Beckmann et al., 2015; Boucas et al., 2015; Castello et al., 2012; Castello et al., 2016; Conrad et al., 2016; Garcia-Moreno et al., 2019; Huang et al., 2018; Kwon et al., 2013; Liao et al., 2016; Liepelt et al., 2016; Mullari et al., 2017; Perez-Perri et al., 2018; Queiroz et al., 2019; Treiber et al., 2017; Trendel et al., 2019; Urdaneta et al., 2019). Because we expected that the large integrated RBP atlas (6518 RBPs) includes nearly all of the 1349 organ RBPs, we were surprised to find that nearly a quarter of our dataset of high-confidence organ RBPs (291) had not been formerly described as RBPs (Figure 4A, third panel from the left, Table S1). Of those 291 novel RBPs, 52 were active in all three organs examined, 202 in liver and kidney, and 28 in kidney alone (Figure 4B). These novel RBPs lack RBDs or RNA-related functions (Figure 4A). Remarkably, more than half of these are metabolic enzymes according to the PANTHER classification system; other represented protein classes include “protein-binding activity modulator”, “extracellular matrix proteins”, and “protein modifying enzymes” (Figure 4C). This is in sharp contrast with the “nucleic acid-binding protein” and “translational protein” classifications that prevail within the entire organ RBP dataset (Figure 2C).

Thus, we identified numerous organ RBPs previously not found in cultured cells.

### The RNA-bound proteomes of organs lack RBPs commonly identified in cultured cells

Conversely, we asked whether the RNA-bound proteomes of organs lack RBPs that have been commonly identified in cultured cells. Using the integrated RBP atlas as before, we noticed that 313 RBPs were commonly identified in cultured cells (at least 50% of previous reports), but absent from the organ RBP datasets (Figure S4A, B, Table S1). Around two thirds of these RBPs (204) could not be detected in the total proteomic samples (Figure S4B), suggesting that they are poorly or not expressed in organs. In a conservative approach, we excluded these proteins from further analysis, as if they were not (sufficiently highly) expressed (although some low expression RBPs can be enriched and detected in eRIC eluates even when undetected in total proteomic samples (purple dots in Figure S3C).

We thus focused on the 109 RBPs that were detected in proteomic inputs of brain, kidney and liver, but remained undetected by *ex vivo* eRIC in all three organs. Interestingly, most of these proteins correspond to bona-fide RBPs connected with nucleic acid metabolism and translation (Figure S4C), and are associated with GO terms related to “RNA binding” and “RNA processing” (Figure S4D, Table S6). Interestingly, the most highly enriched GO terms are linked to those aspects of RNA biology that do not involve polyadenylated RNAs (e.g. structural constituents of ribosomes, snoRNA binding, tRNA binding and ribosome biogenesis). For these RBPs, the explanation hence appears to be technical and reflect the excellent specificity of eRIC for poly(A) RNAs. Nonetheless, 26 out of the 109 proteins expressed in mouse organs but not identified by eRIC are associated with the GO term “mRNA metabolic process” (Figure S4D), raising the question of why these were not captured. We noticed that their expression level tends to be globally lower compared to that of eRIC hits related to the same GO term (Figure S4E, left panel). Yet, this is not the case for several proteins (e.g. Hnrnpa3, U2af2, Hnrnpa0) that show moderate to high expression in each of the organs tested (Figure S4E, right panel). Overall, this suggests that the binding of several proteins to poly(A) RNA may be enhanced in cultured cells relative to organs for reasons that remain to be explored.

### Striking prevalence of metabolic enzymes as RBPs in organs

The most striking finding regarding RNA-binding proteins in mouse organs is the prevalence of enzymes amongst these. Particularly, enzymes of intermediary metabolism interact with poly(A) RNA in mouse organs (Figure 4A, fourth panel from the left, Table S4), with at least 15% of kidney and liver RBPs and about 8% of brain RBPs being annotated as enzymes of intermediary metabolism. While RIC experiments with cultured cells already noted the systematic identification of metabolic enzymes (reviewed in (Castello et al., 2015; Hentze et al., 2018), the relative fraction did not exceed 7% of the total RBPs active in cultured cells (Figure 4A, first panel from the right).

An enrichment analysis shows that the >250 enzymes that bind poly(A) RNA in mouse organs are involved in various metabolic pathways connected to energy, carbon, fatty acid, or amino acid metabolism, with over 20 metabolic pathways being significantly enriched (Figure 5A, left panel). In comparison, only 10 of these metabolic pathways are also over-represented in the RBP datasets from cultured cells. The fraction of enzymes of a given pathway that bind poly(A) RNA can be rather high and exceed 50% (Figure 5A, right panel). This is the case for enzymes of the TCA cycle, pyruvate metabolism, or arginine biosynthesis, with at least 60% of the enzymes of these pathways being *ex vivo* eRIC hits. Taking pyruvate metabolism (Figure 5B), glycolysis (Figure S5A) and the TCA cycle (Figure S5B) as examples, enzyme-RBPs appear to be broadly distributed over the pathway as opposed to being involved in specific reactions.

**Figure 5.**
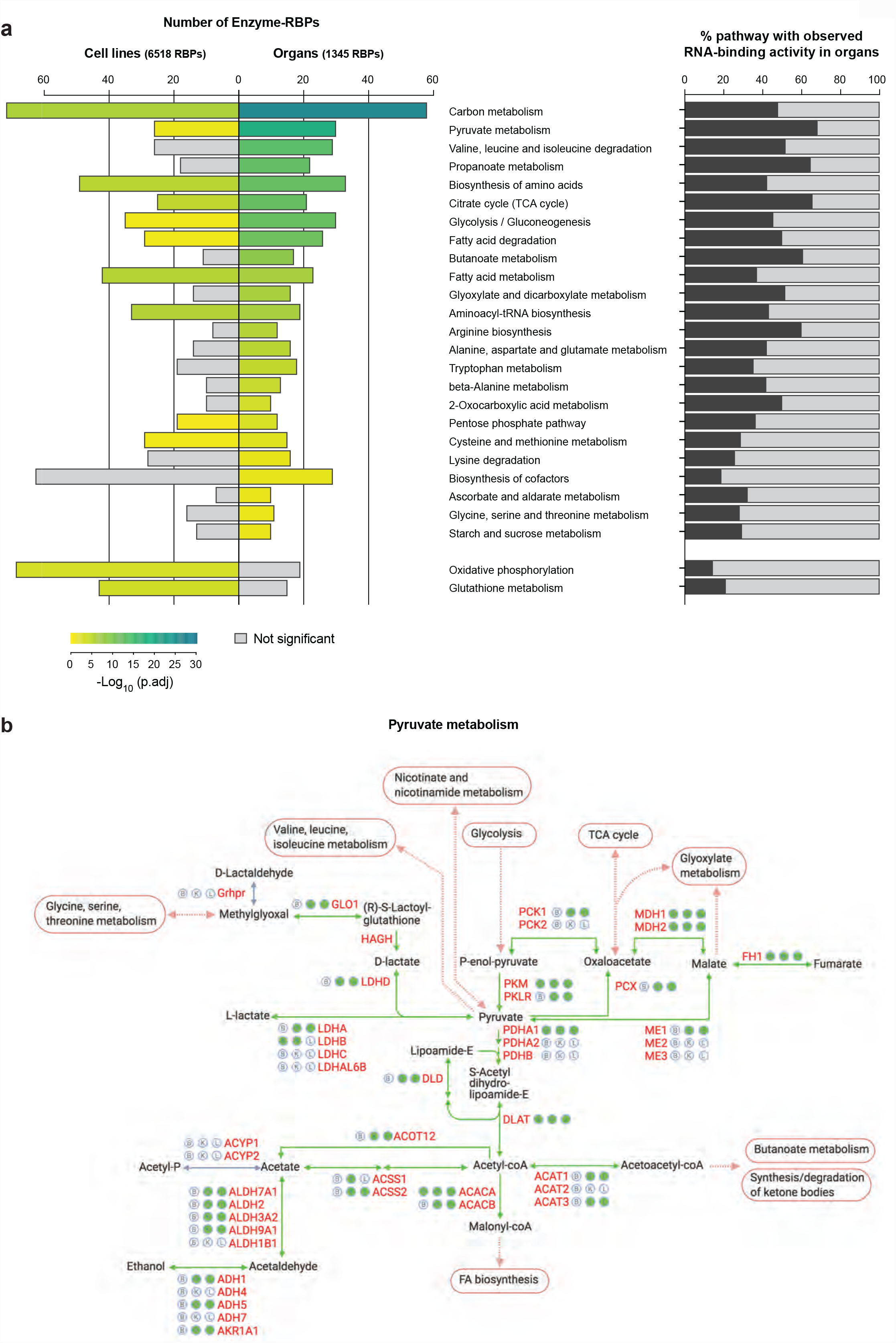
Pervasive RNA binding of metabolic enzymes in mouse organs. (A) Left: KEGG pathway enrichment analysis among the RBPs identified in at least one organ, or, for comparison, in at least one published RBP library in cultured cells. The most enriched metabolic pathways among organ RBPs are displayed; two pathways highly represented among cell culture RBPs but not among organ RBPs are shown at the bottom (functional enrichment analysis in g:Profiler with g:SCS multiple testing correction method). The size of the bars indicates the number of proteins identified as RBP for a given KEGG pathway. Right: Fraction of proteins in a given KEGG pathway that have been identified as RBP by *ex vivo* eRIC in brain, kidney or liver. (B) Schematic representation (based on the KEGG database) of pyruvate metabolism in mouse. Filled green circles next to protein names denote RNA-binding activity of the corresponding enzyme in brain (B), kidney (K) or liver (L); empty circles denote absence of evidence for RNA association. Reactions catalyzed by enzyme-RBPs are represented as green arrows.

Against this trend only two pathways, oxidative phosphorylation and glutathione metabolism, are highly represented in cell culture RBP libraries but not in the organ RBP datasets (Figure 5A). Possibly the exposure of cultured cells to supra-physiological concentrations of oxygen and chronic oxidative stress could explain this observation, although this is speculative at present.

In sum, the binding of intermediary enzymes to poly(A) RNA is even more common in organs than previously observed for cultured cells. It is widespread across multiple metabolic pathways, further highlighting tightly knit connections between gene expression and metabolism that await dissection in detail.

### Enzymes using nucleotide cofactors are highly enriched amongst organ RBPs

Which catalytic activities are most prevalent among the enzyme-RBPs identified in mouse organsã We selected all RBPs that belong to the protein group “metabolite interconversion enzyme” according to the PANTHER classification system (Mi et al., 2021) and noticed that oxidoreductases, transferases, and hydrolases appear to be most highly represented (Figure 6A, top; Table S4). While this corresponds to the overall occurrence of these enzymatic groups in the organs analyzed (Figure 6A, bottom), the catalytic activity “ligase” is clearly enriched among enzyme-RBPs, while hydrolases are under-represented (Figure 6B, top panel). A more refined analysis of enzyme subgroups shows an over-representation of two specific subtypes of oxidoreductases amongst the RBPs, namely dehydrogenases and peroxidases, while other oxidoreductase subtypes are not enriched (reductase, oxygenase) or even slightly under-represented (oxidase) (Figure 6B, bottom panel); there is also a modest enrichment for nucleotidyltransferases, and a tendency for under-representation of lipases and phosphatases (both hydrolase subtypes) amongst RBPs, as well as of kinases (transferase subtype) (Figure 6B, bottom panel).

**Figure 6.**
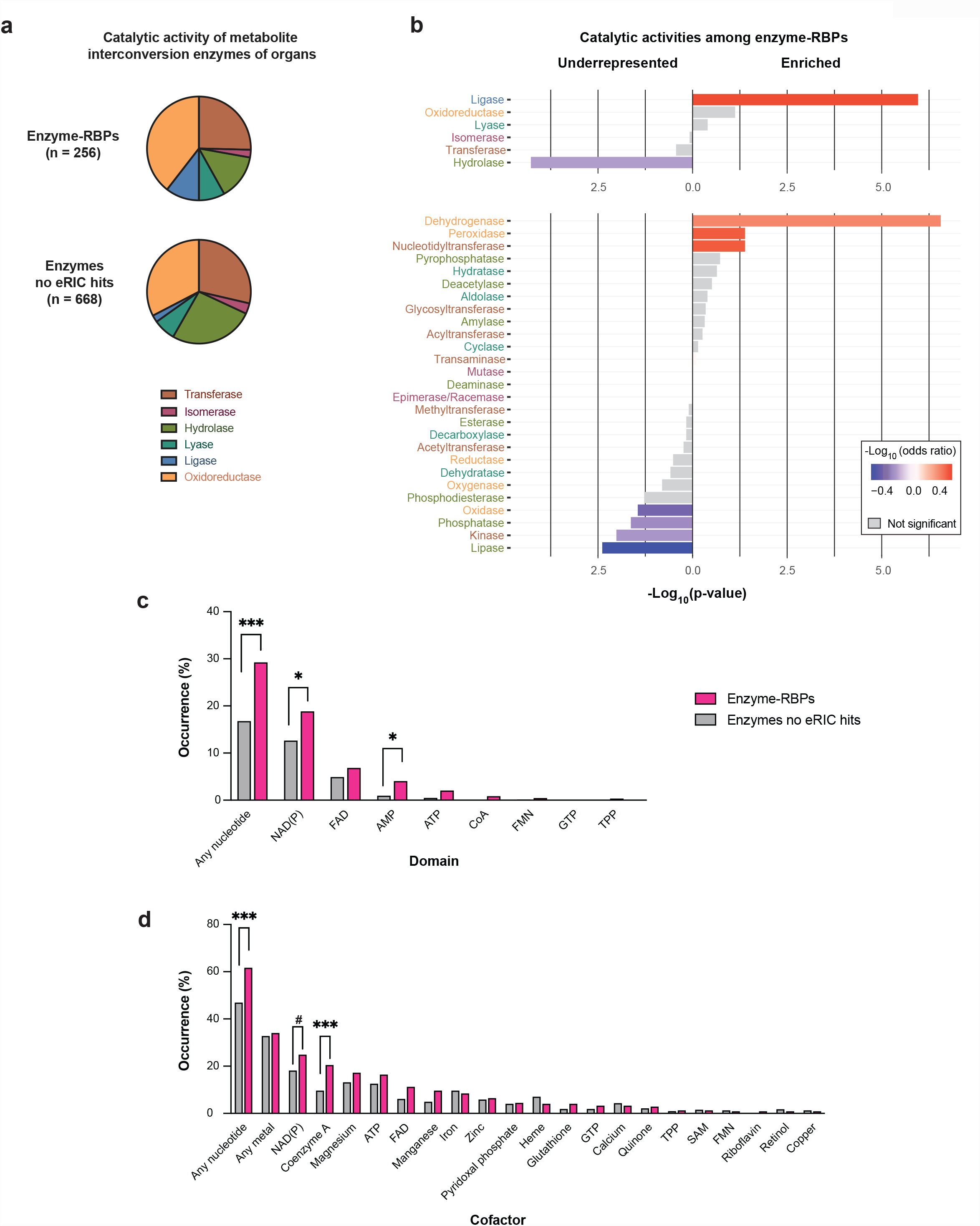
Enzyme-RBPs broadly interact with nucleotide co-factors. (A) Enzymes of intermediary metabolism (metabolite interconversion enzymes in PANTHER protein class (Mi et al., 2021)) identified as RBP in brain, kidney or liver (designated Enzyme-RBPs) were classified based on their catalytic activity; for comparison, the same classification was applied to enzymes of intermediary metabolism that were expressed in at least one organ but were not identified as RBP (designated no-eRIC hits). (B) Catalytic activity enrichment analysis among the enzyme-RBPs identified in mouse organs (Fisher’s exact test without correction for multiple comparison). The upper and bottom panels show, respectively, the main types and corresponding subtypes of enzymatic activities (indicated by the same distinct color in both panels). (C,D) The bar graphs indicate the proportion of enzymes expressed in organs and identified or not as RBPs that bear typical nucleotide binding-domains (C) or employ specific co-factors (D). *p.adj < 0.05, ***p.adj < 0.001, #p.adj = 0.065 (Fisher’s exact test with Benjamini-Hochberg correction).

Domains involved in mono- or di-nucleotide binding (e.g. ATP, NAD(P)+ and FAD) have been proposed as potential interfaces for RNA-binding (Castello et al., 2015; Hentze et al., 2018; Liao et al., 2016). Interestingly, the GO terms “NAD binding” and “NADP binding” are enriched among the whole repertoire of RBPs identified in kidney and liver (Figure 2B, right panel). Furthermore, nearly 30% of the enzyme-RBPs identified in mouse brain, kidney or liver bear at least one classifiable nucleotide binding domain (see Table S7 for domain classification), which is twice as frequent as the non-RBP enzymes expressed in the same organs (Figure 6C). Domains involved in NAD(P) (in ∼19% of enzyme-RBPs vs 12.6% of non-RBP enzymes) and AMP (4% vs 0.9%) binding are particularly frequent among enzyme-RBPs (Figure 6C).

In addition to these architectural features, some enzymes may correspondingly use co-factors biochemically via protein domains that are not yet characterized or still ill-defined. Therefore, we considered co-factor usage in addition to co-factor binding domains (Figure 6D), taking into account a relatively broad range of co-factors frequently used in enzymatic reactions, including metals. This analysis confirmed that enzyme-RBPs preferentially use nucleotide cofactors compared to non-RBP enzymes, especially NAD(P) (24.8% vs 18.1%) coenzyme A (20.4% vs 9.6%), ATP (16.4% vs 12.5%) and FAD (11.2% vs 6.1%). All other co-factors, including metals, are equally used by enzyme- and non-enzyme-RBPs (Figure 6D).

## Discussion

Since pioneering work about a decade ago, numerous proteome-wide analyses revealed that RNA-bound proteomes extend well beyond the previously known set of RBPs involved in core steps of RNA metabolism, at least in cultured cells and non-mammalian model organisms (Baltz et al., 2012; Castello et al., 2012; Despic et al., 2017; Matia-Gonzalez et al., 2015; Perez-Perri et al., 2018; Queiroz et al., 2019; Sysoev et al., 2016; Trendel et al., 2019; Urdaneta et al., 2019; Wessels et al., 2016). The resulting atlases of RBPs have paved the way for unexpected insights into RNA biology, especially regarding the role of protein-RNA interactions in cell biological processes such as e.g. metabolism, DNA methylation, protein ubiquitination or autophagy (Choudhury et al., 2017; Guiducci et al., 2019; Horos et al., 2019; Huppertz et al., 2020; Zhou et al., 2015). Nonetheless, corresponding data at the organismal level including an assessment of organ RBPs had not been reported, likely for technical reasons.

Very recently, an approach named FAX-RIC was used to characterize the RBPs of mouse liver (Na et al., 2021). Because the applicability of UV crosslinking is constrained by the limited penetration depth of UV light, the authors alternatively employed 4% formaldehyde to crosslink proteins to RNA. Although formaldehyde is advantageous over UV light for crosslinking proteins to double-stranded RNA, unlike UV crosslinking, formaldehyde additionally promotes protein-protein crosslinks and hence necessitates measures to distinguish protein-protein crosslinking from direct RNA-protein interactions. FAX-RIC identified a total of 119 RBPs in mouse liver, far fewer than anticipated, missing numerous “housekeeping RBPs” that are part of the core mRNA-bound proteome, and reflecting the non-trivial challenges associated with the adaptation of cell-based RBP discovery technologies to mammalian organs.

Here we combined organ cryosectioning and UV crosslinking with the strengths of cell-based eRIC. Our data show that *ex vivo* eRIC comprehensively interrogates the RNA-bound proteomes of intact organs, uncovering both shared and organ-specific features of RBPs from mouse brain, kidney, or liver. In comparison to FAX-RIC, eRIC uncovered more than 1200 high-confidence poly(A) RNA-binding proteins from the same organ (Figure 3). Thus, to the best of our knowledge, the data presented here represent the first comprehensive RBP profiling in intact mammalian organs.

While eRIC selects for proteins that bind poly(A) RNA, other RBP profiling methods such as OOPS, XRNAX, or PTex have been used to identify RBPs regardless of the RNA biotype that they bind to (Queiroz et al., 2019; Trendel et al., 2019; Urdaneta et al., 2019). These methods show a prevalence for proteins that interact with highly abundant, non-poly(A) RNA species such as rRNAs or tRNAs. While being more inclusive regarding the RNA biotype, they miss a significant fraction of less abundant RBPs that may exclusively or predominantly bind to the polyadenylated transcripts captured by eRIC, since poly(A) RNAs represent only a minor fraction (∼3-5 %) of the total cellular RNA. It is highly plausible that the cryosectioning-crosslink strategy we implemented here could also be used in combination with alternative approaches for total RNA purification. Furthermore, tissue cryosectioning and UV crosslinking could in principle also be followed by immunoprecipation of an RBP of interest to identify its target RNAs by CLIP-ing methods from organ samples.

Considering the enormous efforts to investigate the mammalian brain (International Brain Initiative, 2020; Yuste and Bargmann, 2017) and the essential roles of RBPs in neuronal functions (Ravanidis et al., 2018; Schieweck et al., 2021), the determination of CNS-/brain-related proteome-wide RBP datasets has been overdue. Therefore, we expect the data reported here to be particularly valuable for neurobiologists. The adapted *ex vivo* eRIC pipeline can in principle be applied to many other animal models or human samples, opening novel opportunities for studying RBP biology in health and disease.

We observed unexpected and marked differences between the RNA interactomes of organs versus cultured cells. On the one hand, *ex vivo* eRIC identified 291 RBPs in organs not detected in any of the 31 published RBP profiles of cell lines; this is remarkable, because the cumulative organ dataset is 5-times smaller than that of cell lines. On the other hand, dozens of previously identified RBPs were not detected by *ex vivo* eRIC despite being present in the tissues analyzed. Whether this reflects differences in RBP activity due to e.g. the metabolic environment (nutrient availability, oxygen levels, etc), or the activity of oncogenic pathways in cell lines versus normal cells of a healthy tissue is not known.

We noticed that the overall quantity of proteins crosslinked to a given quantity of poly(A) RNA differs substantially between organs, with protein binding to RNA being globally superior in kidney compared to liver and brain (Figure 3B,C). Most remarkably, only 4 of the 1349 proteins of the combined organ eRIC dataset were not identified as RBP in kidneys (Figure 3A). Total proteome analyses indicate these differences are not primarily due to organ-specific alterations of overall protein abundance (Figure 3B). Similarly, the integrity and purity of the RNA recovered by eRIC are comparable across all three tissues (Figure 1B,C; Figure S1A), excluding these technical artifacts. Furthermore, all *ex vivo* eRIC samples were analyzed in one single MS run. Organ-specific differences in UV crosslinking efficiency (due to e.g. dissimilarities in UV penetrance or the impact of UV-absorbing molecules) were also excluded as a technical reason: an orthogonal method (PLA) enabling the detection of protein-RNA complexes in situ independently of UV irradiation confirmed higher poly(A) RNA association in kidney of four unrelated RBPs that are actually expressed at a similar or even higher level in brain or liver (Figure S2 and Figure 3d). Organ-specific differences in global poly(A) RNA-protein interactions thus appear to represent a biological phenomenon and not a technical bias. The reasons for these remarkable differences and their functional consequences will be exciting to discover.

Previous RBP profiling studies revealed that a significant fraction of the RNA-bound proteomes of cells consists of intermediary metabolism enzymes (Castello et al., 2015; Hentze et al., 2018). *Ex vivo* eRIC shows that this is not a peculiarity of cultured cells. To the contrary, RNA-binding metabolic enzymes are even more prevalent in mammalian organs than cells (Figure 4A). While we cannot formerly exclude that some of the many individual enzyme-RNA interactions could be false positives, our observations further amplify the notion of physiologically important functions for these associations: RNA-binding enzymes have been shown to moonlight as trans-acting factors to control RNA fate, as illustrated by e.g. ACO1 or GAPDH (Chang et al., 2013; Constable et al., 1992); conversely, direct RNA binding to enzymes has been uncovered to riboregulate their catalytic activities (Huppertz et al., 2020; Wang et al., 2020; Zhou et al., 2015).

Our data indicate that the binding of enzymes to poly(A) RNA is widespread across and within multiple metabolic pathways, and overall even more prominent in organs than in cultured cells (Figure 5, Figure S5). Yet, compared to other metabolic pathways, enzymes involved in glutathione metabolism and oxidative phosphorylation appear to be more prone to RNA binding in cultured cells than in organs (Figure 5A). As glutathione is a major antioxidant, the overrepresentation of the glutathione metabolism pathway in RBP profiling studies of cultured cells might reflect the supra-physiological oxygen tension in cell culture and resulting chronic oxidative stress (Halliwell, 2003). Similarly, the over-representation of enzymes involved in oxidative phosphorylation may reflect differences in the activity of the electron transport chain. Performing eRIC studies in cells/organs subjected to either acute or chronic changes in e.g. oxygen levels, energy supply/demand may help address these points.

A large fraction (60%) of the enzyme-RBPs identified in this study bind mono- or di-nucleotide cofactors, with “NAD(P) binding” being particularly frequent among the kidney and liver RBP repertoires (Figure 6C, D). Furthermore, the detected enzyme-RBPs are highly enriched for specific enzymatic functions that rely on the use of nucleotide cofactors: ligases, which employ ATP or a similar energy donor, dehydrogenases that typically use NAD or NADP as proton acceptors, or CoA that we identify for the first time as a co-factor enriched in RBPs (Figure 6B). As ATP, NAD, FAD and CoA on the one hand and RNA on the other share the AMP handle as a structural feature, nucleotide-binding domains and RNA-binding interfaces may be evolutionarily connected (Hentze, 1994; Hentze et al., 2018).

Taken together, we report the first comprehensive atlases of RNA-binding proteins in mammalian organs. Far from only confirming the expected, our data reveal numerous surprising differences that form the basis for further explorations of the scope of RNA biology in mammalian physiology and disease. The method described here should be broadly applicable to other organs and organisms.

## Acknowledgments

B.G. is supported by grants from the Deutsche Forschungsgemeinschaft (GA2075/3-1, GA2075/5-1, GA2075/6.1). MWH appreciates valuable support from MOLIT (Heilbronn, Germany) and the Manfred Lautenschläger Foundation. We thank Alfredo Castello (Glasgow) as well as Thileepan Sekaran and Sudeep Sahadevan from the Hentze laboratory for their support with data analysis.

## Author Contributions

J.I.P.-P., B.G. and M.W.H. conceived the project and designed the experiments. B.G. and D.F.-A. conducted the experimental work with help from J.I.P.-P. and I.H. J.I.P.-P. and T.S. performed data analysis with help from B.G. M.R. and F.S. performed, respectively, the MS data acquisition and analyses. J.I.P.-P., B.G. and M.W.H. wrote the paper with input from all authors.

## STAR Methods

### Coupling of capture probes to beads

See (Perez-Perri et al., 2021) for a detailed step-by-step protocol. The capture probe (HPLC purified; Exiqon) contains a primary amine at the 5’ end, a flexible C6 linker, and 20 thymidine nucleotides in which every other base is a LNA: /5AmMC6/+TT+TT+TT+TT+TT+TT+TT+TT+TT+TT (+T: LNA thymidine, T: DNA thymidine). Prior coupling, carboxylated magnetic beads (50 mg/mL; Perkin Elmer, M-PVA C11) were washed three times with five volumes of 50 mM 2-(N-morpholino)ethanesulfonic acid (MES; Carl Roth, 4256.5) buffer, pH 6.0. The washed beads were then combined with a mix made of one volume of probe solution (100 µM in nuclease-free water, Ambion) and five volumes of freshly prepared *N*-(3-dimethylaminopropyl)-*N*′-ethylcarbodiimide hydrochloride (EDC-HCl; Sigma-Aldrich, E7750) solution (20 mg/mL in MES buffer).

The coupling reaction was performed at 50 °C for 5 h with constant agitation (800 rpm in a Thermomixer). The beads were then washed twice in phosphate-buffered saline (PBS) and incubated for 1 h at 37 °C in 200 mM ethanolamine pH 8.5 (with constant agitation at 800 rpm) to inactivate residual carboxyl groups. The coupled beads were finally washed three times with 1 M NaCl, and stored in 0.1% PBS–Tween at 4 °C until use.

### Mouse husbandry and organ dissection

Male mice on a homogenous C57BL6/J genetic background were housed under specific pathogen-free and light-, temperature- (21°C), and humidity (50-60% relative humidity)-controlled conditions. Food (Teklad, 2018S) and water were available ad libitum. The mice were sacrificed at 11 to 13 weeks of age by cervical dislocation; they were fastened 2h hours prior sacrifice (with access to water only). Organs were immediately harvested and flash frozen in liquid nitrogen. They were stored at -80°C until use.

### eRIC: Cryosectioning, UV irradiation and cell lysis

Organ sections (thickness 30 µm) were prepared in a Cryostat (Leica Biosystems, Leica CM3050 S) set to -20 °C and deposited onto SuperFrost glass slides (Carl Roth, H880); the glass slides were pre-cooled to -20°C in the cryostat chamber to maintain the samples at the lowest temperature possible. ∼10 sections were placed on a glass slide, and a total of around 150 sections were prepared for each sample per mouse (material of 2 mice were combined for each. Tissue sections on glass slides were then transferred onto metal plates placed in direct contact with dry ice to preserve sample integrity during UV irradiation. The tissue sections were exposed to UV light (λ = 254 nm) at a dose of 1 J/cm^2^ in a XL 1500 UV Spectrolinker (Spectronics Corporation). After UV exposure, organ sections were recovered by scraping directly into 15 mL ice-cold lysis buffer (see composition below) supplemented with protease inhibitors (Roche, 11873580001) and RNase inhibitor (1:1000, produced at the Protein Expression and Purification Core Facility, EMBL Heidelberg). For the non-crosslink control, every other section was lysed directly without exposure to UV. The lysates were pipetted up and down several times to dislodge tissue sections. They were finally passed once through a 25-Gauge needle (BD Microlance, 300400) and 7 times through a 27-Gauge needle (BD Microlance, 302200). The homogenates were snap frozen in liquid nitrogen and stored at −80 °C until use.

### eRIC: Capture of RNP complexes

See (Perez-Perri et al., 2021) for a reference step-by-step protocol. Cell lysates were thawed at 37 °C, incubated for 15 min at 45 °C, cooled down on ice, and the debris was pelleted 5 min at 16.000 x g at 4 °C. The supernatant was transferred to 15 mL DNA LoBind tubes (Eppendorf, 0030122208) and complemented with 5 mM extra of dithiothreitol (DTT; Biomol, 04020.100). 200 µL of each sample were taken as input, and the rest was mixed with 15 mg of capture probe-coupled beads previously equilibrated 3 times with 3 volumes of lysis buffer. The samples were incubated 1h at 37°C with gentle rotation. The beads were then collected on a magnet, and the supernatant was transferred to a fresh 15 mL DNA LoBind tube for a second round of capture. After the capture, the beads were transferred to 5 mL DNA LoBind tubes (Eppendorf, 0030108310) and washed first with the lysis buffer, and then twice with each of the buffers 1, 2 and 3 (see composition below). Each wash was performed with 5 mL of the corresponding buffer for 5 min at 37°C with gently rotation. A “pre-elution” step was performed by incubating the washed beads with 220 µL of nuclease-free water (Ambion) for 10 min at 40 °C and 800 rpm. The bead suspension was then divided into two aliquots: 200 µL were used for RNase-mediated elution of RNA-bound proteins, the rest (20 µL) was heat-eluted to recover the nucleic acid. The beads were collected on a magnet and the supernatant discarded. For RNase elution of proteins, the beads were resuspended in 150 µL of 1× RNase buffer (see composition below) containing 5 mM DTT, 0.01% NP40, ∼200 U RNase T1 (Sigma-Aldrich, R1003–100KU), and ∼200 U RNase A (Sigma-Aldrich, R5503). Following a 60 min incubation at 37 °C, 800 rpm, the beads were collected on a magnet, and the eluate was transferred to a fresh tube. The eluate was placed once more on the magnet to eliminate any remaining beads, and safely stored on ice. Heat elution of RNA was performed by incubating the aliquoted beads in 15 µL of water for 5 min at 95 °C, 800 rpm. The beads were immediately collected on the magnet, and the supernatant recovered as quickly as possible to avoid a drop in temperature (with the consequent re-capture of RNP complexes). Any trace of beads was removed by a second round of collection as explained. Corresponding eluates from the two consecutive rounds of capture were combined. Eluates obtained by RNase treatment (total volume: ∼400 µL) were supplemented with 0.05 % of SDS (2 µL of 10% SDS), concentrated at 45 °C to a volume of ∼100 µL in a SpeedVac, snap frozen, and stored at −80 °C.

Lysis buffer: 20 mM Tris-HCl (pH 7.5), 500 mM LiCl, 1 mM EDTA, 5 mM DTT, 0.5% (w/v) LiDS.

Buffer 1: 20 mM Tris-HCl (pH 7.5), 500 mM LiCl, 1 mM EDTA, 5 mM DTT, 0.1% (w/v) LiDS.

Buffer 2: 20 mM Tris-HCl (pH 7.5), 500 mM LiCl, 1 mM EDTA, 5 mM DTT, 0.02% (v/v) NP40.

Buffer 3: 20 mM Tris-HCl (pH 7.5), 200 mM LiCl, 1 mM EDTA, 5 mM DTT, 0.02% (v/v) NP40.

10× RNase buffer: 100 mM Tris-HCl (pH 7.5), 1.5 M NaCl

### RNA extraction, capillary electrophoresis, cDNA synthesis and real-time quantitative PCR

The concentration of the captured RNA (heat-eluted) was estimated using a NanoDrop spectrophotometer (Thermo Fisher Scientific). 10 ng of eluted RNA was analyzed using an Agilent 2100 Bioanalyzer System using the RNA 6000 Pico Kit, following the manufacturer’s instructions. Total RNA in the input was extracted using Trizol LS (Thermo Fisher) and analyzed the same way.

For RT-qPCR analysis of RNA, ∼100 ng of captured RNA versus input RNA were treated with DNase I (Thermo Fisher), and reverse transcribed using SuperScript III (Life Technologies) together with random hexamers (Life Technologies), following the manufacturer’s instructions. For qPCR analysis of DNA, the same reaction was performed in parallel but without the DNase and reverse transcriptase enzymes. Real-time qPCR was performed using a SYBR Green PCR Master Mix (Life Technologies, 4309155) together with a QuantStudio 6 Flex system (Life Technologies) and the following primers (5′ to 3′, forward: f, reverse: r): 28S rRNA (f: TTACCCTACTGATGATGTGTTGTTG, r: CCTGCGGTTCCTCTCGTA), β-actin (f: CGCGAGAAGATGACCCAGAT, r: TCACCGGAGTCCATCACGAT), GAPDH (f: GTGGAGATTGTTGCCATCAACGA, r: CCCATTCTCGGCCTTGACTGT) and 18S rRNA (f: GAAACTGCGAATGGCTCATTAAA, r: CACAGTTATCCAAGTGGGAGAGG)

### Sample preparation for mass spectrometry (MS) and TMT labelling

eRIC samples were concentrated using a SpeedVac apparatus, and treated with 10 mM DTT in HEPES buffer (50 mM HEPES, pH 8.5) for 30 min at 56ºC to reduce disulphide bridges in proteins. Reduced cysteines were then alkylated for 30 min at room temperature with 20 mM 2-chloroacetamide in HEPES buffer (protected from light). Samples were prepared using the SP3 protocol (Hughes et al., 2014; Hughes et al., 2019) and trypsin (sequencing grade, Promega) was added at an enzyme to protein ratio of 1:50. Following an overnight incubation at 37°C, the digested peptides were recovered in HEPES buffer applying two successive elution steps. The peptides were subsequently labelled with TMT10plex (Werner et al., 2014) Isobaric Label Reagent (ThermoFisher) according to the manufacturer’s instructions. The samples were combined and cleaned using an OASIS® HLB µElution Plate (Waters). Offline high pH reverse phase fractionation was carried out on an Agilent 1200 Infinity high-performance liquid chromatography system, equipped with a Gemini C18 column (3 µm, 110 Å, 100 × 1.0 mm, Phenomenex).

### Liquid chromatography with tandem mass spectrometry (LC−MS/MS)

An UltiMate 3000 RSLC nano LC system (Dionex) fitted with a trapping cartridge (µ-Precolumn C18 PepMap 100, 5µm, 300 µm i.d. × 5 mm, 100 Å) and an analytical column (nanoEase™ M/Z HSS T3 column 75 µm × 250 mm C18, 1.8 µm, 100 Å, Waters) was used. Trapping was carried out with a constant flow of trapping solution (0.05% trifluoroacetic acid in water) at 30 µL/min onto the trapping column for 6 minutes. Subsequently, peptides were eluted via the analytical column running solvent A (0.1% formic acid in water, 3% DMSO) with a constant flow of 0.3 µL/min, with increasing percentage of solvent B (0.1% formic acid in acetonitrile, 3% DMSO) from 2% to 8% in 4 min, from 8% to 28% for a further 104 min, from 28% to 40% in another 4 min, and finally from 40% to 80% for 4 min, followed by re-equilibration back to 2% B in 4 min. The outlet of the analytical column was coupled directly to an Orbitrap Fusion™ Lumos™ Tribrid™ Mass Spectrometer (Thermo Scientific) using the Nanospray Flex™ ion source in positive ion mode.

The peptides were introduced into the Fusion Lumos using a Pico-Tip Emitter 360 µm OD × 20 µm ID; 10 µm tip (New Objective) and an applied spray voltage of 2.4 kV. The capillary temperature was set to 275°C. A full mass scan was acquired with a mass range of 375-1500 m/z in profile mode in the orbitrap with a resolution of 120000. The filling time was set to a maximum of 50 ms with a limitation of 4×10^5^ ions. Data dependent acquisition (DDA) was performed with the resolution of the Orbitrap set to 30000, with a fill time of 94 ms and a limitation of 1×105 ions. A normalized collision energy of 38 was applied. MS^2^ data was acquired in profile mode.

### MS data analysis

IsobarQuant (Franken et al., 2015) and Mascot (v2.2.07) were used to process the acquired data, which was searched against the *Mus musculus* (UP000000589) Uniprot proteome database containing common contaminants and reversed sequences. The following modifications were included into the search parameters: Carbamidomethyl (C) and TMT10 (K) (fixed modification), Acetyl (Protein N-term), Oxidation (M) and TMT10 (N-term) (variable modifications). A mass error tolerance of 10 ppm and 0.02 Da was set, respectively, for the full scan (MS1) and MS/MS (MS2) spectra. Further parameters were: Trypsin digestion with a maximum two missed cleavages tolerated; a minimum peptide length of seven amino acids; at least two unique peptides required for protein identification. The false discovery rate was set to 0.01 on both the peptide and protein level.

The protein output files of IsobarQuant were processed in R (ISBN 3-900051-07-0). Raw TMT reporter ion intensities (‘signal_sum’) were first cleaned for batch effects using the ‘removeBatchEffects’ function of the limma package (Ritchie et al., 2015) and further normalized using the vsn (variance stabilization normalization) package (Huber et al., 2002). Two different normalization strategies were used, as indicated. In the first analysis, we estimated different normalization coefficients for each tissue and condition (+UV, -UV) of the eRIC samples. In the second approach, we estimated only a single normalization coefficient for +UV eRIC samples of the three tissues. For both types of analysis, proteins were tested for differential expression using the limma package. The replicate information was added as a factor in the design matrix given as an argument to the ‘lmFit’ function of the limma package. Proteins with a FDR smaller than 0.05 and a fold-change of at least 100 % were annotated as hits, and proteins with a FDR below 0.2 % and a fold-change of at least 50 % as candidates.

### eRIC hit classification, gene ontology (GO), and domain analysis

Mouse and human RNA interactome studies along with functional annotations were downloaded from the RBPbase (https://rbpbase.shiny.embl.de, v.0.2.0). The R package ‘ggupset’ was used for the visualization of overlaps between data sets with upset plots (Conway et al., 2017). Fisher’s Exact with independent hypothesis weighting (IHW) for multiple hypothesis testing correction was used for overrepresentation analysis of protein domain information from MouseMine (Motenko et al., 2015).

GO-term enrichment analysis in Figure 2B were conducted with AmiGO 2 (Carbon et al., 2009) (http://amigo.geneontology.org/amigo), using the following parameters: analysis type: PANTHER overrepresentation test (Released 20200407); Annotation version and release date: GO ontology database Released 2020-02-21; reference list: *Mus musculus* (all genes in database); annotation data set: GO biological process complete, GO molecular function complete or GO cellular component complete, as indicated; test type: Fisher’s exact with Bonferroni correction for multiple testing. Graphical representations were made with the ggplot2 R package (Ginestet, 2011). GO-term enrichment analysis in Figure S4D were performed in g:Profiler (https://biit.cs.ut.ee/gprofiler/gost) (Raudvere et al., 2019; Reimand et al., 2007), using the following parameters: organism: *Mus musculus*; statistical domain scope: only annotated genes; significance threshold: g:SCS threshold (tailor-made algorithm for multiple testing correction); data sources: GO molecular function and GO biological process. Due to space constrains, a selection of GO terms was included in Figure 2B and Figure S4D. The corresponding full lists of GO-enriched terms can be found in Tables S5 and S6, respectively.

PANTHER protein class analysis was performed using the PANTHER classification system (Mi et al., 2021) (http://www.pantherdb.org) (v.16.0). The percentages shown were calculated against the total number of protein class hits.

KEGG pathway enrichment analysis was performed in g:Profiler (Raudvere et al., 2019) using the following parameters: organism: *Mus musculus*; statistical domain scope: only annotated genes; significance threshold: g:SCS threshold (tailor-made algorithm for multiple testing correction); data source: KEGG. The results were manually curated to exclusively select pathways of intermediary metabolism.

### Analysis of nucleotide-binding domains and cofactors

These analyses were performed on proteins listed in the PANTHER protein class: “metabolite interconversion enzyme” (enzyme). InterPro domain and cofactor annotations were retrieved from the UniProt mouse database (release-2021_03). Protein domains were classified as shown in Table S7. Cofactors annotations were extracted from the sections “Catalytic activity”, “Cofactor” and “Keywords”. Enrichment of domains/cofactors was performed on enzymes that scored as eRIC hits in at least one organ, and compared to enzymes that were detected in inputs but not in eRIC eluates. The p-value was computed by Fisher’s exact test, and corrected for multiple testing by the Benjamini-Hochberg method.

### Proximity Ligation Assay (PLA)

The PLA was adapted from a previously described protocol (Roussis et al., 2016). An anchor probe was designed to target the 3’UTR/poly(A) boundaries of mRNAs. The probe is composed (5’ to 3’) of a 5’biotin tag [BtnTg], a poly(dT) 18mer followed by two random nucleotides (“NN”). The control probe lacks the biotin tag but is otherwise identical.

Mouse organs (kidney, liver, and brain) were obtained as described above. 10 µm-thick sections were prepared in a cryostat (Leica Biosystems, Leica CM3050 S), transferred onto SuperFrost Plus adhesion slides (Carl Roth), encircled by a hydrophobic boarder with a 2 mm-thick pap pen (Sigma Aldrich), and fixed with 4% paraformaldehyde at room temperature for 20 minutes. The fixed tissue sections were washed with PBS and permeabilized with PBS containing 1% BSA and 0.1% Triton X-100 at room temperature for 30 minutes. Tissue sections were then washed once for 5 minutes with 0.1 M Triethanolamine containing acetic anhydride and twice with PBS-T (0.02% Tween20). The sections were further washed twice with hybridization buffer (1x Denhardt’s solution, 0.1% (v/v) Tween20, 0.1% (w/v) CHAPS, 5 mM EDTA, 1 mg/mL RNase free tRNA, 100 µg/mL heparin) and then incubated in hybridization buffer containing 100 nM of probe in a wet chamber at 37°C overnight. The probe was boiled for five minutes at 95°C prior to addition. After hybridization, the sections were subjected to 5 minute washes first in 50% (v/v) deionized formamide/5xSSC (saline-sodium citrate), then in 25% (v/v) deionized formamide/1xSSC, 12.5% (v/v) deionized formamide/2xSSC, 2xSSC/0.1% (v/v) Tween-20, and finally 0,2xSSC/0.1% (v/v) Tween-20. Subsequently, the Duolink PLA Fluorescence protocol (Sigma) was followed using an anti-biotin antibody (mouse: 1:400; ab201341, Abcam) and either the anti-ENO1 (rabbit: 1:400; Proteintech, 11204-1-AP), anti-SLC3A2 (rabbit: 1:400; Santa Cruz Biotechnology, sc-9160), anti-DDX6 (rabbit: 1:400; Novus Biologicals, NB200-192) or anti-PKM1 antibody (rabbit: 1:400; Cell Signalling, D30G6) for detection of the protein–RNA signal. The primary antibodies were incubated at room temperature for 90 minutes. After the last wash of the Duolink PLA Fluorescence protocol, the slides were incubated at room temperature for 45 minutes with antibody diluent mixed with DAPI (final concentration of 0.1 µg/µL) and nanobodies targeting rabbit IgG coupled with Alexa488 (alpaca nanobody; Chromotek, srbAF488-1-100). The tissue sections were washed once with PBS-T and a glass cover was mounted using ProLong Diamond Antifade Mountant (ThermoFisher Scientific, P36961). The prepared slides were stored at 4°C until fluorescence microscopy.

Microscopy was performed using a LSM 780 Laser Scanning Microscope (ZEISS) equipped with an AxioCamera and a 63x/1.4 objective with immersion oil (Immersol 518F, ZEISS, 10539438, Lot No. 170201). The microscope was operated using the ZEN 2012 software (ZEISS). The DAPI signal was recorded in one plane to act as a reference for counting the number of cells. The PLA (Alexa 594) and the protein (Alexa 488 nanobody) signals were recorded as a Z-stack (10 pictures for 10 µm stack). Two images were taken per tissue section from brain, kidney, and liver from three different mice. The images were acquired as .lsm files using the same settings (gain, laser power, pinhole and offset) for the same protein and analysed using the Fiji software. The .lsm files were split into individual channels, the Z-stacks for the PLA signal and the Alexa 488 signal were projected into a single plane and the brightness was set to the same level in all images of the same channel to enable the comparison of the results. The individual channels were ultimately saved as .tiff files.

To count the PLA signals per cell for each of the different conditions, the CellProfiler Software version 4.1.3 was used. The range of the signal spot size was set to 8 to 20 pixels and the range of the nuclear size (DAPI signal) was set to 100 to 400 pixels. In both instances, the global threshold strategy minimum cross-entropy was used. The threshold smoothing factor was 1.3488 for the PLA signals and 20 for the nuclei. The Alexa 488 signal was used as an outline of the cells. Clumped objects were separated by intensity and objects touching the border of the images were discarded. The PLA signal per cell were counted by combining the information of the cellular outline (Alexa 488) and the DAPI signal. The statistical analysis of these results was performed using Graphpad Prism version 9 (one-way ANOVA with Tukey *post-hoc* test).

## Supplemental Information Titles and Legends

**Figure S1. Technical aspects of *ex vivo* eRIC**

(A) Representative capillary electrophoresis-based analysis of the RNA material isolated from inputs (top) or by *ex vivo* eRIC from brain, kidney and liver (bottom). The sharp peak of 25 nt corresponds to the marker. [nt] length of RNA in number of nucleotides, [FU] fluorescence units. (B) Normalized signal sums of eRIC eluates (y-axis) versus input samples (x-axis). Note the poor correlation.

**Figure S2. Validation of organ-specific differences in RNA binding by PLA**

(A) Schematic representation of the proximity ligation assay (PLA) for RNA-protein interactions (Roussis et al., 2016) employed to validate the organ-specific activities identified by *ex vivo* eRIC of four selected RBPs. Organ sections were fixed and subsequently incubated with i) a biotinylated “anchored” oligo(dT) probe, ii) antibodies raised in different species against the target RBP and biotin, and iii) secondary antibodies conjugated to PLA oligonucleotide probes. If the two probes are sufficiently close (<40 nm), they are circularized and used as templates for localized rolling-circle amplification (RCA). The amplification product is finally detected by hybridization of fluorescently labelled complementary oligonucleotides. Created with BioRender.com. (B) Normalized MS signal sum of indicated RBPs in input samples (left panel) and eRIC eluates (third panel from the left) for each organ analyzed. *FDR < 0.05, **FDR<0.01, ***FDR<0.001, ****FDR<0001 (*t* test with FDR correction for multiple testing). In left panel, circles: UV-irradiated samples, triangle: non-irradiated control. Second panel: Total protein intensity assessed by immunofluorescence (IF); right panel: number of interaction sites per cell between the indicated RBP and the oligo(dT) probe (visualized as distinct dots in the PLA). *p.adj < 0.05, **p.adj<0.01, ***p.adj<0.001, ****p.adj<0001 (one-way ANOVA with Tukey *post-hoc* test). Horizontal lines represent arithmetic means.

**Figure S3. Organ-specific differences of RBP activity**

(A-C) Data were analyzed assuming equal mean signal intensity across the RNA-bound proteomes studied. (A) Normalized signal sum of identified RBPs in each of the two crosslinked *ex vivo* eRIC eluates (upper panel) and input samples (bottom panel). Center lines indicate medians, box borders represent the interquartile range (IQR), and whiskers extend to ±1.5 time the IQR; outliers are shown as black dots. (B) Hierarchical clustering and heatmap of the RBPs identified from brain, kidney and liver, showing protein abundance in eRIC (left columns) and input samples (right columns). (C) Pairwise comparison (expressed as log2 ratio) of RBP signal intensities in eRIC eluates (y-axis) and input samples (x-axis) across the three organs analyzed. The names of representative RBPs exhibiting differences in RNA-binding but no corresponding change in overall protein abundance are indicated; when dot density is too high, the corresponding protein names are indicated inside rounded rectangles. Purple dots correspond to RBPs detected in eRIC eluates but giving no signal in input; an artificial input intensity of low magnitude was imputed to these proteins in order to incorporate them in the graphic.

**Figure S4. The RNA-binding proteomes of brain, kidney and liver lack RBPs commonly identified in cultured cells**

(A-D) The eRIC dataset was compared to a combined list of 6518 RBPs from 30 published RBP libraries generated from cultured cells. (A) Number of RBPs (y-axis) identified across an increasing number of studies (x-axis) that were not detected in mouse organs by *ex vivo* eRIC. (B) RBPs not detected by eRIC in organs and detected in at least 50% of previous RBP libraries were classified into two groups based on whether or not they were detected in inputs. The “selected RBPs” were employed for further analysis. (C-D) Protein class distribution (C) and GO term enrichment (D) of the RBPs selected in (B). (E) Input levels of the RBPs in “mRNA metabolic process” that were either identified by *ex vivo* eRIC (left panel) or selected in (B) (left panel and heatmap). Left panel: ***p.adj<0.001, ****p.adj<0001 (one-way ANOVA with Sidák *post-hoc* test). Horizontal lines represent medians. Right panel: Numbers next to RBP names are the number of cell lines studies (total: 30) where the proteins were hits.

**Figure S5. Enzyme-RBPs in energy metabolism**

(A,B) Analysis as in Figure 5B for glycolysis (A) and the tricarboxylic acid cycle (TCA) (B).

**Table S1**. eRIC hits in brain, kidney and liver (related to Figure 1). Also provided: occurrence of organ RBPs in previous RBP profiling studies (related to Figure 4) and identity of RBPs not detected in organs but commonly identified in cultured cells (at least 50% of previous reports) (related to Figure S5).

**Table S2**. Pairwise comparison of eRIC and input samples across organs. eRIC data was normalized without assuming equal mean protein intensity across organs (related to Figure 3).

**Table S3**. Pairwise comparison of eRIC and input samples across organs. eRIC data was normalized assuming same mean protein intensity across organs (related to Figure S3)

**Table S4**. Annotation of enzymes of intermediary metabolism (“metabolite interconversion enzymes” in PANTHER protein class) detected in inputs or eRIC of brain, kidney or liver (related to Figure 6)

**Table S5**. Full results of gene ontology enrichment analysis of eRIC hits in brain, kidney and liver (related to Figure 2B). BP, biological process; MF, molecular function; CC, cellular component.

**Table S6**. Full results of gene ontology enrichment analysis of RBPs identified in cultured cells (>=50% previous reports) and inputs but not in eRIC of organs (related to Figure S4D). BP, biological process; MF, molecular function; CC, cellular component.

**Table S7**. Selected InterPro domains involved in nucleotide binding (related to Figure 6C)

