## Supplementary figures and images for "The RNA-binding protein landscapes differ between mammalian organs and cultured cells"

### Supplementary Figures

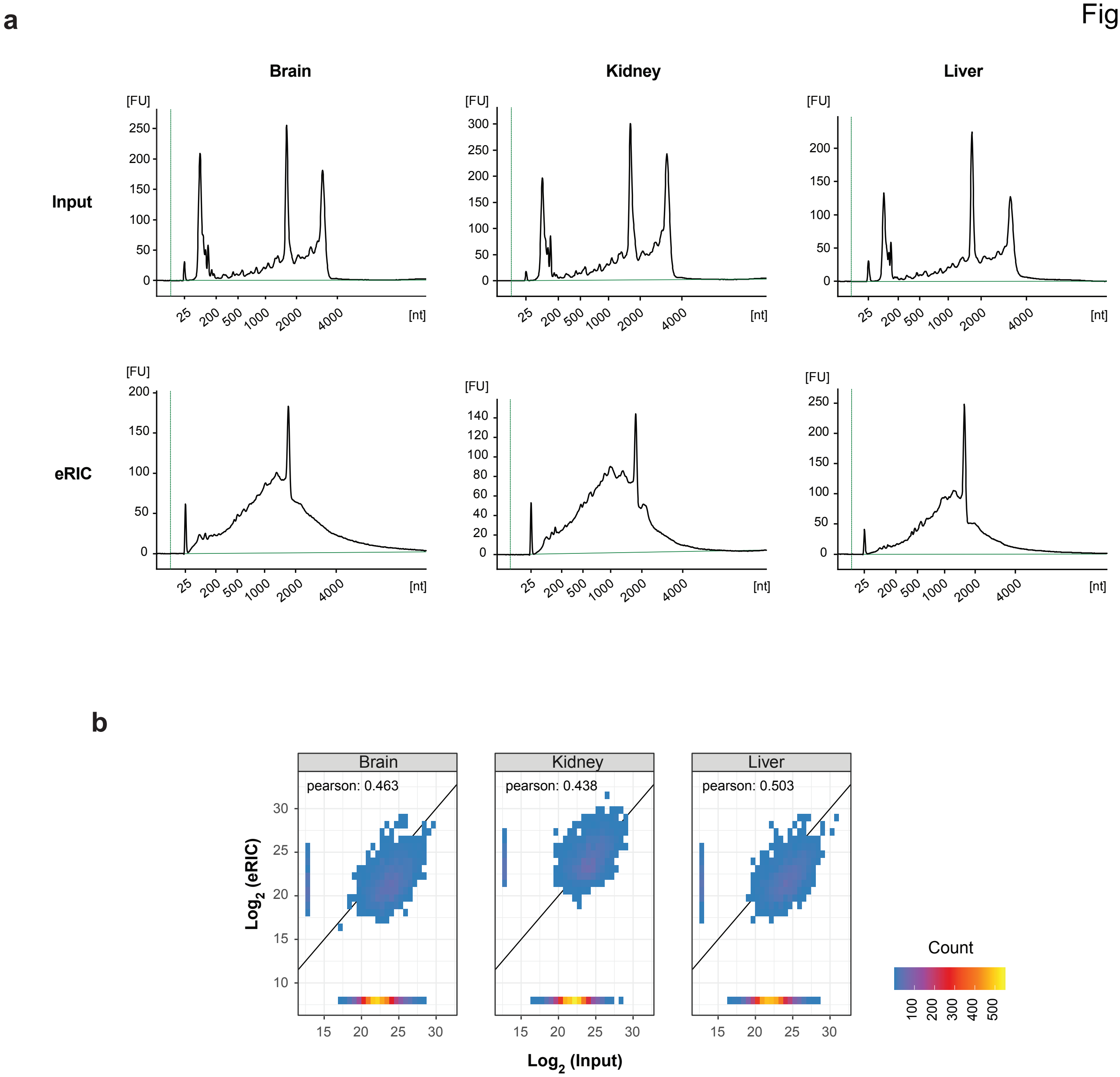

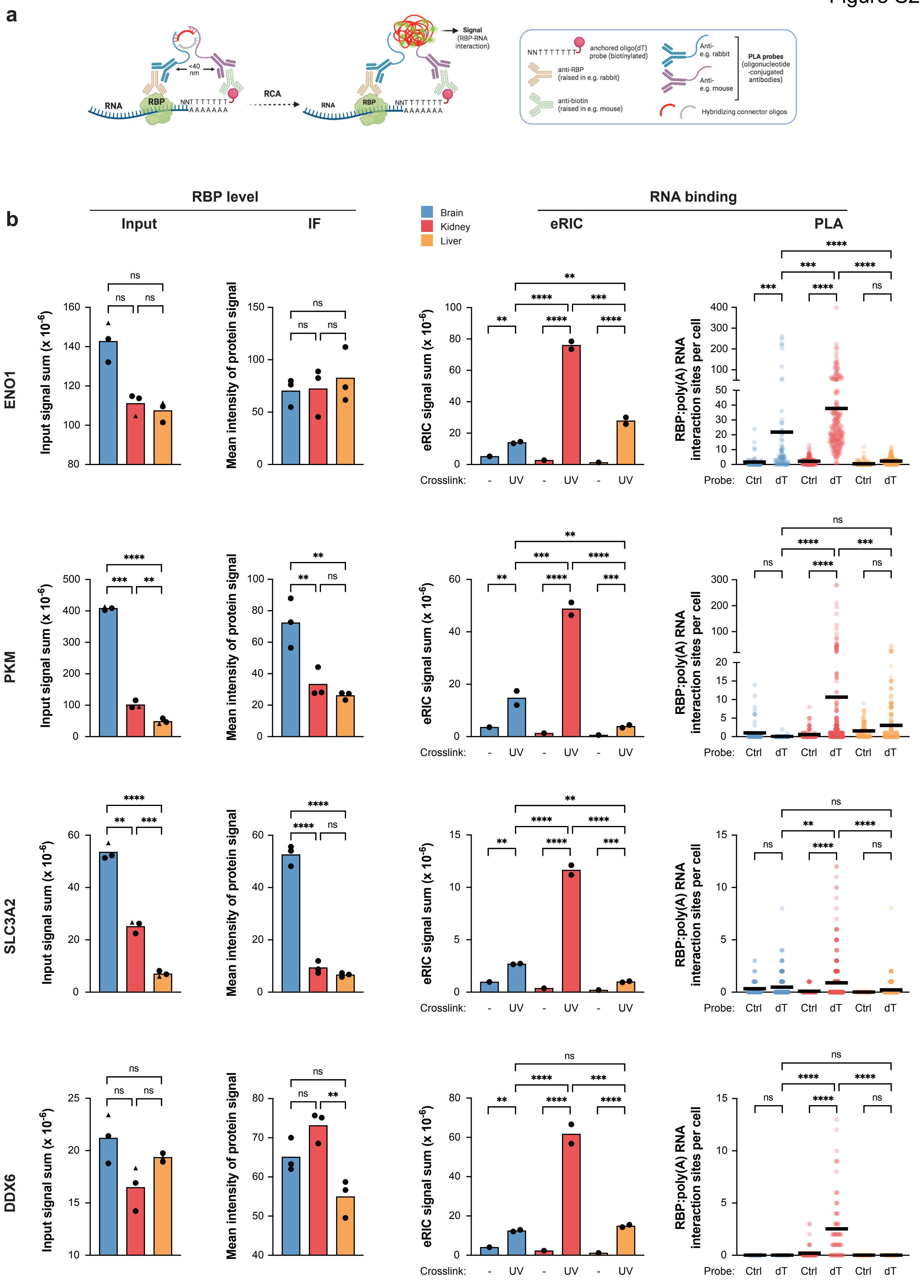

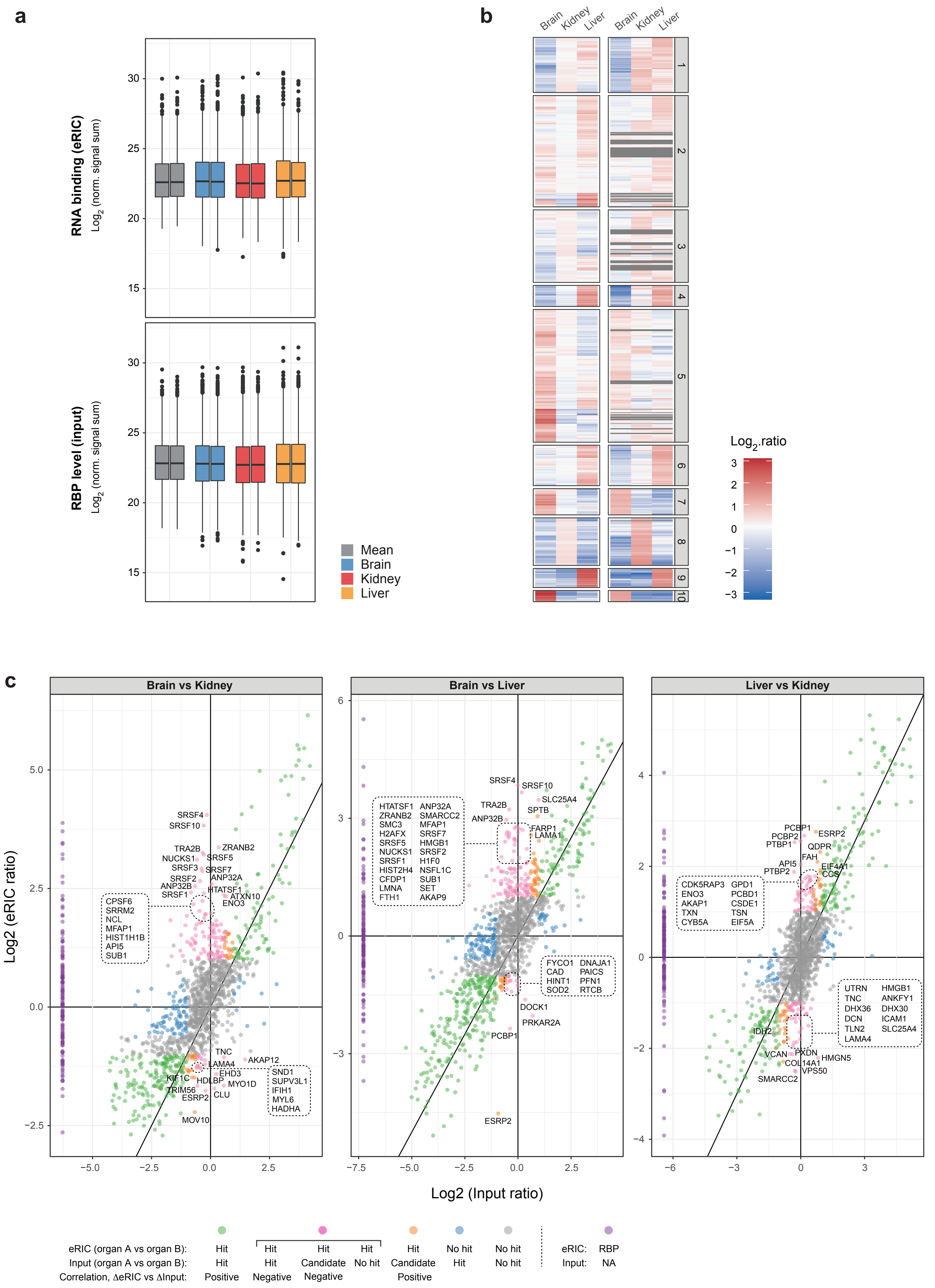

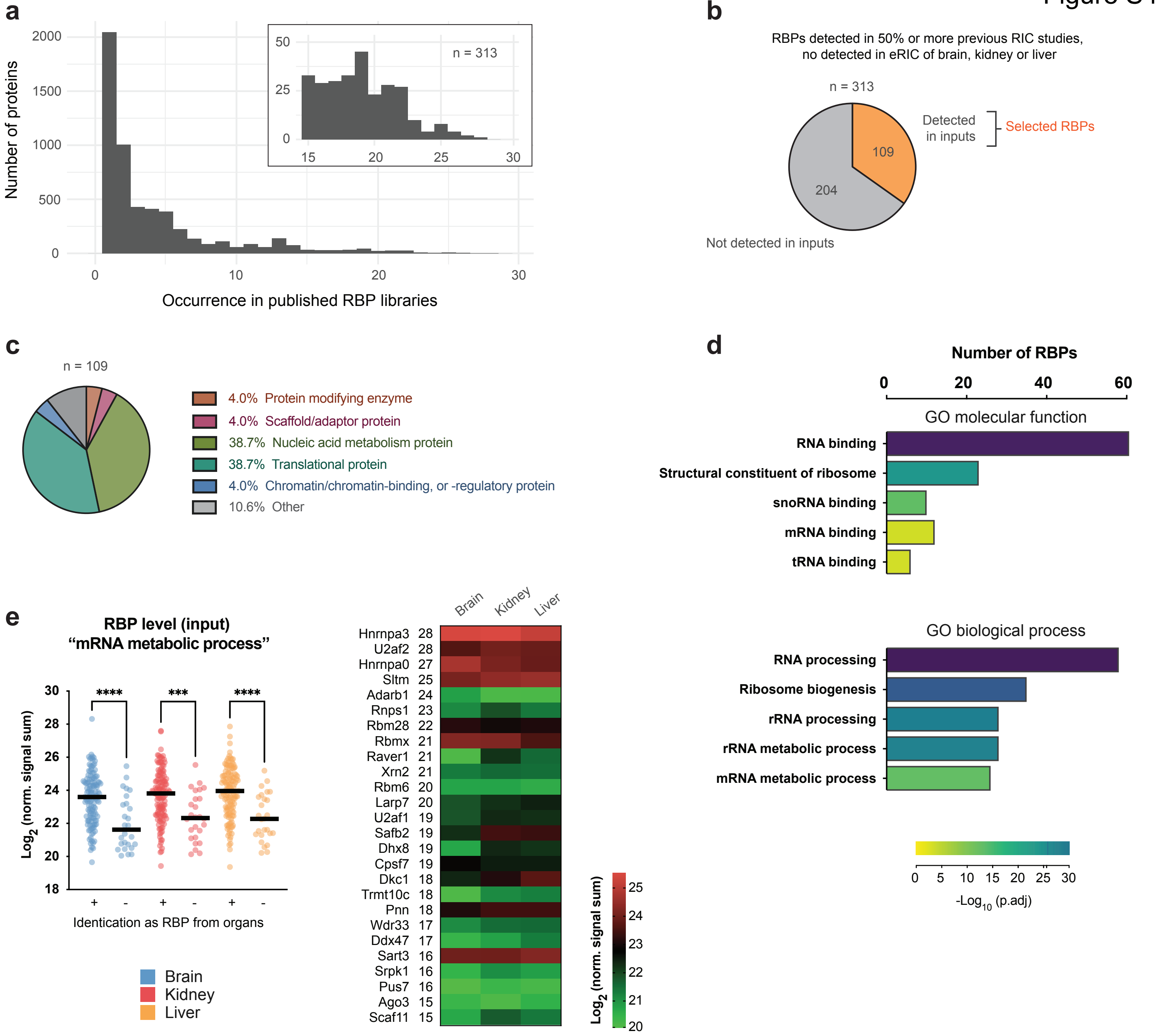

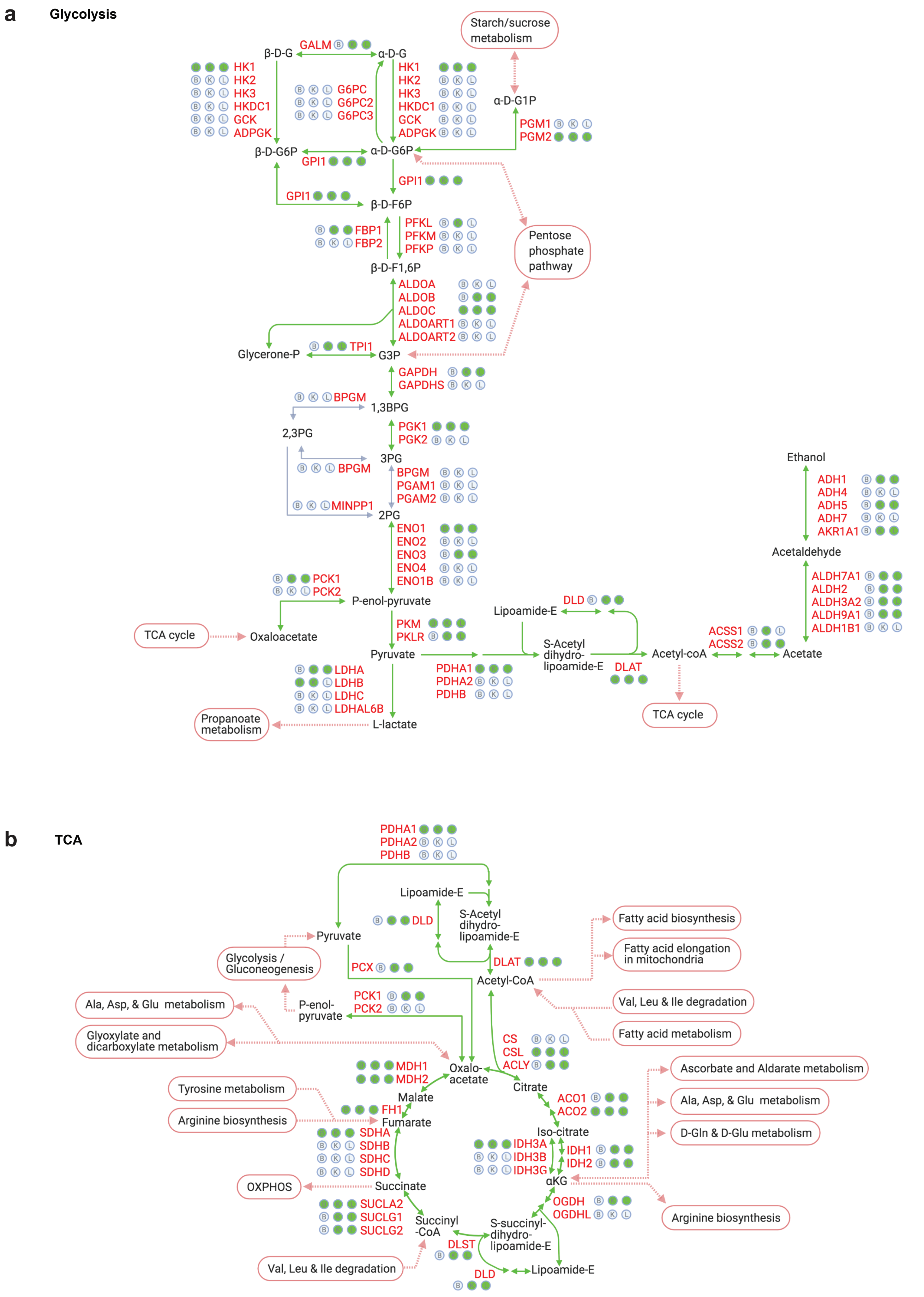
