## Supplementary material for "The RNA-binding protein landscapes differ between mammalian organs and cultured cells": Table S5

**PANTHER 15.0 released!**Analysis Summary: Please report in publication ?**Analysis Type:** PANTHER Overrepresentation Test (Released 20200407)**Annotation Version and Release Date:** GO Ontology database Released 2020-02-21**Analyzed List:** upload\_1 (Mus musculus)Change**Reference List:** Mus musculus (all genes in database)Change**Annotation Data Set:**  ?**Test Type:** ☒ Fisher's Exact ☐ Binomial**Correction:** ☐ Calculate False Discovery Rate ☒ Use the Bonferroni correction for multiple testing ? ☐ No correction**Results** ?

|  | Reference list | upload_1 |
| --- | --- | --- |
| Uniquely Mapped IDs: | 22265 out of 22265 | 622 out of 622 |
| Unmapped IDs: | 0 | 0 |
| Multiple mapping information: | 0 | 0 |

Bonferroni count: 8894

Export Table XML with user input ids JSON with user input ids

Displaying only results for Bonferroni-corrected for P < 0.05, click here to display all results

|  | Mus musculus (REF) | upload_1 (▼ Hierarchy **NEW!** ?) |  |  |  |  |
| --- | --- | --- | --- | --- | --- | --- |
| GO biological process complete | # | # | expected | Fold Enrichment | +/- | P value |
| viral translational termination-reinitiation | 5 | 5 | .14 | 35.80 | + | 2.92E-02 |
| ↳ viral process | 131 | 19 | 3.66 | 5.19 | + | 2.43E-04 |
| ↳ symbiotic process | 228 | 30 | 6.37 | 4.71 | + | 2.09E-07 |
| ↳ viral translation | 14 | 7 | .39 | 17.90 | + | 7.86E-03 |
| ↳ viral gene expression | 20 | 8 | .56 | 14.32 | + | 4.86E-03 |
| positive regulation of mRNA binding | 8 | 6 | .22 | 26.85 | + | 8.72E-03 |
| ↳ regulation of mRNA binding | 10 | 7 | .28 | 25.06 | + | 1.44E-03 |
| ↳ regulation of RNA binding | 12 | 8 | .34 | 23.86 | + | 2.39E-04 |
| ↳ regulation of molecular function | 2559 | 111 | 71.49 | 1.55 | + | 4.04E-02 |
| ↳ biological regulation | 12152 | 436 | 339.48 | 1.28 | + | 6.40E-11 |
| ↳ positive regulation of RNA binding | 9 | 6 | .25 | 23.86 | + | 1.42E-02 |
| positive regulation of establishment of protein localization to telomere | 10 | 7 | .28 | 25.06 | + | 1.44E-03 |
| ↳ positive regulation of establishment of protein localization | 462 | 37 | 12.91 | 2.87 | + | 3.63E-04 |
| ↳ positive regulation of biological process | 6100 | 261 | 170.41 | 1.53 | + | 1.33E-10 |
| ↳ regulation of biological process | 11534 | 414 | 322.22 | 1.28 | + | 2.04E-09 |
| ↳ regulation of establishment of protein localization | 771 | 53 | 21.54 | 2.46 | + | 7.95E-05 |
| ↳ regulation of protein localization | 1075 | 77 | 30.03 | 2.56 | + | 2.39E-09 |
| ↳ regulation of localization | 2886 | 141 | 80.62 | 1.75 | + | 5.38E-07 |
| ↳positive regulation of protein localization to chromosome, telomeric region | 13 | Z | .36 | 19.27 | + | 5.36E-03 |
| ↳regulation of protein localization to chromosome, telomeric region | 15 | Z | .42 | 16.70 | + | 1.13E-02 |
| ↳regulation of cellular protein localization | 560 | 54 | 15.64 | 3.45 | + | 3.01E-10 |
| ↳regulation of cellular localization | 837 | 70 | 23.38 | 2.99 | + | 3.07E-11 |
| ↳positive regulation of cellular protein localization | 308 | 31 | 8.60 | 3.60 | + | 3.93E-05 |
| ↳regulation of establishment of protein localization to telomere | 11 | Z | .31 | 22.78 | + | 2.31E-03 |
| ↳regulation of establishment of protein localization to chromosome | 12 | Z | .34 | 20.88 | + | 3.57E-03 |
| tRNA aminoacylation for protein translation | 40 | 18 | 1.12 | 16.11 | + | 7.47E-11 |
| ↳tRNA aminoacylation | 43 | 18 | 1.20 | 14.98 | + | 2.02E-10 |
| ↳amino acid activation | 44 | 18 | 1.23 | 14.64 | + | 2.77E-10 |
| ↳cellular amino acid metabolic process | 231 | 29 | 6.45 | 4.49 | + | 1.24E-06 |
| ↳primary metabolic process | 6269 | 316 | 175.13 | 1.80 | + | 1.20E-27 |
| ↳metabolic process | 7222 | 351 | 201.76 | 1.74 | + | 1.06E-29 |
| ↳carboxylic acid metabolic process | 761 | 52 | 21.26 | 2.45 | + | 1.27E-04 |
| ↳oxoacid metabolic process | 803 | 52 | 22.43 | 2.32 | + | 6.25E-04 |
| ↳organic acid metabolic process | 829 | 52 | 23.16 | 2.25 | + | 1.82E-03 |
| ↳cellular metabolic process | 6331 | 332 | 176.86 | 1.88 | + | 1.30E-33 |
| ↳cellular process | 14059 | 536 | 392.76 | 1.36 | + | 1.31E-32 |
| ↳small molecule metabolic process | 1432 | 73 | 40.00 | 1.82 | + | 1.34E-02 |
| ↳organic substance metabolic process | 6701 | 332 | 187.20 | 1.77 | + | 1.39E-28 |
| ↳organonitrogen compound metabolic process | 4270 | 202 | 119.29 | 1.69 | + | 9.80E-11 |
| ↳nitrogen compound metabolic process | 5718 | 301 | 159.74 | 1.88 | + | 5.73E-29 |
| ↳tRNA metabolic process | 165 | 20 | 4.61 | 4.34 | + | 1.52E-03 |
| ↳ncRNA metabolic process | 416 | 31 | 11.62 | 2.67 | + | 1.89E-02 |
| ↳RNA metabolic process | 1232 | 126 | 34.42 | 3.66 | + | 4.54E-31 |
| ↳nucleic acid metabolic process | 1767 | 141 | 49.36 | 2.86 | + | 8.48E-25 |
| ↳macromolecule metabolic process | 5056 | 285 | 141.25 | 2.02 | + | 1.19E-31 |
| ↳nucleobase-containing compound metabolic process | 2191 | 165 | 61.21 | 2.70 | + | 2.36E-27 |
| ↳cellular nitrogen compound metabolic process | 2789 | 208 | 77.91 | 2.67 | + | 3.25E-36 |
| ↳organic cyclic compound metabolic process | 2616 | 174 | 73.08 | 2.38 | + | 5.37E-23 |
| ↳heterocycle metabolic process | 2328 | 171 | 65.04 | 2.63 | + | 2.37E-27 |
| ↳cellular aromatic compound metabolic process | 2402 | 169 | 67.10 | 2.52 | + | 7.26E-25 |
| ↳translation | 306 | 61 | 8.55 | 7.14 | + | 2.10E-26 |
| ↳cellular protein metabolic process | 2844 | 161 | 79.45 | 2.03 | + | 4.45E-14 |
| ↳protein metabolic process | 3420 | 171 | 95.54 | 1.79 | + | 2.90E-10 |
| ↳cellular macromolecule metabolic process | 3958 | 206 | 110.57 | 1.86 | + | 1.25E-15 |
| ↳gene expression | 1616 | 158 | 45.14 | 3.50 | + | 2.07E-38 |
| ↳peptide biosynthetic process | 326 | 61 | 9.11 | 6.70 | + | 4.34E-25 |
| ↳organonitrogen compound biosynthetic process | 1070 | 90 | 29.89 | 3.01 | + | 1.45E-15 |
| ↳organic substance biosynthetic process | 2079 | 124 | 58.08 | 2.14 | + | 2.71E-11 |
| ↳biosynthetic process | 2161 | 128 | 60.37 | 2.12 | + | 1.50E-11 |
| ↳peptide metabolic process | 448 | 67 | 12.52 | 5.35 | + | 7.95E-23 |
| ↳cellular amide metabolic process | 665 | 74 | 18.58 | 3.98 | + | 1.85E-18 |
| ↳amide biosynthetic process | 421 | 64 | 11.76 | 5.44 | + | 5.28E-22 |
| ↳cellular nitrogen compound biosynthetic process | 1118 | 93 | 31.23 | 2.98 | + | 6.56E-16 |
| ↳cellular biosynthetic process | 1985 | 122 | 55.45 | 2.20 | + | 6.36E-12 |
| ↳cellular macromolecule biosynthetic process | 1193 | 88 | 33.33 | 2.64 | + | 8.61E-12 |
| macromolecule biosynthetic process | 1227 | 88 | 34.28 | 2.57 | + | 5.55E-11 |
| stress granule assembly | 16 | 7 | .45 | 15.66 | + | 1.58E-02 |
| cellular component assembly | 1941 | 124 | 54.22 | 2.29 | + | 2.42E-13 |
| cellular component organization | 5045 | 245 | 140.94 | 1.74 | + | 3.35E-16 |
| cellular component organization or biogenesis | 5239 | 250 | 146.36 | 1.71 | + | 1.04E-15 |
| cellular component biogenesis | 2168 | 130 | 60.57 | 2.15 | + | 2.99E-12 |
| organelle organization | 3054 | 174 | 85.32 | 2.04 | + | 6.32E-16 |
| regulation of alternative mRNA splicing, via spliceosome | 66 | 26 | 1.84 | 14.10 | + | 1.09E-15 |
| regulation of mRNA splicing, via spliceosome | 110 | 37 | 3.07 | 12.04 | + | 1.75E-21 |
| regulation of RNA splicing | 145 | 43 | 4.05 | 10.62 | + | 1.42E-23 |
| regulation of gene expression | 3827 | 191 | 106.91 | 1.79 | + | 4.06E-12 |
| regulation of macromolecule metabolic process | 5484 | 255 | 153.20 | 1.66 | + | 9.63E-15 |
| regulation of metabolic process | 5976 | 270 | 166.95 | 1.62 | + | 2.31E-14 |
| regulation of RNA metabolic process | 3165 | 147 | 88.42 | 1.66 | + | 6.05E-06 |
| regulation of nucleobase-containing compound metabolic process | 3401 | 161 | 95.01 | 1.69 | + | 1.49E-07 |
| regulation of primary metabolic process | 5349 | 244 | 149.43 | 1.63 | + | 1.17E-12 |
| regulation of nitrogen compound metabolic process | 5198 | 242 | 145.21 | 1.67 | + | 1.59E-13 |
| regulation of cellular metabolic process | 5552 | 254 | 155.10 | 1.64 | + | 1.07E-13 |
| regulation of cellular process | 10826 | 394 | 302.44 | 1.30 | + | 3.86E-09 |
| regulation of mRNA processing | 149 | 43 | 4.16 | 10.33 | + | 3.57E-23 |
| regulation of mRNA metabolic process | 255 | 65 | 7.12 | 9.12 | + | 7.64E-34 |
| alternative mRNA splicing, via spliceosome | 18 | 7 | .50 | 13.92 | + | 2.96E-02 |
| mRNA splicing, via spliceosome | 196 | 42 | 5.48 | 7.67 | + | 2.76E-18 |
| mRNA processing | 405 | 73 | 11.31 | 6.45 | + | 9.02E-30 |
| RNA processing | 752 | 87 | 21.01 | 4.14 | + | 2.03E-23 |
| mRNA metabolic process | 531 | 85 | 14.83 | 5.73 | + | 6.98E-32 |
| RNA splicing, via transesterification reactions with bulged adenosine as nucleophile | 196 | 42 | 5.48 | 7.67 | + | 2.76E-18 |
| RNA splicing, via transesterification reactions | 196 | 42 | 5.48 | 7.67 | + | 2.76E-18 |
| RNA splicing | 318 | 68 | 8.88 | 7.65 | + | 1.85E-31 |
| 3'-UTR-mediated mRNA destabilization | 18 | 7 | .50 | 13.92 | + | 2.96E-02 |
| mRNA destabilization | 32 | 13 | .89 | 14.54 | + | 1.14E-06 |
| RNA destabilization | 35 | 13 | .98 | 13.30 | + | 2.79E-06 |
| regulation of RNA stability | 109 | 27 | 3.05 | 8.87 | + | 3.92E-12 |
| regulation of cellular catabolic process | 698 | 65 | 19.50 | 3.33 | + | 2.91E-12 |
| regulation of catabolic process | 843 | 73 | 23.55 | 3.10 | + | 1.36E-12 |
| posttranscriptional regulation of gene expression | 427 | 81 | 11.93 | 6.79 | + | 7.28E-35 |
| regulation of biological quality | 3959 | 213 | 110.60 | 1.93 | + | 3.97E-18 |
| positive regulation of nucleobase-containing compound metabolic process | 1811 | 86 | 50.59 | 1.70 | + | 2.48E-02 |
| positive regulation of cellular metabolic process | 3270 | 142 | 91.35 | 1.55 | + | 1.11E-03 |
| positive regulation of cellular process | 5424 | 235 | 151.53 | 1.55 | + | 2.28E-09 |
| positive regulation of metabolic process | 3560 | 155 | 99.45 | 1.56 | + | 1.84E-04 |
| positive regulation of nitrogen compound metabolic process | 3086 | 139 | 86.21 | 1.61 | + | 1.47E-04 |
| positive regulation of macromolecule metabolic process | 3256 | 141 | 90.96 | 1.55 | + | 1.45E-03 |
| positive regulation of cellular catabolic process | 367 | 35 | 10.25 | 3.41 | + | 1.45E-05 |
| positive regulation of catabolic process | 436 | 40 | 12.18 | 3.28 | + | 2.79E-06 |
| positive regulation of mRNA catabolic process | 50 | 15 | 1.40 | 10.74 | + | 1.45E-06 |
| positive regulation of mRNA metabolic process | 93 | 24 | 2.60 | 9.24 | + | 7.87E-11 |
| ↪regulation of mRNA catabolic process | 121 | 29 | 3.38 | 8.58 | + | 6.72E-13 |
| ↪regulation of mRNA stability | 98 | 27 | 2.74 | 9.86 | + | 4.09E-13 |
| ↪negative regulation of translation | 130 | 34 | 3.63 | 9.36 | + | 1.34E-16 |
| ↪negative regulation of cellular protein metabolic process | 1004 | 71 | 28.05 | 2.53 | + | 5.74E-08 |
| ↪negative regulation of cellular metabolic process | 2469 | 129 | 68.97 | 1.87 | + | 7.94E-08 |
| ↪negative regulation of metabolic process | 2766 | 145 | 77.27 | 1.88 | + | 1.59E-09 |
| ↪negative regulation of biological process | 5169 | 247 | 144.40 | 1.71 | + | 1.74E-15 |
| ↪negative regulation of cellular process | 4632 | 217 | 129.40 | 1.68 | + | 1.15E-11 |
| ↪regulation of cellular protein metabolic process | 2531 | 137 | 70.71 | 1.94 | + | 1.19E-09 |
| ↪regulation of protein metabolic process | 2723 | 147 | 76.07 | 1.93 | + | 9.65E-11 |
| ↪negative regulation of protein metabolic process | 1072 | 72 | 29.95 | 2.40 | + | 2.98E-07 |
| ↪negative regulation of nitrogen compound metabolic process | 2272 | 118 | 63.47 | 1.86 | + | 9.41E-07 |
| ↪negative regulation of macromolecule metabolic process | 2494 | 132 | 69.67 | 1.89 | + | 1.55E-08 |
| ↪negative regulation of cellular amide metabolic process | 149 | 35 | 4.16 | 8.41 | + | 7.05E-16 |
| ↪regulation of cellular amide metabolic process | 384 | 71 | 10.73 | 6.62 | + | 1.80E-29 |
| ↪regulation of translation | 333 | 70 | 9.30 | 7.52 | + | 4.46E-32 |
| ↪regulation of cellular macromolecule biosynthetic process | 3333 | 156 | 93.11 | 1.68 | + | 9.83E-07 |
| ↪regulation of macromolecule biosynthetic process | 3419 | 159 | 95.51 | 1.66 | + | 1.05E-06 |
| ↪regulation of biosynthetic process | 3644 | 167 | 101.80 | 1.64 | + | 7.38E-07 |
| ↪regulation of cellular biosynthetic process | 3571 | 164 | 99.76 | 1.64 | + | 1.07E-06 |
| ↪negative regulation of macromolecule biosynthetic process | 1414 | 71 | 39.50 | 1.80 | + | 2.83E-02 |
| ↪negative regulation of gene expression | 1643 | 93 | 45.90 | 2.03 | + | 2.28E-06 |
| positive regulation of gene silencing by miRNA | 24 | 9 | .67 | 13.42 | + | 1.46E-03 |
| ↪positive regulation of posttranscriptional gene silencing | 25 | 9 | .70 | 12.89 | + | 1.94E-03 |
| ↪regulation of posttranscriptional gene silencing | 44 | 10 | 1.23 | 8.14 | + | 1.48E-02 |
| ↪regulation of gene silencing by miRNA | 41 | 10 | 1.15 | 8.73 | + | 8.51E-03 |
| ↪regulation of gene silencing by RNA | 44 | 10 | 1.23 | 8.14 | + | 1.48E-02 |
| negative regulation of mRNA splicing, via spliceosome | 24 | 9 | .67 | 13.42 | + | 1.46E-03 |
| ↪negative regulation of RNA splicing | 29 | 10 | .81 | 12.34 | + | 5.68E-04 |
| ↪negative regulation of mRNA processing | 32 | 10 | .89 | 11.19 | + | 1.22E-03 |
| ↪negative regulation of mRNA metabolic process | 82 | 21 | 2.29 | 9.17 | + | 4.26E-09 |
| regulation of cytoplasmic translation | 25 | 9 | .70 | 12.89 | + | 1.94E-03 |
| mRNA cis splicing, via spliceosome | 28 | 10 | .78 | 12.78 | + | 4.33E-04 |
| tricarboxylic acid cycle | 32 | 10 | .89 | 11.19 | + | 1.22E-03 |
| ↪generation of precursor metabolites and energy | 286 | 29 | 7.99 | 3.63 | + | 1.07E-04 |
| translational initiation | 55 | 17 | 1.54 | 11.06 | + | 5.83E-08 |
| nuclear migration | 31 | 9 | .87 | 10.39 | + | 8.74E-03 |
| ↪nucleus localization | 36 | 9 | 1.01 | 8.95 | + | 2.51E-02 |
| ↪organelle localization | 474 | 46 | 13.24 | 3.47 | + | 2.12E-08 |
| ↪cellular localization | 2053 | 166 | 57.35 | 2.89 | + | 4.85E-31 |
| ↪localization | 4864 | 233 | 135.88 | 1.71 | + | 4.04E-14 |
| ↪intracellular transport | 1182 | 116 | 33.02 | 3.51 | + | 1.11E-26 |
| ↪transport | 3572 | 190 | 99.79 | 1.90 | + | 6.94E-15 |
| ↪establishment of localization | 3711 | 197 | 103.67 | 1.90 | + | 1.73E-15 |
| ↪establishment of organelle localization | 331 | 34 | 9.25 | 3.68 | + | 4.30E-06 |
| positive regulation of telomere maintenance via telomere lengthening | 35 | 10 | .98 | 10.23 | + | 2.46E-03 |
| ↪positive regulation of telomere maintenance | 50 | 11 | 1.40 | 7.88 | + | 5.92E-03 |
| ↳positive regulation of DNA metabolic process | 212 | 21 | 5.92 | 3.55 | + | 1.55E-02 |
| ↳regulation of DNA metabolic process | 368 | 28 | 10.28 | 2.72 | + | 4.07E-02 |
| ↳positive regulation of organelle organization | 596 | 50 | 16.65 | 3.00 | + | 3.10E-07 |
| ↳positive regulation of cellular component organization | 1254 | 86 | 35.03 | 2.45 | + | 9.87E-10 |
| ↳regulation of cellular component organization | 2537 | 157 | 70.87 | 2.22 | + | 5.18E-17 |
| ↳regulation of organelle organization | 1270 | 95 | 35.48 | 2.68 | + | 1.81E-13 |
| ↳regulation of telomere maintenance | 81 | 15 | 2.26 | 6.63 | + | 4.20E-04 |
| positive regulation of dendritic spine morphogenesis | 29 | 8 | .81 | 9.87 | + | 4.87E-02 |
| ↳positive regulation of dendritic spine development | 65 | 14 | 1.82 | 7.71 | + | 2.26E-04 |
| ↳regulation of dendritic spine development | 97 | 16 | 2.71 | 5.90 | + | 6.28E-04 |
| ↳regulation of dendrite development | 194 | 21 | 5.42 | 3.87 | + | 4.18E-03 |
| ↳regulation of neuron projection development | 609 | 43 | 17.01 | 2.53 | + | 9.43E-04 |
| ↳regulation of plasma membrane bounded cell projection organization | 781 | 52 | 21.82 | 2.38 | + | 2.38E-04 |
| ↳regulation of cell projection organization | 790 | 52 | 22.07 | 2.36 | + | 4.70E-04 |
| ↳regulation of neuron differentiation | 776 | 48 | 21.68 | 2.21 | + | 7.99E-03 |
| ↳regulation of neurogenesis | 953 | 58 | 26.62 | 2.18 | + | 7.84E-04 |
| ↳generation of neurons | 1663 | 90 | 46.46 | 1.94 | + | 3.63E-05 |
| ↳neurogenesis | 1771 | 93 | 49.48 | 1.88 | + | 9.34E-05 |
| ↳nervous system development | 2266 | 114 | 63.30 | 1.80 | + | 1.27E-05 |
| ↳regulation of cell development | 1092 | 66 | 30.51 | 2.16 | + | 1.21E-04 |
| ↳regulation of cell differentiation | 1870 | 88 | 52.24 | 1.68 | + | 2.44E-02 |
| ↳regulation of developmental process | 2684 | 118 | 74.98 | 1.57 | + | 9.98E-03 |
| ↳regulation of nervous system development | 1070 | 61 | 29.89 | 2.04 | + | 3.77E-03 |
| ↳regulation of multicellular organismal process | 3223 | 143 | 90.04 | 1.59 | + | 2.54E-04 |
| ↳positive regulation of dendrite development | 110 | 16 | 3.07 | 5.21 | + | 2.92E-03 |
| ↳positive regulation of neuron projection development | 369 | 30 | 10.31 | 2.91 | + | 5.25E-03 |
| ↳positive regulation of cell projection organization | 474 | 36 | 13.24 | 2.72 | + | 1.87E-03 |
| ↳positive regulation of neuron differentiation | 469 | 33 | 13.10 | 2.52 | + | 3.59E-02 |
| ↳positive regulation of neurogenesis | 583 | 38 | 16.29 | 2.33 | + | 3.34E-02 |
| ↳positive regulation of cell development | 665 | 42 | 18.58 | 2.26 | + | 2.36E-02 |
| ↳positive regulation of cell morphogenesis involved in differentiation | 189 | 22 | 5.28 | 4.17 | + | 7.08E-04 |
| ↳regulation of cell morphogenesis involved in differentiation | 348 | 34 | 9.72 | 3.50 | + | 1.41E-05 |
| ↳regulation of cell morphogenesis | 537 | 47 | 15.00 | 3.13 | + | 3.29E-07 |
| ↳regulation of anatomical structure morphogenesis | 1090 | 62 | 30.45 | 2.04 | + | 2.93E-03 |
| ↳regulation of dendritic spine morphogenesis | 56 | 11 | 1.56 | 7.03 | + | 1.57E-02 |
| ↳regulation of postsynapse organization | 131 | 16 | 3.66 | 4.37 | + | 2.35E-02 |
| ↳regulation of synapse organization | 273 | 28 | 7.63 | 3.67 | + | 1.53E-04 |
| ↳regulation of synapse structure or activity | 285 | 30 | 7.96 | 3.77 | + | 2.72E-05 |
| positive regulation of mRNA splicing, via spliceosome | 33 | 9 | .92 | 9.76 | + | 1.36E-02 |
| ↳positive regulation of mRNA processing | 44 | 11 | 1.23 | 8.95 | + | 1.97E-03 |
| ↳positive regulation of gene expression | 1995 | 97 | 55.73 | 1.74 | + | 1.77E-03 |
| ↳positive regulation of RNA splicing | 50 | 15 | 1.40 | 10.74 | + | 1.45E-06 |
| positive regulation of translation | 123 | 33 | 3.44 | 9.60 | + | 2.56E-16 |
| ↳positive regulation of biosynthetic process | 1953 | 92 | 54.56 | 1.69 | + | 1.32E-02 |
| ↳positive regulation of cellular amide metabolic process | 149 | 34 | 4.16 | 8.17 | + | 5.19E-15 |
| ↳positive regulation of cellular biosynthetic process | 1912 | 91 | 53.41 | 1.70 | + | 1.03E-02 |
| spliceosomal complex assembly | 45 | 12 | 1.26 | 9.55 | + | 3.01E-04 |
| ↳ribonucleoprotein complex assembly | 163 | 25 | 4.55 | 5.49 | + | 5.44E-07 |
| ↳cellular protein-containing complex assembly | 653 | 47 | 18.24 | 2.58 | + | 1.47E-04 |
| ↳protein-containing complex assembly | 1003 | 67 | 28.02 | 2.39 | + | 1.97E-06 |
| ↳protein-containing complex subunit organization | 1144 | 77 | 31.96 | 2.41 | + | 4.86E-08 |
| ↳ribonucleoprotein complex biogenesis | 392 | 33 | 10.95 | 3.01 | + | 6.78E-04 |
| ↳ribonucleoprotein complex subunit organization | 170 | 26 | 4.75 | 5.47 | + | 2.38E-07 |
| regulation of translational initiation | 59 | 15 | 1.65 | 9.10 | + | 1.02E-05 |
| mRNA stabilization | 41 | 10 | 1.15 | 8.73 | + | 8.51E-03 |
| ↳negative regulation of mRNA catabolic process | 51 | 11 | 1.42 | 7.72 | + | 7.03E-03 |
| ↳negative regulation of RNA catabolic process | 61 | 13 | 1.70 | 7.63 | + | 7.92E-04 |
| ↳negative regulation of cellular catabolic process | 241 | 27 | 6.73 | 4.01 | + | 5.05E-05 |
| ↳negative regulation of catabolic process | 301 | 28 | 8.41 | 3.33 | + | 1.01E-03 |
| ↳RNA stabilization | 48 | 11 | 1.34 | 8.20 | + | 4.17E-03 |
| posttranscriptional gene silencing by RNA | 47 | 11 | 1.31 | 8.38 | + | 3.47E-03 |
| ↳gene silencing by RNA | 77 | 12 | 2.15 | 5.58 | + | 4.57E-02 |
| ↳posttranscriptional gene silencing | 48 | 11 | 1.34 | 8.20 | + | 4.17E-03 |
| Golgi to plasma membrane transport | 52 | 11 | 1.45 | 7.57 | + | 8.31E-03 |
| ↳vesicle-mediated transport to the plasma membrane | 85 | 15 | 2.37 | 6.32 | + | 7.35E-04 |
| ↳vesicle-mediated transport | 1271 | 81 | 35.51 | 2.28 | + | 1.72E-07 |
| ↳post-Golgi vesicle-mediated transport | 88 | 15 | 2.46 | 6.10 | + | 1.10E-03 |
| ↳Golgi vesicle transport | 250 | 35 | 6.98 | 5.01 | + | 8.73E-10 |
| cortical cytoskeleton organization | 59 | 12 | 1.65 | 7.28 | + | 3.85E-03 |
| ↳cytoskeleton organization | 1070 | 75 | 29.89 | 2.51 | + | 1.60E-08 |
| regulation of actin filament depolymerization | 51 | 10 | 1.42 | 7.02 | + | 4.71E-02 |
| ↳regulation of actin polymerization or depolymerization | 185 | 22 | 5.17 | 4.26 | + | 5.03E-04 |
| ↳regulation of actin filament-based process | 409 | 31 | 11.43 | 2.71 | + | 1.36E-02 |
| ↳regulation of cytoskeleton organization | 559 | 51 | 15.62 | 3.27 | + | 1.12E-08 |
| ↳regulation of supramolecular fiber organization | 368 | 34 | 10.28 | 3.31 | + | 5.19E-05 |
| ↳regulation of actin filament length | 188 | 22 | 5.25 | 4.19 | + | 6.50E-04 |
| ↳regulation of cellular component size | 407 | 39 | 11.37 | 3.43 | + | 1.46E-06 |
| ↳regulation of anatomical structure size | 571 | 47 | 15.95 | 2.95 | + | 2.21E-06 |
| ↳regulation of protein depolymerization | 89 | 13 | 2.49 | 5.23 | + | 3.53E-02 |
| ↳regulation of protein-containing complex disassembly | 121 | 16 | 3.38 | 4.73 | + | 9.18E-03 |
| positive regulation of axon extension | 57 | 11 | 1.59 | 6.91 | + | 1.83E-02 |
| ↳positive regulation of axonogenesis | 105 | 14 | 2.93 | 4.77 | + | 3.95E-02 |
| ↳regulation of axonogenesis | 207 | 20 | 5.78 | 3.46 | + | 3.86E-02 |
| ↳positive regulation of developmental growth | 216 | 21 | 6.03 | 3.48 | + | 2.04E-02 |
| ↳positive regulation of growth | 299 | 26 | 8.35 | 3.11 | + | 9.35E-03 |
| ↳positive regulation of cell growth | 187 | 21 | 5.22 | 4.02 | + | 2.40E-03 |
| ↳regulation of axon extension | 113 | 15 | 3.16 | 4.75 | + | 1.90E-02 |
| negative regulation of protein polymerization | 73 | 14 | 2.04 | 6.86 | + | 8.03E-04 |
| ↳negative regulation of protein-containing complex assembly | 136 | 18 | 3.80 | 4.74 | + | 1.94E-03 |
| ↳regulation of protein-containing complex assembly | 434 | 43 | 12.12 | 3.55 | + | 6.07E-08 |
| ↳regulation of cellular component biogenesis | 964 | 70 | 26.93 | 2.60 | + | 1.95E-08 |
| ↳negative regulation of cellular component organization | 734 | 47 | 20.51 | 2.29 | + | 3.87E-03 |
| ↳regulation of protein polymerization | 224 | 31 | 6.26 | 4.95 | + | 3.01E-08 |
| axo-dendritic transport | 73 | 13 | 2.04 | 6.37 | + | 4.89E-03 |
| ↳transport along microtubule | 155 | 19 | 4.33 | 4.39 | + | 2.63E-03 |
| ↳microtubule-based transport | 184 | 19 | 5.14 | 3.70 | + | 2.76E-02 |
| ↳microtubule-based movement | 267 | 24 | 7.46 | 3.22 | + | 1.42E-02 |
| ↳microtubule-based process | 666 | 45 | 18.61 | 2.42 | + | 1.80E-03 |
| ↳cytoskeleton-dependent intracellular transport | 186 | 20 | 5.20 | 3.85 | + | 8.59E-03 |
| mRNA transport | 107 | 18 | 2.99 | 6.02 | + | 7.32E-05 |
| ↳RNA transport | 150 | 21 | 4.19 | 5.01 | + | 7.92E-05 |
| ↳nucleic acid transport | 150 | 21 | 4.19 | 5.01 | + | 7.92E-05 |
| ↳nucleobase-containing compound transport | 183 | 22 | 5.11 | 4.30 | + | 4.22E-04 |
| ↳organic substance transport | 1856 | 118 | 51.85 | 2.28 | + | 1.61E-12 |
| ↳nitrogen compound transport | 1550 | 112 | 43.30 | 2.59 | + | 2.62E-15 |
| ↳establishment of RNA localization | 152 | 21 | 4.25 | 4.95 | + | 9.76E-05 |
| ↳RNA localization | 170 | 24 | 4.75 | 5.05 | + | 5.93E-06 |
| ↳macromolecule localization | 2266 | 153 | 63.30 | 2.42 | + | 3.83E-20 |
| endoplasmic reticulum to Golgi vesicle-mediated transport | 105 | 17 | 2.93 | 5.80 | + | 3.13E-04 |
| positive regulation of viral process | 82 | 13 | 2.29 | 5.67 | + | 1.57E-02 |
| positive regulation of DNA biosynthetic process | 76 | 12 | 2.12 | 5.65 | + | 4.06E-02 |
| protein folding | 146 | 22 | 4.08 | 5.39 | + | 1.03E-05 |
| protein stabilization | 174 | 26 | 4.86 | 5.35 | + | 3.77E-07 |
| ↳regulation of protein stability | 278 | 30 | 7.77 | 3.86 | + | 1.60E-05 |
| lysosomal transport | 94 | 14 | 2.63 | 5.33 | + | 1.23E-02 |
| ↳vacuolar transport | 131 | 16 | 3.66 | 4.37 | + | 2.35E-02 |
| Golgi organization | 111 | 16 | 3.10 | 5.16 | + | 3.26E-03 |
| ↳endomembrane system organization | 367 | 41 | 10.25 | 4.00 | + | 5.82E-09 |
| protein import into nucleus | 98 | 14 | 2.74 | 5.11 | + | 1.91E-02 |
| ↳import into nucleus | 102 | 14 | 2.85 | 4.91 | + | 2.92E-02 |
| ↳nucleocytoplasmic transport | 203 | 21 | 5.67 | 3.70 | + | 8.21E-03 |
| ↳nuclear transport | 203 | 21 | 5.67 | 3.70 | + | 8.21E-03 |
| ↳establishment of protein localization to organelle | 302 | 38 | 8.44 | 4.50 | + | 1.54E-09 |
| ↳protein localization to organelle | 622 | 65 | 17.38 | 3.74 | + | 1.53E-14 |
| ↳cellular protein localization | 1423 | 114 | 39.75 | 2.87 | + | 3.42E-19 |
| ↳cellular macromolecule localization | 1430 | 114 | 39.95 | 2.85 | + | 4.96E-19 |
| ↳protein localization | 1963 | 136 | 54.84 | 2.48 | + | 4.64E-18 |
| ↳establishment of protein localization | 1330 | 105 | 37.16 | 2.83 | + | 6.81E-17 |
| ↳protein transport | 1246 | 98 | 34.81 | 2.82 | + | 2.15E-15 |
| ↳peptide transport | 1276 | 98 | 35.65 | 2.75 | + | 9.78E-15 |
| ↳amide transport | 1301 | 99 | 36.35 | 2.72 | + | 1.14E-14 |
| ↳protein localization to nucleus | 172 | 19 | 4.81 | 3.95 | + | 1.11E-02 |
| ↳intracellular protein transport | 765 | 70 | 21.37 | 3.28 | + | 4.42E-13 |
| maintenance of location in cell | 103 | 14 | 2.88 | 4.87 | + | 3.23E-02 |
| cytosolic transport | 132 | 17 | 3.69 | 4.61 | + | 5.92E-03 |
| regulation of microtubule cytoskeleton organization | 203 | 26 | 5.67 | 4.58 | + | 7.62E-06 |
| ↳regulation of microtubule-based process | 241 | 28 | 6.73 | 4.16 | + | 1.26E-05 |
| regulation of nucleocytoplasmic transport | 125 | 16 | 3.49 | 4.58 | + | 1.35E-02 |
| ↳regulation of intracellular transport | 352 | 33 | 9.83 | 3.36 | + | 6.28E-05 |
| ↳regulation of transport | 1972 | 100 | 55.09 | 1.82 | + | 1.33E-04 |
| regulation of actin filament polymerization | 168 | 21 | 4.69 | 4.47 | + | 4.66E-04 |
| negative regulation of supramolecular fiber organization | 154 | 19 | 4.30 | 4.42 | + | 2.41E-03 |
| establishment of vesicle localization | 142 | 17 | 3.97 | 4.29 | + | 1.48E-02 |
| ↳ establishment of localization in cell | 353 | 29 | 9.86 | 2.94 | + | 6.64E-03 |
| ↳ vesicle localization | 154 | 18 | 4.30 | 4.18 | + | 1.01E-02 |
| negative regulation of cytoskeleton organization | 161 | 19 | 4.50 | 4.22 | + | 4.47E-03 |
| ATP metabolic process | 170 | 19 | 4.75 | 4.00 | + | 9.44E-03 |
| establishment of protein localization to membrane | 180 | 19 | 5.03 | 3.78 | + | 2.05E-02 |
| endosomal transport | 203 | 20 | 5.67 | 3.53 | + | 2.94E-02 |
| vesicle organization | 234 | 23 | 6.54 | 3.52 | + | 5.71E-03 |
| protein targeting | 217 | 21 | 6.06 | 3.46 | + | 2.18E-02 |
| establishment or maintenance of cell polarity | 207 | 20 | 5.78 | 3.46 | + | 3.86E-02 |
| positive regulation of protein-containing complex assembly | 221 | 21 | 6.17 | 3.40 | + | 2.84E-02 |
| negative regulation of neuron death | 247 | 23 | 6.90 | 3.33 | + | 1.34E-02 |
| ↳ negative regulation of cell death | 1041 | 59 | 29.08 | 2.03 | + | 7.31E-03 |
| ↳ regulation of cell death | 1672 | 84 | 46.71 | 1.80 | + | 3.17E-03 |
| regulation of muscle system process | 242 | 22 | 6.76 | 3.25 | + | 3.24E-02 |
| actin cytoskeleton organization | 500 | 38 | 13.97 | 2.72 | + | 8.43E-04 |
| ↳ actin filament-based process | 555 | 41 | 15.50 | 2.64 | + | 5.37E-04 |
| cell part morphogenesis | 507 | 37 | 14.16 | 2.61 | + | 3.17E-03 |
| cellular response to nitrogen compound | 512 | 35 | 14.30 | 2.45 | + | 2.84E-02 |
| ↳ response to nitrogen compound | 851 | 55 | 23.77 | 2.31 | + | 2.51E-04 |
| ↳ response to chemical | 3484 | 144 | 97.33 | 1.48 | + | 1.35E-02 |
| ↳ cellular response to chemical stimulus | 2359 | 106 | 65.90 | 1.61 | + | 1.76E-02 |
| neuron projection development | 691 | 47 | 19.30 | 2.43 | + | 8.03E-04 |
| ↳ neuron development | 843 | 49 | 23.55 | 2.08 | + | 3.73E-02 |
| response to organonitrogen compound | 745 | 49 | 20.81 | 2.35 | + | 8.58E-04 |
| ↳ response to organic substance | 2488 | 115 | 69.51 | 1.65 | + | 1.09E-03 |
| cellular macromolecule catabolic process | 762 | 48 | 21.29 | 2.25 | + | 3.99E-03 |
| ↳ catabolic process | 1730 | 84 | 48.33 | 1.74 | + | 1.51E-02 |
| ↳ macromolecule catabolic process | 858 | 51 | 23.97 | 2.13 | + | 9.68E-03 |
| ↳ organic substance catabolic process | 1433 | 76 | 40.03 | 1.90 | + | 1.60E-03 |
| regulation of protein transport | 739 | 44 | 20.64 | 2.13 | + | 4.99E-02 |
| positive regulation of transport | 1099 | 63 | 30.70 | 2.05 | + | 1.93E-03 |
| cellular response to stress | 1446 | 74 | 40.40 | 1.83 | + | 9.84E-03 |
| Unclassified | 1901 | 14 | 53.11 | .26 | - | 0.00E00 |
| sensory perception | 1642 | 7 | 45.87 | .15 | - | 7.96E-09 |
| G protein-coupled receptor signaling pathway | 1851 | 7 | 51.71 | .14 | - | 3.75E-11 |

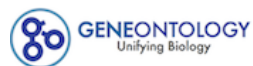

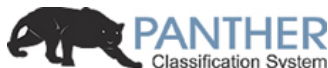

LOGIN REGISTER CONTACT US

Home About PANTHER Data PANTHER Tools PANTHER Services Workspace Downloads Help/Tutorial

**PANTHER 15.0 released!**

Analysis Summary: Please report in publication ?

**Analysis Type:** PANTHER Overrepresentation Test (Released 20200407)

**Annotation Version and Release Date:** GO Ontology database Released 2020-02-21

**Analyzed List:** upload\_1 (Mus musculus)

Change

**Reference List:** Mus musculus (all genes in database)

Change

**Annotation Data Set:** GO molecular function complete ?

**Test Type:** ☒ Fisher's Exact ☐ Binomial

**Correction:** ☐ Calculate False Discovery Rate ☒ Use the Bonferroni correction for multiple testing ? ☐ No correction

**Results** ?

|  | Reference list | upload_1 |
| --- | --- | --- |
| Uniquely Mapped IDs: | 22265 out of 22265 | 622 out of 622 |
| Unmapped IDs: | 0 | 0 |
| Multiple mapping information: | 0 | 0 |

Bonferroni count: 2756

Export Table XML with user input ids JSON with user input ids

Displaying only results for Bonferroni-corrected for $P < 0.05$, click here to display all results

|  | Mus musculus (REF) | upload_1 (▼ Hierarchy NEW! ?) |  |  |  |  |
| --- | --- | --- | --- | --- | --- | --- |
| GO molecular function complete | # | # | expected | Fold Enrichment | +/- | P value |
| N6-methyladenosine-containing RNA binding | 8 | 6 | .22 | 26.85 | + | 2.70E-03 |
| ↳ RNA binding | 1086 | 202 | 30.34 | 6.66 | + | 9.99E-97 |
| ↳ nucleic acid binding | 3161 | 236 | 88.31 | 2.67 | + | 2.21E-43 |
| ↳ organic cyclic compound binding | 5175 | 346 | 144.57 | 2.39 | + | 6.07E-62 |
| ↳ binding | 13351 | 568 | 372.98 | 1.52 | + | 1.11E-64 |
| ↳ heterocyclic compound binding | 5072 | 344 | 141.69 | 2.43 | + | 7.35E-63 |
| RNA stem-loop binding | 14 | 8 | .39 | 20.45 | + | 1.79E-04 |
| sequence-specific mRNA binding | 13 | 6 | .36 | 16.52 | + | 2.17E-02 |
| ↳ mRNA binding | 272 | 79 | 7.60 | 10.40 | + | 5.27E-46 |
| aminoacyl-tRNA ligase activity | 41 | 18 | 1.15 | 15.72 | + | 3.25E-11 |
| ↳ ligase activity, forming carbon-oxygen bonds | 41 | 18 | 1.15 | 15.72 | + | 3.25E-11 |
| ↳ ligase activity | 146 | 25 | 4.08 | 6.13 | + | 2.04E-08 |
| ↳ catalytic activity | 5659 | 230 | 158.09 | 1.45 | + | 1.02E-06 |
| ↳ catalytic activity, acting on a tRNA | 113 | 18 | 3.16 | 5.70 | + | 4.82E-05 |
| ↳ catalytic activity, acting on RNA | 337 | 41 | 9.41 | 4.35 | + | 1.42E-10 |
| RNA cap binding | 17 | 7 | .47 | 14.74 | + | 6.75E-03 |
| mRNA 5'-UTR binding | 24 | 9 | .67 | 13.42 | + | 4.53E-04 |
| mRNA 3'-UTR AU-rich region binding | 27 | 10 | .75 | 13.26 | + | 1.01E-04 |
| ↳ mRNA 3'-UTR binding | 87 | 29 | 2.43 | 11.93 | + | 1.13E-16 |
| ↳ AU-rich element binding | 28 | 10 | .78 | 12.78 | + | 1.34E-04 |
| RNA helicase activity | 53 | 17 | 1.48 | 11.48 | + | 1.10E-08 |
| ↳helicase activity | 148 | 21 | 4.13 | 5.08 | + | 1.98E-05 |
| ↳ATPase activity, coupled | 279 | 28 | 7.79 | 3.59 | + | 7.25E-05 |
| ↳ATPase activity | 412 | 37 | 11.51 | 3.21 | + | 6.91E-06 |
| ↳nucleoside-triphosphatase activity | 748 | 61 | 20.90 | 2.92 | + | 1.79E-09 |
| ↳pyrophosphatase activity | 801 | 66 | 22.38 | 2.95 | + | 1.20E-10 |
| ↳hydrolase activity, acting on acid anhydrides, in phosphorus-containing anhydrides | 804 | 66 | 22.46 | 2.94 | + | 1.41E-10 |
| ↳hydrolase activity, acting on acid anhydrides | 804 | 66 | 22.46 | 2.94 | + | 1.41E-10 |
| ↳hydrolase activity | 2449 | 113 | 68.42 | 1.65 | + | 5.50E-04 |
| translation repressor activity | 22 | 7 | .61 | 11.39 | + | 2.71E-02 |
| ↳translation regulator activity | 126 | 40 | 3.52 | 11.36 | + | 3.59E-23 |
| translation initiation factor activity | 48 | 15 | 1.34 | 11.19 | + | 2.78E-07 |
| ↳translation factor activity, RNA binding | 77 | 25 | 2.15 | 11.62 | + | 7.39E-14 |
| ↳translation regulator activity, nucleic acid binding | 95 | 29 | 2.65 | 10.93 | + | 8.33E-16 |
| pre-mRNA binding | 33 | 10 | .92 | 10.85 | + | 4.81E-04 |
| ribosome binding | 64 | 19 | 1.79 | 10.63 | + | 1.89E-09 |
| ↳ribonucleoprotein complex binding | 146 | 30 | 4.08 | 7.36 | + | 2.28E-12 |
| ↳protein-containing complex binding | 1451 | 119 | 40.54 | 2.94 | + | 1.47E-21 |
| Ran GTPase binding | 34 | 10 | .95 | 10.53 | + | 6.08E-04 |
| ↳Ras GTPase binding | 399 | 38 | 11.15 | 3.41 | + | 9.18E-07 |
| ↳small GTPase binding | 416 | 40 | 11.62 | 3.44 | + | 2.39E-07 |
| ↳GTPase binding | 515 | 51 | 14.39 | 3.54 | + | 1.94E-10 |
| ↳enzyme binding | 2331 | 176 | 65.12 | 2.70 | + | 2.47E-30 |
| ↳protein binding | 9112 | 438 | 254.55 | 1.72 | + | 4.29E-45 |
| poly-purine tract binding | 32 | 9 | .89 | 10.07 | + | 3.39E-03 |
| ↳single-stranded RNA binding | 89 | 22 | 2.49 | 8.85 | + | 6.75E-10 |
| double-stranded RNA binding | 82 | 23 | 2.29 | 10.04 | + | 1.96E-11 |
| miRNA binding | 30 | 8 | .84 | 9.55 | + | 1.87E-02 |
| ↳regulatory RNA binding | 43 | 9 | 1.20 | 7.49 | + | 2.73E-02 |
| ADP binding | 40 | 10 | 1.12 | 8.95 | + | 2.17E-03 |
| ↳adenyl ribonucleotide binding | 1453 | 115 | 40.59 | 2.83 | + | 1.65E-19 |
| ↳adenyl nucleotide binding | 1464 | 117 | 40.90 | 2.86 | + | 3.03E-20 |
| ↳purine nucleotide binding | 1798 | 142 | 50.23 | 2.83 | + | 4.26E-25 |
| ↳nucleotide binding | 2030 | 155 | 56.71 | 2.73 | + | 2.11E-26 |
| ↳small molecule binding | 2421 | 173 | 67.63 | 2.56 | + | 6.35E-27 |
| ↳nucleoside phosphate binding | 2030 | 155 | 56.71 | 2.73 | + | 2.11E-26 |
| ↳purine ribonucleotide binding | 1786 | 140 | 49.89 | 2.81 | + | 2.31E-24 |
| ↳ribonucleotide binding | 1802 | 141 | 50.34 | 2.80 | + | 1.68E-24 |
| ↳carbohydrate derivative binding | 2123 | 148 | 59.31 | 2.50 | + | 4.07E-21 |
| ↳anion binding | 2703 | 179 | 75.51 | 2.37 | + | 2.94E-24 |
| ↳ion binding | 5485 | 244 | 153.23 | 1.59 | + | 7.14E-12 |
| structural constituent of cytoskeleton | 64 | 15 | 1.79 | 8.39 | + | 8.21E-06 |
| ↳structural molecule activity | 576 | 53 | 16.09 | 3.29 | + | 8.99E-10 |
| Hsp90 protein binding | 47 | 10 | 1.31 | 7.62 | + | 7.70E-03 |
| ↳heat shock protein binding | 140 | 23 | 3.91 | 5.88 | + | 2.94E-07 |
| unfolded protein binding | 92 | 19 | 2.57 | 7.39 | + | 4.23E-07 |
| rRNA binding | 70 | 13 | 1.96 | 6.65 | + | 9.92E-04 |
| single-stranded DNA binding | 111 | 16 | 3.10 | 5.16 | + | 1.01E-03 |
| ubiquitin protein ligase binding | 308 | 35 | 8.60 | 4.07 | + | 5.93E-08 |
| ↳ubiquitin-like protein ligase binding | 323 | 36 | 9.02 | 3.99 | + | 5.04E-08 |
| protein C-terminus binding | 231 | 25 | 6.45 | 3.87 | + | 1.07E-04 |
| Rho GTPase binding | 162 | 17 | 4.53 | 3.76 | + | 2.30E-02 |
| actin filament binding | 201 | 21 | 5.62 | 3.74 | + | 2.20E-03 |
| ↳ actin binding | 426 | 37 | 11.90 | 3.11 | + | 1.58E-05 |
| ↳ cytoskeletal protein binding | 976 | 81 | 27.27 | 2.97 | + | 7.56E-14 |
| microtubule binding | 247 | 25 | 6.90 | 3.62 | + | 3.48E-04 |
| ↳ tubulin binding | 346 | 32 | 9.67 | 3.31 | + | 4.43E-05 |
| ATP binding | 1390 | 113 | 38.83 | 2.91 | + | 5.67E-20 |
| ↳ drug binding | 1644 | 128 | 45.93 | 2.79 | + | 1.34E-21 |
| ↳ purine ribonucleoside triphosphate binding | 1713 | 138 | 47.85 | 2.88 | + | 4.86E-25 |
| GTP binding | 355 | 27 | 9.92 | 2.72 | + | 1.87E-02 |
| ↳ purine ribonucleoside binding | 362 | 28 | 10.11 | 2.77 | + | 9.43E-03 |
| ↳ ribonucleoside binding | 365 | 29 | 10.20 | 2.84 | + | 3.85E-03 |
| ↳ nucleoside binding | 375 | 29 | 10.48 | 2.77 | + | 6.34E-03 |
| ↳ purine nucleoside binding | 366 | 28 | 10.22 | 2.74 | + | 1.15E-02 |
| ↳ guanyl ribonucleotide binding | 379 | 29 | 10.59 | 2.74 | + | 7.71E-03 |
| ↳ guanyl nucleotide binding | 379 | 29 | 10.59 | 2.74 | + | 7.71E-03 |
| protein kinase binding | 727 | 54 | 20.31 | 2.66 | + | 1.02E-06 |
| ↳ kinase binding | 813 | 58 | 22.71 | 2.55 | + | 9.94E-07 |
| protein domain specific binding | 786 | 55 | 21.96 | 2.50 | + | 5.38E-06 |
| identical protein binding | 1983 | 120 | 55.40 | 2.17 | + | 1.17E-11 |
| Unclassified | 2116 | 17 | 59.11 | .29 | - | 0.00E00 |
| signaling receptor activity | 2339 | 2 | 65.34 | .03 | - | 3.30E-23 |
| ↳ molecular transducer activity | 2344 | 2 | 65.48 | .03 | - | 3.35E-23 |

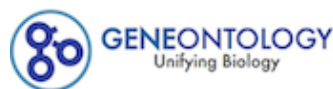

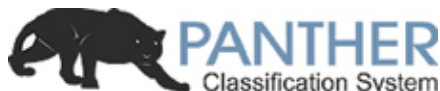

LOGIN REGISTER CONTACT US

Home About PANTHER Data PANTHER Tools PANTHER Services Workspace Downloads Help/Tutorial

### PANTHER 15.0 released!

**Analysis Summary:** Please report in publication ?

**Analysis Type:** PANTHER Overrepresentation Test (Released 20200407)

**Annotation Version and Release Date:** GO Ontology database Released 2020-02-21

**Analyzed List:** upload\_1 (Mus musculus)

Change

**Reference List:** Mus musculus (all genes in database)

Change

**Annotation Data Set:**  ?

**Test Type:** ☒ Fisher's Exact ☐ Binomial

**Correction:** ☐ Calculate False Discovery Rate ☒ Use the Bonferroni correction for multiple testing ? ☐ No correction

### Results ?

|  | Reference list | upload_1 |
| --- | --- | --- |
| Uniquely Mapped IDs: | 22265 out of 22265 | 622 out of 622 |
| Unmapped IDs: | 0 | 0 |
| Multiple mapping information: | 0 | 0 |

Bonferroni count: 1452

Export Table XML with user input ids JSON with user input ids

Displaying only results for Bonferroni-corrected for $P < 0.05$, click here to display all results

|  | Mus musculus (REF) | upload_1 (▼ Hierarchy **NEW!** ?) |  |  |  |  |
| --- | --- | --- | --- | --- | --- | --- |
| GO cellular component complete | # | # | expected | Fold Enrichment | +/- | P value |
| dendritic filopodium | 5 | 5 | .14 | 35.80 | + | 4.77E-03 |
| ↳ dendrite | 705 | 68 | 19.70 | 3.45 | + | 1.77E-14 |
| ↳ neuron projection | 1516 | 112 | 42.35 | 2.64 | + | 6.30E-17 |
| ↳ plasma membrane bounded cell projection | 2281 | 136 | 63.72 | 2.13 | + | 1.30E-13 |
| ↳ cell projection | 2499 | 151 | 69.81 | 2.16 | + | 4.02E-16 |
| ↳ cellular anatomical entity | 18638 | 604 | 520.68 | 1.16 | + | 3.98E-23 |
| ↳ dendritic tree | 708 | 68 | 19.78 | 3.44 | + | 2.17E-14 |
| ↳ somatodendritic compartment | 1028 | 92 | 28.72 | 3.20 | + | 1.84E-18 |
| ↳filopodium | 93 | 12 | 2.60 | 4.62 | + | 4.14E-02 |
| ↳actin-based cell projection | 216 | 22 | 6.03 | 3.65 | + | 9.42E-04 |
| eukaryotic translation initiation factor 3 complex, eIF3m | 8 | 6 | .22 | 26.85 | + | 1.42E-03 |
| ↳eukaryotic translation initiation factor 3 complex | 15 | 11 | .42 | 26.25 | + | 4.24E-08 |
| ↳cytoplasm | 10945 | 546 | 305.76 | 1.79 | + | 9.57E-87 |
| ↳intracellular | 13691 | 601 | 382.47 | 1.57 | + | 3.23E-92 |
| ↳protein-containing complex | 5309 | 340 | 148.31 | 2.29 | + | 9.02E-56 |
| chaperonin-containing T-complex | 10 | 7 | .28 | 25.06 | + | 2.36E-04 |
| ↳chaperone complex | 22 | 12 | .61 | 19.52 | + | 6.74E-08 |
| ↳cytosol | 3533 | 271 | 98.70 | 2.75 | + | 7.79E-55 |
| aminoacyl-tRNA synthetase multienzyme complex | 12 | 8 | .34 | 23.86 | + | 3.90E-05 |
| zona pellucida receptor complex | 13 | 7 | .36 | 19.27 | + | 8.76E-04 |
| messenger ribonucleoprotein complex | 13 | 6 | .36 | 16.52 | + | 1.15E-02 |
| ↳ribonucleoprotein complex | 703 | 103 | 19.64 | 5.24 | + | 2.66E-37 |
| postsynaptic cytoskeleton | 19 | 8 | .53 | 15.07 | + | 5.81E-04 |
| ↳postsynapse | 727 | 87 | 20.31 | 4.28 | + | 3.76E-25 |
| ↳synapse | 1435 | 143 | 40.09 | 3.57 | + | 1.02E-35 |
| ↳cell junction | 2028 | 162 | 56.65 | 2.86 | + | 2.60E-30 |
| ↳cytoskeleton | 2135 | 131 | 59.64 | 2.20 | + | 8.51E-14 |
| ↳intracellular non-membrane-bounded organelle | 4162 | 252 | 116.27 | 2.17 | + | 4.42E-32 |
| ↳intracellular organelle | 11954 | 540 | 333.95 | 1.62 | + | 5.06E-65 |
| ↳organelle | 12303 | 543 | 343.70 | 1.58 | + | 1.01E-61 |
| ↳non-membrane-bounded organelle | 4181 | 253 | 116.80 | 2.17 | + | 3.09E-32 |
| cytoplasmic stress granule | 66 | 26 | 1.84 | 14.10 | + | 1.79E-16 |
| ↳cytoplasmic ribonucleoprotein granule | 206 | 46 | 5.75 | 7.99 | + | 9.84E-22 |
| ↳ribonucleoprotein granule | 217 | 50 | 6.06 | 8.25 | + | 2.39E-24 |
| ↳supramolecular complex | 1197 | 122 | 33.44 | 3.65 | + | 1.47E-30 |
| postsynaptic cytosol | 26 | 9 | .73 | 12.39 | + | 4.16E-04 |
| ↳region of cytosol | 40 | 10 | 1.12 | 8.95 | + | 1.14E-03 |
| endoplasmic reticulum exit site | 21 | 7 | .59 | 11.93 | + | 1.11E-02 |
| ↳endoplasmic reticulum | 1639 | 83 | 45.79 | 1.81 | + | 4.19E-04 |
| ↳endomembrane system | 3884 | 195 | 108.50 | 1.80 | + | 1.56E-13 |
| ↳intracellular membrane-bounded organelle | 10291 | 473 | 287.49 | 1.65 | + | 2.74E-47 |
| ↳membrane-bounded organelle | 11048 | 491 | 308.64 | 1.59 | + | 8.66E-47 |
| myelin sheath | 213 | 66 | 5.95 | 11.09 | + | 1.23E-39 |
| proteasome regulatory particle | 24 | 7 | .67 | 10.44 | + | 2.29E-02 |
| ↳proteasome accessory complex | 26 | 7 | .73 | 9.64 | + | 3.55E-02 |
| ↳proteasome complex | 66 | 11 | 1.84 | 5.97 | + | 1.04E-02 |
| ↳endopeptidase complex | 67 | 11 | 1.87 | 5.88 | + | 1.19E-02 |
| ↳peptidase complex | 93 | 12 | 2.60 | 4.62 | + | 4.14E-02 |
| ↳catalytic complex | 1341 | 74 | 37.46 | 1.98 | + | 9.71E-05 |
| polysome | 73 | 17 | 2.04 | 8.34 | + | 4.17E-07 |
| smooth endoplasmic reticulum | 36 | 8 | 1.01 | 7.95 | + | 3.09E-02 |
| catalytic step 2 spliceosome | 83 | 18 | 2.32 | 7.76 | + | 3.41E-07 |
| ↳spliceosomal complex | 197 | 38 | 5.50 | 6.90 | + | 1.04E-15 |
| ↳nucleus | 6824 | 330 | 190.64 | 1.73 | + | 6.11E-27 |
| mitochondrial nucleoid | 47 | 10 | 1.31 | 7.62 | + | 4.06E-03 |
| ↳mitochondrial matrix | 272 | 33 | 7.60 | 4.34 | + | 2.45E-08 |
| ↳intracellular organelle lumen | 4311 | 242 | 120.43 | 2.01 | + | 2.85E-25 |
| ↳organelle lumen | 4312 | 242 | 120.46 | 2.01 | + | 2.88E-25 |
| ↳membrane-enclosed lumen | 4312 | 242 | 120.46 | 2.01 | + | 2.88E-25 |
| ↳mitochondrion | 1803 | 124 | 50.37 | 2.46 | + | 1.31E-16 |
| ↳nucleoid | 47 | 10 | 1.31 | 7.62 | + | 4.06E-03 |
| intercalated disc | 62 | 13 | 1.73 | 7.51 | + | 1.53E-04 |
| ↳cell-cell contact zone | 85 | 13 | 2.37 | 5.47 | + | 3.66E-03 |
| vesicle coat | 53 | 10 | 1.48 | 6.75 | + | 1.04E-02 |
| ↳coated vesicle membrane | 76 | 12 | 2.12 | 5.65 | + | 6.62E-03 |
| ↳vesicle | 2039 | 94 | 56.96 | 1.65 | + | 3.55E-03 |
| ↳organelle membrane | 2118 | 114 | 59.17 | 1.93 | + | 5.67E-08 |
| ↳whole membrane | 1152 | 61 | 32.18 | 1.90 | + | 6.20E-03 |
| ↳coated vesicle | 188 | 18 | 5.25 | 3.43 | + | 2.11E-02 |
| ↳bounding membrane of organelle | 1098 | 56 | 30.67 | 1.83 | + | 3.67E-02 |
| ↳membrane coat | 95 | 13 | 2.65 | 4.90 | + | 1.09E-02 |
| ↳coated membrane | 95 | 13 | 2.65 | 4.90 | + | 1.09E-02 |
| U2-type spliceosomal complex | 87 | 16 | 2.43 | 6.58 | + | 2.67E-05 |
| small nuclear ribonucleoprotein complex | 60 | 10 | 1.68 | 5.97 | + | 2.71E-02 |
| ↳Sm-like protein family complex | 71 | 12 | 1.98 | 6.05 | + | 3.53E-03 |
| P-body | 72 | 12 | 2.01 | 5.97 | + | 4.02E-03 |
| Golgi-associated vesicle membrane | 62 | 10 | 1.73 | 5.77 | + | 3.49E-02 |
| small ribosomal subunit | 78 | 12 | 2.18 | 5.51 | + | 8.41E-03 |
| ↳ribosomal subunit | 202 | 20 | 5.64 | 3.54 | + | 4.49E-03 |
| ↳ribosome | 234 | 27 | 6.54 | 4.13 | + | 4.64E-06 |
| nuclear matrix | 88 | 13 | 2.46 | 5.29 | + | 5.16E-03 |
| ↳nuclear periphery | 113 | 17 | 3.16 | 5.39 | + | 1.33E-04 |
| ↳nuclear lumen | 3899 | 207 | 108.92 | 1.90 | + | 1.57E-17 |
| growth cone | 205 | 30 | 5.73 | 5.24 | + | 3.11E-09 |
| ↳site of polarized growth | 213 | 31 | 5.95 | 5.21 | + | 1.51E-09 |
| ↳distal axon | 379 | 37 | 10.59 | 3.49 | + | 4.41E-07 |
| ↳axon | 715 | 66 | 19.97 | 3.30 | + | 4.11E-13 |
| dendritic spine | 193 | 25 | 5.39 | 4.64 | + | 2.13E-06 |
| ↳neuron spine | 199 | 26 | 5.56 | 4.68 | + | 8.48E-07 |
| postsynaptic density | 396 | 49 | 11.06 | 4.43 | + | 1.27E-13 |
| ↳asymmetric synapse | 400 | 50 | 11.17 | 4.47 | + | 4.16E-14 |
| ↳neuron to neuron synapse | 427 | 52 | 11.93 | 4.36 | + | 2.54E-14 |
| ↳postsynaptic specialization | 435 | 50 | 12.15 | 4.11 | + | 9.00E-13 |
| nuclear speck | 315 | 37 | 8.80 | 4.20 | + | 3.40E-09 |
| ↳nuclear body | 682 | 55 | 19.05 | 2.89 | + | 2.06E-08 |
| ↳nucleoplasm | 3324 | 181 | 92.86 | 1.95 | + | 1.25E-15 |
| glutamatergic synapse | 508 | 57 | 14.19 | 4.02 | + | 2.00E-14 |
| perikaryon | 138 | 15 | 3.86 | 3.89 | + | 2.80E-02 |
| ↳neuronal cell body | 710 | 68 | 19.83 | 3.43 | + | 2.48E-14 |
| ↳cell body | 801 | 76 | 22.38 | 3.40 | + | 4.27E-16 |
| lamellipodium | 166 | 17 | 4.64 | 3.67 | + | 1.63E-02 |
| ↳cell leading edge | 389 | 37 | 10.87 | 3.40 | + | 8.58E-07 |
| perinuclear region of cytoplasm | 657 | 67 | 18.35 | 3.65 | + | 2.28E-15 |
| microtubule | 420 | 42 | 11.73 | 3.58 | + | 1.34E-08 |
| ↳microtubule cytoskeleton | 1173 | 74 | 32.77 | 2.26 | + | 3.70E-07 |
| ↳polymeric cytoskeletal fiber | 682 | 61 | 19.05 | 3.20 | + | 2.23E-11 |
| ↳supramolecular fiber | 902 | 76 | 25.20 | 3.02 | + | 1.99E-13 |
| ↳supramolecular polymer | 909 | 76 | 25.39 | 2.99 | + | 2.94E-13 |
| cell cortex | 316 | 31 | 8.83 | 3.51 | + | 1.12E-05 |
| mitochondrial protein complex | 260 | 23 | 7.26 | 3.17 | + | 4.87E-03 |
| nuclear envelope | 423 | 34 | 11.82 | 2.88 | + | 1.95E-04 |
| ↳organelle envelope | 1069 | 71 | 29.86 | 2.38 | + | 9.96E-08 |
| ↳envelope | 1070 | 71 | 29.89 | 2.38 | + | 1.03E-07 |
| presynapse | 590 | 47 | 16.48 | 2.85 | + | 9.79E-07 |
| actin cytoskeleton | 497 | 38 | 13.88 | 2.74 | + | 1.19E-04 |
| mitochondrial membrane | 617 | 38 | 17.24 | 2.20 | + | 1.90E-02 |
| nucleolus | 808 | 46 | 22.57 | 2.04 | + | 1.84E-02 |
| plasma membrane region | 1218 | 63 | 34.03 | 1.85 | + | 7.52E-03 |
| Golgi apparatus | 1419 | 73 | 39.64 | 1.84 | + | 1.38E-03 |
| extracellular space | 1951 | 21 | 54.50 | .39 | - | 2.98E-04 |
| ↳extracellular region | 2797 | 42 | 78.14 | .54 | - | 6.46E-03 |
| integral component of plasma membrane | 1499 | 12 | 41.88 | .29 | - | 1.01E-04 |
| ↳intrinsic component of plasma membrane | 1576 | 13 | 44.03 | .30 | - | 8.06E-05 |
| ↳intrinsic component of membrane | 6027 | 59 | 168.37 | .35 | - | 1.03E-23 |
| ↳integral component of membrane | 5854 | 55 | 163.54 | .34 | - | 3.40E-24 |
| Unclassified | 1476 | 10 | 41.23 | .24 | - | 0.00E00 |

About | Release Information | Contact Us | System Requirements | Privacy Policy | Disclaimer

© Copyright 2020 Paul Thomas All Rights Reserved.

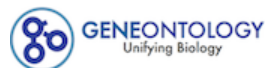

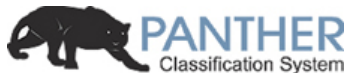

LOGIN REGISTER CONTACT US

Home About PANTHER Data PANTHER Tools PANTHER Services Workspace Downloads Help/Tutorial

**PANTHER 15.0 released!**

Analysis Summary: Please report in publication ?

**Analysis Type:** PANTHER Overrepresentation Test (Released 20200407)

**Annotation Version and Release Date:** GO Ontology database Released 2020-02-21

**Analyzed List:** upload\_1 (Mus musculus)

Change

**Reference List:** Mus musculus (all genes in database)

Change

**Annotation Data Set:** GO biological process complete ?

**Test Type:** ☒ Fisher's Exact ☐ Binomial

**Correction:** ☐ Calculate False Discovery Rate ☒ Use the Bonferroni correction for multiple testing ? ☐ No correction

**Results** ?

|  | Reference list | upload_1 |
| --- | --- | --- |
| Uniquely Mapped IDs: | 22265 out of 22265 | 1344 out of 1344 |
| Unmapped IDs: | 0 | 0 |
| Multiple mapping information: | 0 | 0 |

Bonferroni count: 8894

Export Table XML with user input ids JSON with user input ids

Displaying only results for Bonferroni-corrected for P < 0.05, click here to display all results

|  | Mus musculus (REF) | upload_1 (Hierarchy NEW! ?) |  |  |  |  |
| --- | --- | --- | --- | --- | --- | --- |
| GO biological process complete | # | # | expected | Fold Enrichment | +/- | P value |
| fatty acid beta-oxidation using acyl-CoA dehydrogenase | 10 | 9 | .60 | 14.91 | + | 2.99E-03 |
| ↳ fatty acid beta-oxidation | 49 | 25 | 2.96 | 8.45 | + | 1.29E-09 |
| ↳ fatty acid oxidation | 69 | 28 | 4.17 | 6.72 | + | 3.89E-09 |
| ↳ fatty acid metabolic process | 309 | 67 | 18.65 | 3.59 | + | 5.29E-13 |
| ↳ monocarboxylic acid metabolic process | 467 | 107 | 28.19 | 3.80 | + | 3.66E-24 |
| ↳ carboxylic acid metabolic process | 761 | 198 | 45.94 | 4.31 | + | 1.12E-55 |
| ↳ oxoacid metabolic process | 803 | 200 | 48.47 | 4.13 | + | 1.08E-53 |
| ↳ organic acid metabolic process | 829 | 203 | 50.04 | 4.06 | + | 1.44E-53 |
| ↳ cellular metabolic process | 6331 | 729 | 382.16 | 1.91 | + | 8.44E-78 |
| ↳ metabolic process | 7222 | 788 | 435.95 | 1.81 | + | 4.41E-77 |
| ↳ cellular process | 14059 | 1142 | 848.65 | 1.35 | + | 4.52E-63 |
| ↳ small molecule metabolic process | 1432 | 263 | 86.44 | 3.04 | + | 1.26E-49 |
| ↳ organic substance metabolic process | 6701 | 748 | 404.50 | 1.85 | + | 6.13E-75 |
| ↳ cellular lipid metabolic process | 801 | 91 | 48.35 | 1.88 | + | 5.09E-04 |
| ↳ lipid metabolic process | 1063 | 112 | 64.17 | 1.75 | + | 5.95E-04 |
| ↳ primary metabolic process | 6269 | 699 | 378.42 | 1.85 | + | 9.74E-67 |
| ↳ lipid oxidation | 75 | 28 | 4.53 | 6.18 | + | 1.98E-08 |
| ↳ lipid modification | 182 | 35 | 10.99 | 3.19 | + | 2.11E-04 |
| ↳oxidation-reduction process | 787 | 136 | 47.51 | 2.86 | + | 5.24E-21 |
| ↳fatty acid catabolic process | 73 | 28 | 4.41 | 6.35 | + | 1.17E-08 |
| ↳cellular lipid catabolic process | 164 | 32 | 9.90 | 3.23 | + | 6.20E-04 |
| ↳cellular catabolic process | 1492 | 215 | 90.06 | 2.39 | + | 1.79E-25 |
| ↳catabolic process | 1730 | 248 | 104.43 | 2.37 | + | 5.42E-30 |
| ↳lipid catabolic process | 259 | 44 | 15.63 | 2.81 | + | 9.62E-05 |
| ↳organic substance catabolic process | 1433 | 222 | 86.50 | 2.57 | + | 1.49E-30 |
| ↳monocarboxylic acid catabolic process | 93 | 35 | 5.61 | 6.23 | + | 2.95E-11 |
| ↳carboxylic acid catabolic process | 194 | 64 | 11.71 | 5.47 | + | 3.51E-20 |
| ↳organic acid catabolic process | 194 | 64 | 11.71 | 5.47 | + | 3.51E-20 |
| ↳small molecule catabolic process | 290 | 83 | 17.51 | 4.74 | + | 1.81E-23 |
| positive regulation of establishment of protein localization to telomere | 10 | 8 | .60 | 13.25 | + | 2.52E-02 |
| ↳positive regulation of establishment of protein localization | 462 | 71 | 27.89 | 2.55 | + | 2.88E-07 |
| ↳positive regulation of biological process | 6100 | 522 | 368.22 | 1.42 | + | 1.25E-14 |
| ↳regulation of biological process | 11534 | 822 | 696.24 | 1.18 | + | 2.12E-07 |
| ↳biological regulation | 12152 | 877 | 733.54 | 1.20 | + | 1.16E-10 |
| ↳regulation of establishment of protein localization | 771 | 105 | 46.54 | 2.26 | + | 3.15E-09 |
| ↳regulation of protein localization | 1075 | 147 | 64.89 | 2.27 | + | 3.34E-14 |
| ↳regulation of localization | 2886 | 290 | 174.21 | 1.66 | + | 3.26E-13 |
| ↳regulation of cellular protein localization | 560 | 93 | 33.80 | 2.75 | + | 2.18E-12 |
| ↳regulation of cellular localization | 837 | 118 | 50.52 | 2.34 | + | 1.55E-11 |
| ↳positive regulation of cellular protein localization | 308 | 55 | 18.59 | 2.96 | + | 2.75E-07 |
| ↳regulation of establishment of protein localization to telomere | 11 | 8 | .66 | 12.05 | + | 4.13E-02 |
| viral translation | 14 | 10 | .85 | 11.83 | + | 2.89E-03 |
| ↳viral gene expression | 20 | 11 | 1.21 | 9.11 | + | 5.11E-03 |
| ↳viral process | 131 | 29 | 7.91 | 3.67 | + | 2.41E-04 |
| ↳symbiotic process | 228 | 46 | 13.76 | 3.34 | + | 3.44E-07 |
| ↳interspecies interaction between organisms | 1474 | 135 | 88.98 | 1.52 | + | 4.01E-02 |
| tricarboxylic acid metabolic process | 15 | 10 | .91 | 11.04 | + | 4.57E-03 |
| alternative mRNA splicing, via spliceosome | 18 | 11 | 1.09 | 10.12 | + | 2.32E-03 |
| ↳mRNA splicing, via spliceosome | 196 | 50 | 11.83 | 4.23 | + | 2.28E-11 |
| ↳mRNA processing | 405 | 104 | 24.45 | 4.25 | + | 7.65E-27 |
| ↳RNA processing | 752 | 139 | 45.39 | 3.06 | + | 4.66E-24 |
| ↳RNA metabolic process | 1232 | 197 | 74.37 | 2.65 | + | 3.29E-28 |
| ↳nucleic acid metabolic process | 1767 | 221 | 106.66 | 2.07 | + | 7.36E-19 |
| ↳macromolecule metabolic process | 5056 | 530 | 305.20 | 1.74 | + | 3.40E-36 |
| ↳nucleobase-containing compound metabolic process | 2191 | 308 | 132.26 | 2.33 | + | 1.98E-37 |
| ↳cellular nitrogen compound metabolic process | 2789 | 436 | 168.35 | 2.59 | + | 1.66E-70 |
| ↳nitrogen compound metabolic process | 5718 | 640 | 345.16 | 1.85 | + | 1.26E-58 |
| ↳organic cyclic compound metabolic process | 2616 | 359 | 157.91 | 2.27 | + | 4.87E-43 |
| ↳heterocycle metabolic process | 2328 | 332 | 140.53 | 2.36 | + | 2.59E-42 |
| ↳cellular aromatic compound metabolic process | 2402 | 337 | 144.99 | 2.32 | + | 1.24E-41 |
| ↳gene expression | 1616 | 277 | 97.55 | 2.84 | + | 2.05E-47 |
| ↳mRNA metabolic process | 531 | 130 | 32.05 | 4.06 | + | 1.33E-32 |
| ↳RNA splicing, via transesterification reactions with bulged adenosine as nucleophile | 196 | 50 | 11.83 | 4.23 | + | 2.28E-11 |
| ↳RNA splicing, via transesterification reactions | 196 | 50 | 11.83 | 4.23 | + | 2.28E-11 |
| ↳RNA splicing | 318 | 87 | 19.20 | 4.53 | + | 1.41E-23 |
| acyl-CoA biosynthetic process | 20 | 12 | 1.21 | 9.94 | + | 7.64E-04 |
| ↳ purine ribonucleotide biosynthetic process | 115 | 27 | 6.94 | 3.89 | + | 2.54E-04 |
| ↳ purine ribonucleotide metabolic process | 256 | 68 | 15.45 | 4.40 | + | 3.24E-17 |
| ↳ ribonucleotide metabolic process | 266 | 69 | 16.06 | 4.30 | + | 4.98E-17 |
| ↳ nucleotide metabolic process | 352 | 79 | 21.25 | 3.72 | + | 1.36E-16 |
| ↳ nucleoside phosphate metabolic process | 362 | 79 | 21.85 | 3.62 | + | 5.77E-16 |
| ↳ phosphate-containing compound metabolic process | 1670 | 163 | 100.81 | 1.62 | + | 7.63E-05 |
| ↳ phosphorus metabolic process | 1692 | 164 | 102.14 | 1.61 | + | 9.54E-05 |
| ↳ nucleobase-containing small molecule metabolic process | 426 | 91 | 25.71 | 3.54 | + | 2.82E-18 |
| ↳ organophosphate metabolic process | 710 | 104 | 42.86 | 2.43 | + | 8.04E-11 |
| ↳ ribose phosphate metabolic process | 277 | 71 | 16.72 | 4.25 | + | 2.36E-17 |
| ↳ carbohydrate derivative metabolic process | 794 | 102 | 47.93 | 2.13 | + | 1.62E-07 |
| ↳ purine nucleotide metabolic process | 273 | 71 | 16.48 | 4.31 | + | 1.17E-17 |
| ↳ purine-containing compound metabolic process | 313 | 76 | 18.89 | 4.02 | + | 1.42E-17 |
| ↳ organonitrogen compound metabolic process | 4270 | 474 | 257.75 | 1.84 | + | 1.08E-36 |
| ↳ ribonucleotide biosynthetic process | 124 | 28 | 7.49 | 3.74 | + | 2.90E-04 |
| ↳ nucleotide biosynthetic process | 171 | 37 | 10.32 | 3.58 | + | 5.26E-06 |
| ↳ nucleoside phosphate biosynthetic process | 178 | 39 | 10.74 | 3.63 | + | 1.32E-06 |
| ↳ organophosphate biosynthetic process | 366 | 52 | 22.09 | 2.35 | + | 1.42E-03 |
| ↳ organic substance biosynthetic process | 2079 | 282 | 125.50 | 2.25 | + | 3.44E-31 |
| ↳ biosynthetic process | 2161 | 292 | 130.45 | 2.24 | + | 3.82E-32 |
| ↳ heterocycle biosynthetic process | 718 | 79 | 43.34 | 1.82 | + | 1.47E-02 |
| ↳ cellular biosynthetic process | 1985 | 269 | 119.82 | 2.24 | + | 2.18E-29 |
| ↳ organic cyclic compound biosynthetic process | 848 | 93 | 51.19 | 1.82 | + | 1.65E-03 |
| ↳ cellular nitrogen compound biosynthetic process | 1118 | 197 | 67.49 | 2.92 | + | 2.49E-33 |
| ↳ ribose phosphate biosynthetic process | 131 | 30 | 7.91 | 3.79 | + | 7.18E-05 |
| ↳ purine nucleotide biosynthetic process | 125 | 29 | 7.55 | 3.84 | + | 9.90E-05 |
| ↳ purine-containing compound biosynthetic process | 130 | 30 | 7.85 | 3.82 | + | 6.17E-05 |
| ↳ organonitrogen compound biosynthetic process | 1070 | 197 | 64.59 | 3.05 | + | 1.09E-35 |
| ↳ coenzyme biosynthetic process | 99 | 24 | 5.98 | 4.02 | + | 8.86E-04 |
| ↳ coenzyme metabolic process | 204 | 57 | 12.31 | 4.63 | + | 7.53E-15 |
| ↳ cofactor metabolic process | 368 | 90 | 22.21 | 4.05 | + | 1.57E-21 |
| ↳ cofactor biosynthetic process | 163 | 35 | 9.84 | 3.56 | + | 1.81E-05 |
| ↳ amide biosynthetic process | 421 | 138 | 25.41 | 5.43 | + | 6.42E-47 |
| ↳ cellular amide metabolic process | 665 | 184 | 40.14 | 4.58 | + | 1.00E-54 |
| ↳ ribonucleoside bisphosphate biosynthetic process | 30 | 13 | 1.81 | 7.18 | + | 4.07E-03 |
| ↳ ribonucleoside bisphosphate metabolic process | 94 | 36 | 5.67 | 6.34 | + | 7.68E-12 |
| ↳ nucleoside bisphosphate metabolic process | 94 | 36 | 5.67 | 6.34 | + | 7.68E-12 |
| ↳ nucleoside bisphosphate biosynthetic process | 30 | 13 | 1.81 | 7.18 | + | 4.07E-03 |
| ↳ thioester biosynthetic process | 20 | 12 | 1.21 | 9.94 | + | 7.64E-04 |
| ↳ thioester metabolic process | 78 | 33 | 4.71 | 7.01 | + | 1.22E-11 |
| ↳ sulfur compound metabolic process | 267 | 68 | 16.12 | 4.22 | + | 2.23E-16 |
| ↳ sulfur compound biosynthetic process | 72 | 24 | 4.35 | 5.52 | + | 4.85E-06 |
| ↳ purine nucleoside bisphosphate biosynthetic process | 30 | 13 | 1.81 | 7.18 | + | 4.07E-03 |
| ↳ purine nucleoside bisphosphate metabolic process | 94 | 36 | 5.67 | 6.34 | + | 7.68E-12 |
| ↳ acyl-CoA metabolic process | 78 | 33 | 4.71 | 7.01 | + | 1.22E-11 |
| negative regulation of mRNA splicing, via spliceosome | 24 | 14 | 1.45 | 9.66 | + | 8.30E-05 |
| ↳regulation of mRNA splicing, via spliceosome | 110 | 46 | 6.64 | 6.93 | + | 4.54E-17 |
| ↳regulation of RNA splicing | 145 | 57 | 8.75 | 6.51 | + | 1.04E-20 |
| ↳regulation of gene expression | 3827 | 357 | 231.01 | 1.55 | + | 9.96E-13 |
| ↳regulation of macromolecule metabolic process | 5484 | 494 | 331.04 | 1.49 | + | 1.09E-17 |
| ↳regulation of metabolic process | 5976 | 546 | 360.73 | 1.51 | + | 3.60E-22 |
| ↳regulation of RNA metabolic process | 3165 | 255 | 191.05 | 1.33 | + | 3.10E-02 |
| ↳regulation of nucleobase-containing compound metabolic process | 3401 | 275 | 205.30 | 1.34 | + | 7.35E-03 |
| ↳regulation of primary metabolic process | 5349 | 483 | 322.89 | 1.50 | + | 3.32E-17 |
| ↳regulation of nitrogen compound metabolic process | 5198 | 463 | 313.77 | 1.48 | + | 4.95E-15 |
| ↳regulation of cellular metabolic process | 5552 | 501 | 335.14 | 1.49 | + | 3.22E-18 |
| ↳regulation of cellular process | 10826 | 771 | 653.50 | 1.18 | + | 4.40E-06 |
| ↳regulation of mRNA processing | 149 | 56 | 8.99 | 6.23 | + | 1.56E-19 |
| ↳regulation of mRNA metabolic process | 255 | 93 | 15.39 | 6.04 | + | 1.82E-33 |
| ↳negative regulation of RNA splicing | 29 | 16 | 1.75 | 9.14 | + | 1.33E-05 |
| ↳negative regulation of macromolecule metabolic process | 2494 | 253 | 150.55 | 1.68 | + | 2.40E-11 |
| ↳negative regulation of metabolic process | 2766 | 281 | 166.97 | 1.68 | + | 3.72E-13 |
| ↳negative regulation of biological process | 5169 | 479 | 312.02 | 1.54 | + | 3.15E-19 |
| ↳negative regulation of cellular metabolic process | 2469 | 241 | 149.04 | 1.62 | + | 6.06E-09 |
| ↳negative regulation of cellular process | 4632 | 423 | 279.61 | 1.51 | + | 7.51E-15 |
| ↳negative regulation of nitrogen compound metabolic process | 2272 | 219 | 137.15 | 1.60 | + | 3.10E-07 |
| ↳negative regulation of gene expression | 1643 | 169 | 99.18 | 1.70 | + | 1.09E-06 |
| ↳negative regulation of mRNA processing | 32 | 16 | 1.93 | 8.28 | + | 3.95E-05 |
| ↳negative regulation of mRNA metabolic process | 82 | 33 | 4.95 | 6.67 | + | 3.87E-11 |
| stress granule assembly | 16 | 9 | .97 | 9.32 | + | 4.84E-02 |
| ↳cellular component assembly | 1941 | 244 | 117.17 | 2.08 | + | 1.13E-21 |
| ↳cellular component organization | 5045 | 479 | 304.54 | 1.57 | + | 1.56E-21 |
| ↳cellular component organization or biogenesis | 5239 | 491 | 316.25 | 1.55 | + | 4.37E-21 |
| ↳cellular component biogenesis | 2168 | 259 | 130.87 | 1.98 | + | 1.98E-20 |
| ↳organelle organization | 3054 | 307 | 184.35 | 1.67 | + | 2.60E-14 |
| tricarboxylic acid cycle | 32 | 18 | 1.93 | 9.32 | + | 9.90E-07 |
| ↳aerobic respiration | 66 | 21 | 3.98 | 5.27 | + | 1.13E-04 |
| ↳cellular respiration | 129 | 32 | 7.79 | 4.11 | + | 4.23E-06 |
| ↳energy derivation by oxidation of organic compounds | 191 | 45 | 11.53 | 3.90 | + | 6.10E-09 |
| ↳generation of precursor metabolites and energy | 286 | 74 | 17.26 | 4.29 | + | 2.08E-18 |
| acetyl-CoA metabolic process | 29 | 16 | 1.75 | 9.14 | + | 1.33E-05 |
| glutamate metabolic process | 24 | 13 | 1.45 | 8.97 | + | 5.46E-04 |
| ↳dicarboxylic acid metabolic process | 81 | 36 | 4.89 | 7.36 | + | 1.82E-13 |
| ↳glutamine family amino acid metabolic process | 57 | 21 | 3.44 | 6.10 | + | 1.34E-05 |
| ↳alpha-amino acid metabolic process | 164 | 56 | 9.90 | 5.66 | + | 6.61E-18 |
| ↳cellular amino acid metabolic process | 231 | 79 | 13.94 | 5.67 | + | 1.99E-26 |
| aromatic amino acid family catabolic process | 19 | 10 | 1.15 | 8.72 | + | 2.27E-02 |
| ↳aromatic compound catabolic process | 292 | 56 | 17.63 | 3.18 | + | 1.55E-08 |
| ↳cellular amino acid catabolic process | 90 | 34 | 5.43 | 6.26 | + | 6.69E-11 |
| ↳organonitrogen compound catabolic process | 900 | 116 | 54.33 | 2.14 | + | 5.88E-09 |
| ↳organic cyclic compound catabolic process | 323 | 64 | 19.50 | 3.28 | + | 1.07E-10 |
| ↳aromatic amino acid family metabolic process | 34 | 13 | 2.05 | 6.33 | + | 1.26E-02 |
| translational elongation | 30 | 15 | 1.81 | 8.28 | + | 1.20E-04 |
| ↳cellular macromolecule biosynthetic process | 1193 | 162 | 72.01 | 2.25 | + | 1.04E-15 |
| ↳cellular macromolecule metabolic process | 3958 | 386 | 238.92 | 1.62 | + | 1.85E-17 |
| ↳macromolecule biosynthetic process | 1227 | 163 | 74.07 | 2.20 | + | 3.59E-15 |
| ↳translation | 306 | 112 | 18.47 | 6.06 | + | 4.09E-41 |
| ↳cellular protein metabolic process | 2844 | 288 | 171.67 | 1.68 | + | 1.51E-13 |
| ↳protein metabolic process | 3420 | 332 | 206.44 | 1.61 | + | 7.77E-14 |
| ↳peptide biosynthetic process | 326 | 118 | 19.68 | 6.00 | + | 4.10E-43 |
| ↳peptide metabolic process | 448 | 138 | 27.04 | 5.10 | + | 2.60E-44 |
| tRNA aminoacylation for protein translation | 40 | 20 | 2.41 | 8.28 | + | 4.65E-07 |
| ↳tRNA aminoacylation | 43 | 20 | 2.60 | 7.71 | + | 1.27E-06 |
| ↳amino acid activation | 44 | 20 | 2.66 | 7.53 | + | 1.75E-06 |
| ↳tRNA metabolic process | 165 | 28 | 9.96 | 2.81 | + | 4.42E-02 |
| ↳ncRNA metabolic process | 416 | 55 | 25.11 | 2.19 | + | 4.57E-03 |
| mRNA destabilization | 32 | 16 | 1.93 | 8.28 | + | 3.95E-05 |
| ↳RNA destabilization | 35 | 16 | 2.11 | 7.57 | + | 1.07E-04 |
| ↳regulation of RNA stability | 109 | 43 | 6.58 | 6.54 | + | 5.01E-15 |
| ↳regulation of cellular catabolic process | 698 | 128 | 42.13 | 3.04 | + | 1.34E-21 |
| ↳regulation of catabolic process | 843 | 141 | 50.89 | 2.77 | + | 1.08E-20 |
| ↳posttranscriptional regulation of gene expression | 427 | 135 | 25.78 | 5.24 | + | 2.53E-44 |
| ↳regulation of biological quality | 3959 | 421 | 238.98 | 1.76 | + | 2.67E-27 |
| ↳positive regulation of cellular metabolic process | 3270 | 276 | 197.39 | 1.40 | + | 1.92E-04 |
| ↳positive regulation of cellular process | 5424 | 461 | 327.41 | 1.41 | + | 2.09E-11 |
| ↳positive regulation of metabolic process | 3560 | 310 | 214.90 | 1.44 | + | 6.54E-07 |
| ↳positive regulation of nitrogen compound metabolic process | 3086 | 258 | 186.28 | 1.38 | + | 1.36E-03 |
| ↳positive regulation of macromolecule metabolic process | 3256 | 270 | 196.54 | 1.37 | + | 1.32E-03 |
| ↳positive regulation of cellular catabolic process | 367 | 72 | 22.15 | 3.25 | + | 3.57E-12 |
| ↳positive regulation of catabolic process | 436 | 77 | 26.32 | 2.93 | + | 4.94E-11 |
| ↳positive regulation of mRNA catabolic process | 50 | 23 | 3.02 | 7.62 | + | 6.35E-08 |
| ↳positive regulation of mRNA metabolic process | 93 | 34 | 5.61 | 6.06 | + | 1.46E-10 |
| ↳regulation of mRNA catabolic process | 121 | 49 | 7.30 | 6.71 | + | 7.12E-18 |
| ↳regulation of mRNA stability | 98 | 42 | 5.92 | 7.10 | + | 1.13E-15 |
| ↳negative regulation of translation | 130 | 46 | 7.85 | 5.86 | + | 9.73E-15 |
| ↳negative regulation of cellular protein metabolic process | 1004 | 133 | 60.61 | 2.19 | + | 1.49E-11 |
| ↳regulation of cellular protein metabolic process | 2531 | 274 | 152.78 | 1.79 | + | 4.13E-16 |
| ↳regulation of protein metabolic process | 2723 | 290 | 164.37 | 1.76 | + | 1.66E-16 |
| ↳negative regulation of protein metabolic process | 1072 | 139 | 64.71 | 2.15 | + | 1.35E-11 |
| ↳negative regulation of cellular amide metabolic process | 149 | 49 | 8.99 | 5.45 | + | 8.74E-15 |
| ↳regulation of cellular amide metabolic process | 384 | 113 | 23.18 | 4.87 | + | 4.91E-34 |
| ↳regulation of translation | 333 | 108 | 20.10 | 5.37 | + | 1.52E-35 |
| ↳regulation of cellular macromolecule biosynthetic process | 3333 | 271 | 201.19 | 1.35 | + | 5.96E-03 |
| ↳regulation of macromolecule biosynthetic process | 3419 | 282 | 206.38 | 1.37 | + | 9.94E-04 |
| ↳regulation of biosynthetic process | 3644 | 315 | 219.97 | 1.43 | + | 9.71E-07 |
| ↳regulation of cellular biosynthetic process | 3571 | 299 | 215.56 | 1.39 | + | 8.14E-05 |
| production of miRNAs involved in gene silencing by miRNA | 27 | 13 | 1.63 | 7.98 | + | 1.57E-03 |
| ↳gene silencing by miRNA | 40 | 17 | 2.41 | 7.04 | + | 8.86E-05 |
| ↳posttranscriptional gene silencing by RNA | 47 | 19 | 2.84 | 6.70 | + | 2.34E-05 |
|  |  |  | 4.65 | 5.16 | + | 1.48E-05 |
| ↳gene silencing by RNA | 77 | 24 |  |  |  |  |
| ↳posttranscriptional gene silencing | 48 | 20 | 2.90 | 6.90 | + | 5.88E-06 |
| ↳production of small RNA involved in gene silencing by RNA | 29 | 13 | 1.75 | 7.43 | + | 2.99E-03 |
| ↳dsRNA processing | 29 | 13 | 1.75 | 7.43 | + | 2.99E-03 |
| regulation of alternative mRNA splicing, via spliceosome | 66 | 30 | 3.98 | 7.53 | + | 5.30E-11 |
| translational initiation | 55 | 25 | 3.32 | 7.53 | + | 9.59E-09 |
| NADH metabolic process | 27 | 12 | 1.63 | 7.36 | + | 9.12E-03 |
| alpha-amino acid catabolic process | 75 | 32 | 4.53 | 7.07 | + | 2.72E-11 |
| sulfur amino acid metabolic process | 31 | 13 | 1.87 | 6.95 | + | 5.47E-03 |
| mRNA stabilization | 41 | 17 | 2.47 | 6.87 | + | 1.19E-04 |
| ↳negative regulation of mRNA catabolic process | 51 | 19 | 3.08 | 6.17 | + | 6.91E-05 |
| ↳negative regulation of RNA catabolic process | 61 | 23 | 3.68 | 6.25 | + | 1.54E-06 |
| ↳negative regulation of cellular catabolic process | 241 | 42 | 14.55 | 2.89 | + | 1.09E-04 |
| ↳negative regulation of catabolic process | 301 | 46 | 18.17 | 2.53 | + | 1.01E-03 |
| ↳positive regulation of gene expression | 1995 | 173 | 120.43 | 1.44 | + | 3.99E-02 |
| ↳RNA stabilization | 48 | 19 | 2.90 | 6.56 | + | 3.10E-05 |
| actin filament capping | 32 | 13 | 1.93 | 6.73 | + | 7.30E-03 |
| ↳negative regulation of actin filament depolymerization | 38 | 15 | 2.29 | 6.54 | + | 1.41E-03 |
| ↳negative regulation of cytoskeleton organization | 161 | 33 | 9.72 | 3.40 | + | 1.39E-04 |
| ↳regulation of cytoskeleton organization | 559 | 80 | 33.74 | 2.37 | + | 2.74E-07 |
| ↳regulation of organelle organization | 1270 | 156 | 76.66 | 2.03 | + | 1.50E-11 |
| ↳regulation of cellular component organization | 2537 | 278 | 153.14 | 1.82 | + | 3.79E-17 |
| ↳negative regulation of organelle organization | 392 | 50 | 23.66 | 2.11 | + | 3.26E-02 |
| ↳negative regulation of cellular component organization | 734 | 90 | 44.31 | 2.03 | + | 2.18E-05 |
| ↳negative regulation of protein depolymerization | 72 | 20 | 4.35 | 4.60 | + | 1.61E-03 |
| ↳regulation of protein depolymerization | 89 | 23 | 5.37 | 4.28 | + | 6.08E-04 |
| ↳regulation of protein-containing complex disassembly | 121 | 30 | 7.30 | 4.11 | + | 1.48E-05 |
| ↳negative regulation of protein-containing complex disassembly | 81 | 20 | 4.89 | 4.09 | + | 7.91E-03 |
| ↳negative regulation of supramolecular fiber organization | 154 | 36 | 9.30 | 3.87 | + | 1.45E-06 |
| ↳regulation of supramolecular fiber organization | 368 | 59 | 22.21 | 2.66 | + | 2.40E-06 |
| ↳regulation of actin filament depolymerization | 51 | 17 | 3.08 | 5.52 | + | 1.57E-03 |
| ↳regulation of actin polymerization or depolymerization | 185 | 34 | 11.17 | 3.04 | + | 8.65E-04 |
| ↳regulation of actin filament organization | 271 | 40 | 16.36 | 2.45 | + | 1.33E-02 |
| ↳regulation of actin filament-based process | 409 | 54 | 24.69 | 2.19 | + | 6.27E-03 |
| ↳regulation of actin filament length | 188 | 34 | 11.35 | 3.00 | + | 1.21E-03 |
| ↳regulation of cellular component size | 407 | 62 | 24.57 | 2.52 | + | 5.18E-06 |
| ↳regulation of anatomical structure size | 571 | 84 | 34.47 | 2.44 | + | 3.47E-08 |
| ↳negative regulation of actin filament polymerization | 58 | 18 | 3.50 | 5.14 | + | 1.67E-03 |
| ↳regulation of actin filament polymerization | 168 | 32 | 10.14 | 3.16 | + | 1.00E-03 |
| ↳regulation of protein polymerization | 224 | 44 | 13.52 | 3.25 | + | 1.92E-06 |
| ↳regulation of protein-containing complex assembly | 434 | 64 | 26.20 | 2.44 | + | 9.04E-06 |
| ↳regulation of cellular component biogenesis | 964 | 113 | 58.19 | 1.94 | + | 2.14E-06 |
| ↳negative regulation of protein polymerization | 73 | 22 | 4.41 | 4.99 | + | 1.17E-04 |
| ↳negative regulation of protein-containing complex assembly | 136 | 28 | 8.21 | 3.41 | + | 1.55E-03 |
| nuclear migration | 31 | 12 | 1.87 | 6.41 | + | 2.89E-02 |
| ↳organelle localization | 474 | 70 | 28.61 | 2.45 | + | 1.87E-06 |
| ↳cellular localization | 2053 | 268 | 123.93 | 2.16 | + | 9.79E-27 |
|  |  |  | 293.61 | 1.50 | + | 4.79E-15 |
| localization | 4864 | 440 |  |  |  |  |
| ↳intracellular transport | 1182 | 183 | 71.35 | 2.56 | + | 2.74E-24 |
| ↳transport | 3572 | 347 | 215.62 | 1.61 | + | 9.82E-15 |
| ↳establishment of localization | 3711 | 358 | 224.01 | 1.60 | + | 5.99E-15 |
| ↳establishment of organelle localization | 331 | 55 | 19.98 | 2.75 | + | 4.04E-06 |
| positive regulation of translation | 123 | 46 | 7.42 | 6.20 | + | 1.65E-15 |
| ↳positive regulation of cellular protein metabolic process | 1564 | 143 | 94.41 | 1.51 | + | 2.49E-02 |
| ↳positive regulation of protein metabolic process | 1673 | 151 | 100.99 | 1.50 | + | 2.31E-02 |
| ↳positive regulation of cellular amide metabolic process | 149 | 50 | 8.99 | 5.56 | + | 1.92E-15 |
| serine family amino acid metabolic process | 33 | 12 | 1.99 | 6.02 | + | 4.88E-02 |
| cortical actin cytoskeleton organization | 42 | 15 | 2.54 | 5.92 | + | 4.03E-03 |
| ↳cortical cytoskeleton organization | 59 | 18 | 3.56 | 5.05 | + | 2.07E-03 |
| ↳cytoskeleton organization | 1070 | 131 | 64.59 | 2.03 | + | 5.21E-09 |
| ↳actin cytoskeleton organization | 500 | 75 | 30.18 | 2.48 | + | 1.54E-07 |
| ↳actin filament-based process | 555 | 80 | 33.50 | 2.39 | + | 2.16E-07 |
| alpha-amino acid biosynthetic process | 51 | 18 | 3.08 | 5.85 | + | 3.36E-04 |
| ↳cellular amino acid biosynthetic process | 55 | 19 | 3.32 | 5.72 | + | 1.87E-04 |
| ↳carboxylic acid biosynthetic process | 224 | 50 | 13.52 | 3.70 | + | 1.89E-09 |
| ↳organic acid biosynthetic process | 225 | 51 | 13.58 | 3.76 | + | 6.52E-10 |
| ↳small molecule biosynthetic process | 446 | 76 | 26.92 | 2.82 | + | 3.83E-10 |
| cellular aldehyde metabolic process | 52 | 18 | 3.14 | 5.73 | + | 4.29E-04 |
| positive regulation of RNA splicing | 50 | 17 | 3.02 | 5.63 | + | 1.24E-03 |
| regulation of translational initiation | 59 | 20 | 3.56 | 5.62 | + | 1.04E-04 |
| pyruvate metabolic process | 69 | 22 | 4.17 | 5.28 | + | 4.96E-05 |
| cytoplasmic translation | 64 | 20 | 3.86 | 5.18 | + | 3.21E-04 |
| ribosome assembly | 62 | 19 | 3.74 | 5.08 | + | 9.06E-04 |
| ↳ribonucleoprotein complex biogenesis | 392 | 62 | 23.66 | 2.62 | + | 1.86E-06 |
| cofactor catabolic process | 56 | 17 | 3.38 | 5.03 | + | 4.72E-03 |
| mitochondrial translation | 52 | 15 | 3.14 | 4.78 | + | 3.69E-02 |
| ↳mitochondrial gene expression | 81 | 21 | 4.89 | 4.29 | + | 2.14E-03 |
| maintenance of protein location in cell | 77 | 22 | 4.65 | 4.73 | + | 2.61E-04 |
| ↳maintenance of location in cell | 103 | 27 | 6.22 | 4.34 | + | 3.49E-05 |
| ↳maintenance of location | 173 | 35 | 10.44 | 3.35 | + | 6.89E-05 |
| ↳cellular protein localization | 1423 | 189 | 85.90 | 2.20 | + | 3.82E-18 |
| ↳cellular macromolecule localization | 1430 | 190 | 86.32 | 2.20 | + | 2.73E-18 |
| ↳macromolecule localization | 2266 | 273 | 136.78 | 2.00 | + | 2.72E-22 |
| ↳protein localization | 1963 | 238 | 118.49 | 2.01 | + | 5.22E-19 |
| ↳maintenance of protein location | 108 | 26 | 6.52 | 3.99 | + | 2.96E-04 |
| antibiotic metabolic process | 91 | 26 | 5.49 | 4.73 | + | 1.47E-05 |
| ↳drug metabolic process | 412 | 87 | 24.87 | 3.50 | + | 4.61E-17 |
| glutathione metabolic process | 58 | 16 | 3.50 | 4.57 | + | 2.89E-02 |
| ↳cellular modified amino acid metabolic process | 156 | 34 | 9.42 | 3.61 | + | 2.21E-05 |
| protein localization to endoplasmic reticulum | 59 | 16 | 3.56 | 4.49 | + | 3.48E-02 |
| ↳protein localization to organelle | 622 | 93 | 37.55 | 2.48 | + | 6.24E-10 |
| purine ribonucleoside triphosphate metabolic process | 59 | 16 | 3.56 | 4.49 | + | 3.48E-02 |
| ↳ribonucleoside triphosphate metabolic process | 63 | 17 | 3.80 | 4.47 | + | 1.85E-02 |
| nuclear-transcribed mRNA catabolic process | 94 | 25 | 5.67 | 4.41 | + | 1.02E-04 |
| ↳mRNA catabolic process | 119 | 30 | 7.18 | 4.18 | + | 1.06E-05 |
| ↳RNA catabolic process | 147 | 32 | 8.87 | 3.61 | + | 6.63E-05 |
| ↳nucleobase-containing compound catabolic process | 234 | 37 | 14.13 | 2.62 | + | 7.63E-03 |
| ↳heterocycle catabolic process | 278 | 52 | 16.78 | 3.10 | + | 2.16E-07 |
| ↳cellular nitrogen compound catabolic process | 272 | 51 | 16.42 | 3.11 | + | 3.12E-07 |
| ↳cellular macromolecule catabolic process | 762 | 88 | 46.00 | 1.91 | + | 5.34E-04 |
| ↳macromolecule catabolic process | 858 | 101 | 51.79 | 1.95 | + | 1.96E-05 |
| ribonucleoprotein complex assembly | 163 | 43 | 9.84 | 4.37 | + | 7.69E-10 |
| ↳cellular protein-containing complex assembly | 653 | 92 | 39.42 | 2.33 | + | 2.36E-08 |
| ↳protein-containing complex assembly | 1003 | 141 | 60.54 | 2.33 | + | 1.63E-14 |
| ↳protein-containing complex subunit organization | 1144 | 156 | 69.06 | 2.26 | + | 3.49E-15 |
| ↳ribonucleoprotein complex subunit organization | 170 | 45 | 10.26 | 4.39 | + | 1.84E-10 |
| mRNA transport | 107 | 28 | 6.46 | 4.34 | + | 1.86E-05 |
| ↳RNA transport | 150 | 37 | 9.05 | 4.09 | + | 2.18E-07 |
| ↳nucleic acid transport | 150 | 37 | 9.05 | 4.09 | + | 2.18E-07 |
| ↳nucleobase-containing compound transport | 183 | 39 | 11.05 | 3.53 | + | 2.66E-06 |
| ↳organic substance transport | 1856 | 216 | 112.04 | 1.93 | + | 5.08E-15 |
| ↳nitrogen compound transport | 1550 | 190 | 93.56 | 2.03 | + | 8.20E-15 |
| ↳establishment of RNA localization | 152 | 37 | 9.18 | 4.03 | + | 3.02E-07 |
| ↳RNA localization | 170 | 42 | 10.26 | 4.09 | + | 9.48E-09 |
| carbohydrate catabolic process | 88 | 23 | 5.31 | 4.33 | + | 5.10E-04 |
| ↳carbohydrate metabolic process | 405 | 62 | 24.45 | 2.54 | + | 4.43E-06 |
| axo-dendritic transport | 73 | 18 | 4.41 | 4.08 | + | 2.80E-02 |
| ↳transport along microtubule | 155 | 29 | 9.36 | 3.10 | + | 5.38E-03 |
| ↳movement of cell or subcellular component | 1395 | 133 | 84.21 | 1.58 | + | 7.12E-03 |
| ↳cytoskeleton-dependent intracellular transport | 186 | 34 | 11.23 | 3.03 | + | 9.68E-04 |
| regulation of telomere maintenance | 81 | 19 | 4.89 | 3.89 | + | 2.82E-02 |
| protein folding | 146 | 34 | 8.81 | 3.86 | + | 5.07E-06 |
| positive regulation of viral process | 82 | 19 | 4.95 | 3.84 | + | 3.29E-02 |
| ↳regulation of viral process | 183 | 32 | 11.05 | 2.90 | + | 5.39E-03 |
| ↳regulation of symbiosis, encompassing mutualism through parasitism | 200 | 36 | 12.07 | 2.98 | + | 5.80E-04 |
| protein export from nucleus | 91 | 21 | 5.49 | 3.82 | + | 1.09E-02 |
| ↳nucleocytoplasmic transport | 203 | 36 | 12.25 | 2.94 | + | 8.04E-04 |
| ↳nuclear transport | 203 | 36 | 12.25 | 2.94 | + | 8.04E-04 |
| ↳intracellular protein transport | 765 | 109 | 46.18 | 2.36 | + | 7.39E-11 |
| ↳protein transport | 1246 | 167 | 75.21 | 2.22 | + | 6.22E-16 |
| ↳establishment of protein localization | 1330 | 179 | 80.28 | 2.23 | + | 1.78E-17 |
| ↳peptide transport | 1276 | 168 | 77.02 | 2.18 | + | 2.67E-15 |
| ↳amide transport | 1301 | 169 | 78.53 | 2.15 | + | 5.89E-15 |
| post-Golgi vesicle-mediated transport | 88 | 20 | 5.31 | 3.77 | + | 2.37E-02 |
| ↳Golgi vesicle transport | 250 | 49 | 15.09 | 3.25 | + | 1.95E-07 |
| ↳vesicle-mediated transport | 1271 | 141 | 76.72 | 1.84 | + | 4.09E-07 |
| protein stabilization | 174 | 39 | 10.50 | 3.71 | + | 7.44E-07 |
| ↳regulation of protein stability | 278 | 49 | 16.78 | 2.92 | + | 4.82E-06 |
| endoplasmic reticulum to Golgi vesicle-mediated transport | 105 | 23 | 6.34 | 3.63 | + | 7.55E-03 |
| ATP metabolic process | 170 | 37 | 10.26 | 3.61 | + | 4.57E-06 |
| regulation of viral genome replication | 93 | 20 | 5.61 | 3.56 | + | 4.89E-02 |
| drug catabolic process | 148 | 31 | 8.93 | 3.47 | + | 2.45E-04 |
| cytosolic transport | 132 | 27 | 7.97 | 3.39 | + | 2.85E-03 |
| regulation of cell shape | 158 | 31 | 9.54 | 3.25 | + | 8.88E-04 |
| ↳ regulation of cell morphogenesis | 537 | 83 | 32.42 | 2.56 | + | 3.54E-09 |
| ↳ regulation of anatomical structure morphogenesis | 1090 | 124 | 65.80 | 1.88 | + | 1.70E-06 |
| ↳ regulation of developmental process | 2684 | 239 | 162.02 | 1.48 | + | 3.13E-05 |
| receptor-mediated endocytosis | 138 | 27 | 8.33 | 3.24 | + | 6.11E-03 |
| positive regulation of endocytosis | 123 | 24 | 7.42 | 3.23 | + | 2.60E-02 |
| ↳ positive regulation of transport | 1099 | 131 | 66.34 | 1.97 | + | 2.57E-08 |
| ↳ regulation of transport | 1972 | 207 | 119.04 | 1.74 | + | 8.79E-10 |
| ↳ positive regulation of cellular component organization | 1254 | 153 | 75.70 | 2.02 | + | 4.56E-11 |
| ↳ regulation of vesicle-mediated transport | 598 | 77 | 36.10 | 2.13 | + | 5.12E-05 |
| regulation of nucleocytoplasmic transport | 125 | 24 | 7.55 | 3.18 | + | 3.32E-02 |
| ↳ regulation of intracellular transport | 352 | 59 | 21.25 | 2.78 | + | 7.42E-07 |
| monocarboxylic acid biosynthetic process | 138 | 26 | 8.33 | 3.12 | + | 1.80E-02 |
| monosaccharide metabolic process | 156 | 29 | 9.42 | 3.08 | + | 6.03E-03 |
| establishment of vesicle localization | 142 | 26 | 8.57 | 3.03 | + | 2.86E-02 |
| ↳ establishment of localization in cell | 353 | 47 | 21.31 | 2.21 | + | 2.72E-02 |
| ↳ vesicle localization | 154 | 27 | 9.30 | 2.90 | + | 3.81E-02 |
| positive regulation of intracellular transport | 198 | 36 | 11.95 | 3.01 | + | 4.65E-04 |
| protein import | 145 | 26 | 8.75 | 2.97 | + | 3.99E-02 |
| establishment of protein localization to organelle | 302 | 54 | 18.23 | 2.96 | + | 3.98E-07 |
| cellular carbohydrate metabolic process | 146 | 26 | 8.81 | 2.95 | + | 4.45E-02 |
| positive regulation of cell morphogenesis involved in differentiation | 189 | 33 | 11.41 | 2.89 | + | 3.72E-03 |
| ↳ regulation of cell morphogenesis involved in differentiation | 348 | 52 | 21.01 | 2.48 | + | 2.26E-04 |
| ↳ regulation of cell development | 1092 | 112 | 65.92 | 1.70 | + | 2.37E-03 |
| actin filament organization | 241 | 42 | 14.55 | 2.89 | + | 1.09E-04 |
| ↳ supramolecular fiber organization | 465 | 73 | 28.07 | 2.60 | + | 4.53E-08 |
| regulation of microtubule cytoskeleton organization | 203 | 35 | 12.25 | 2.86 | + | 2.19E-03 |
| ↳ regulation of microtubule-based process | 241 | 40 | 14.55 | 2.75 | + | 7.69E-04 |
| response to metal ion | 270 | 46 | 16.30 | 2.82 | + | 4.14E-05 |
| ↳ response to inorganic substance | 417 | 69 | 25.17 | 2.74 | + | 2.21E-08 |
| ↳ response to chemical | 3484 | 341 | 210.31 | 1.62 | + | 6.50E-15 |
| positive regulation of protein-containing complex assembly | 221 | 37 | 13.34 | 2.77 | + | 1.92E-03 |
| ↳ positive regulation of cellular component biogenesis | 524 | 64 | 31.63 | 2.02 | + | 6.90E-03 |
| endomembrane system organization | 367 | 61 | 22.15 | 2.75 | + | 4.05E-07 |
| protein-containing complex localization | 206 | 34 | 12.43 | 2.73 | + | 7.83E-03 |
| endosomal transport | 203 | 32 | 12.25 | 2.61 | + | 4.70E-02 |
| positive regulation of developmental growth | 216 | 34 | 13.04 | 2.61 | + | 2.36E-02 |
| ↳ positive regulation of growth | 299 | 42 | 18.05 | 2.33 | + | 2.33E-02 |
| protein complex oligomerization | 229 | 35 | 13.82 | 2.53 | + | 2.73E-02 |
| negative regulation of apoptotic signaling pathway | 232 | 35 | 14.00 | 2.50 | + | 3.46E-02 |
| ↳ negative regulation of apoptotic process | 906 | 113 | 54.69 | 2.07 | + | 8.52E-08 |
| ↳ negative regulation of programmed cell death | 926 | 114 | 55.90 | 2.04 | + | 1.39E-07 |
| ↳ regulation of programmed cell death | 1518 | 163 | 91.63 | 1.78 | + | 1.07E-07 |
| ↳ regulation of cell death | 1672 | 183 | 100.93 | 1.81 | + | 9.26E-10 |
| ↳ negative regulation of cell death | 1041 | 127 | 62.84 | 2.02 | + | 1.20E-08 |
| ↳ regulation of apoptotic process | 1493 | 161 | 90.12 | 1.79 | + | 1.09E-07 |
| positive regulation of organelle organization | 596 | 87 | 35.98 | 2.42 | + | 1.53E-08 |
| positive regulation of protein transport | 443 | 63 | 26.74 | 2.36 | + | 4.44E-05 |
| ↳ regulation of protein transport | 739 | 94 | 44.61 | 2.11 | + | 1.94E-06 |
| regulation of peptide transport | 781 | 95 | 47.14 | 2.02 | + | 1.44E-05 |
| response to oxidative stress | 337 | 47 | 20.34 | 2.31 | + | 8.96E-03 |
| response to stress | 3164 | 261 | 190.99 | 1.37 | + | 3.41E-03 |
| negative regulation of proteolysis | 345 | 46 | 20.83 | 2.21 | + | 3.64E-02 |
| regulation of proteolysis | 717 | 78 | 43.28 | 1.80 | + | 2.23E-02 |
| cell part morphogenesis | 507 | 66 | 30.60 | 2.16 | + | 6.21E-04 |
| cellular component morphogenesis | 844 | 98 | 50.95 | 1.92 | + | 5.75E-05 |
| developmental process | 5570 | 451 | 336.23 | 1.34 | + | 1.02E-07 |
| anatomical structure development | 5212 | 414 | 314.62 | 1.32 | + | 1.61E-05 |
| cellular developmental process | 3783 | 305 | 228.36 | 1.34 | + | 2.06E-03 |
| negative regulation of hydrolase activity | 401 | 51 | 24.21 | 2.11 | + | 2.67E-02 |
| negative regulation of catalytic activity | 701 | 76 | 42.32 | 1.80 | + | 3.64E-02 |
| negative regulation of molecular function | 1034 | 106 | 62.42 | 1.70 | + | 4.80E-03 |
| regulation of molecular function | 2559 | 242 | 154.47 | 1.57 | + | 1.15E-07 |
| regulation of catalytic activity | 1881 | 177 | 113.54 | 1.56 | + | 1.84E-04 |
| regulation of hydrolase activity | 994 | 118 | 60.00 | 1.97 | + | 4.18E-07 |
| cell junction organization | 466 | 56 | 28.13 | 1.99 | + | 4.78E-02 |
| cellular response to nitrogen compound | 512 | 61 | 30.91 | 1.97 | + | 2.21E-02 |
| response to nitrogen compound | 851 | 99 | 51.37 | 1.93 | + | 4.32E-05 |
| cellular response to chemical stimulus | 2359 | 234 | 142.40 | 1.64 | + | 2.99E-09 |
| neuron projection development | 691 | 82 | 41.71 | 1.97 | + | 4.84E-04 |
| neuron development | 843 | 89 | 50.89 | 1.75 | + | 1.43E-02 |
| generation of neurons | 1663 | 154 | 100.38 | 1.53 | + | 4.38E-03 |
| neurogenesis | 1771 | 161 | 106.90 | 1.51 | + | 6.92E-03 |
| cell differentiation | 3696 | 293 | 223.10 | 1.31 | + | 1.63E-02 |
| system development | 4226 | 338 | 255.10 | 1.32 | + | 6.50E-04 |
| multicellular organism development | 4844 | 384 | 292.40 | 1.31 | + | 1.32E-04 |
| cell development | 1750 | 167 | 105.64 | 1.58 | + | 2.08E-04 |
| plasma membrane bounded cell projection organization | 1060 | 106 | 63.99 | 1.66 | + | 1.68E-02 |
| cell morphogenesis | 736 | 87 | 44.43 | 1.96 | + | 2.12E-04 |
| response to organic cyclic compound | 621 | 72 | 37.49 | 1.92 | + | 9.15E-03 |
| response to organic substance | 2488 | 251 | 150.19 | 1.67 | + | 6.41E-11 |
| response to organonitrogen compound | 745 | 85 | 44.97 | 1.89 | + | 1.39E-03 |
| response to hormone | 611 | 69 | 36.88 | 1.87 | + | 2.77E-02 |
| response to endogenous stimulus | 1126 | 118 | 67.97 | 1.74 | + | 3.27E-04 |
| regulation of plasma membrane bounded cell projection organization | 781 | 87 | 47.14 | 1.85 | + | 2.59E-03 |
| regulation of cell projection organization | 790 | 87 | 47.69 | 1.82 | + | 4.42E-03 |
| cellular response to cytokine stimulus | 639 | 71 | 38.57 | 1.84 | + | 4.30E-02 |
| cellular response to organic substance | 1822 | 173 | 109.98 | 1.57 | + | 1.54E-04 |
| response to cytokine | 746 | 86 | 45.03 | 1.91 | + | 9.26E-04 |
| cellular homeostasis | 857 | 93 | 51.73 | 1.80 | + | 2.72E-03 |
| homeostatic process | 1644 | 155 | 99.24 | 1.56 | + | 1.63E-03 |
| cellular response to oxygen-containing compound | 912 | 95 | 55.05 | 1.73 | + | 1.18E-02 |
| response to oxygen-containing compound | 1296 | 131 | 78.23 | 1.67 | + | 4.24E-04 |
| regulation of cell migration | 914 | 93 | 55.17 | 1.69 | + | 3.46E-02 |
| regulation of cell motility | 965 | 99 | 58.25 | 1.70 | + | 1.16E-02 |
| regulation of locomotion | 1042 | 112 | 62.90 | 1.78 | + | 2.17E-04 |
| regulation of cellular component movement | 1054 | 116 | 63.62 | 1.82 | + | 3.70E-05 |
| regulation of response to stress | 1295 | 124 | 78.17 | 1.59 | + | 1.35E-02 |
| cellular response to stress | 1446 | 137 | 87.29 | 1.57 | + | 6.15E-03 |
| positive regulation of molecular function | 1477 | 139 | 89.16 | 1.56 | + | 7.72E-03 |
| regulation of multicellular organismal process | 3223 | 278 | 194.55 | 1.43 | + | 2.39E-05 |
| Unclassified | 1901 | 34 | 114.75 | .30 | - | 0.00E00 |
| G protein-coupled receptor signaling pathway | 1851 | 18 | 111.73 | .16 | - | 2.61E-24 |
| ↳ signal transduction | 4827 | 197 | 291.38 | .68 | - | 2.14E-06 |
| ↳ cell communication | 5260 | 235 | 317.51 | .74 | - | 1.06E-03 |
| ↳ signaling | 5138 | 218 | 310.15 | .70 | - | 1.58E-05 |
| positive regulation of B cell activation | 269 | 1 | 16.24 | .06 | - | 3.53E-02 |
| sensory perception of chemical stimulus | 1228 | 1 | 74.13 | .01 | - | 7.81E-27 |
| ↳ sensory perception | 1642 | 17 | 99.12 | .17 | - | 2.51E-20 |
| ↳ nervous system process | 2084 | 55 | 125.80 | .44 | - | 7.65E-09 |

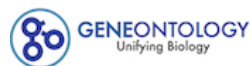

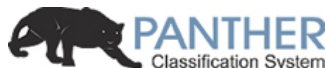

PANTHER 15.0 released!

Analysis Summary: Please report in publication ?

Analysis Type: PANTHER Overrepresentation Test (Released 20200407)

Annotation Version and Release Date: GO Ontology database Released 2020-02-21

Analyzed List: upload\_1 (Mus musculus) Change

Reference List: Mus musculus (all genes in database) Change

Annotation Data Set: GO molecular function complete ?

Test Type: ☒ Fisher's Exact ☐ Binomial

Correction: ☐ Calculate False Discovery Rate ☒ Use the Bonferroni correction for multiple testing ? ☐ No correction

Results ?

|  | Reference list | upload_1 |
| --- | --- | --- |
| Uniquely Mapped IDs: | 22265 out of 22265 | 1344 out of 1344 |
| Unmapped IDs: | 0 | 0 |
| Multiple mapping information: | 0 | 0 |

Bonferroni count: 2756

Export Table XML with user input ids JSON with user input ids

Displaying only results for Bonferroni-corrected for P < 0.05, click here to display all results

|  | Mus musculus (REF) | upload_1 (Hierarchy NEW! ?) |  |  |  |  |
| --- | --- | --- | --- | --- | --- | --- |
| GO molecular function complete | # | # | expected | Fold Enrichment | +/- | P value |
| ligase activity, forming carbon-carbon bonds | 7 | 7 | .42 | 16.57 | + | 1.27E-02 |
| ↳ ligase activity | 146 | 57 | 8.81 | 6.47 | + | 4.26E-21 |
| ↳ catalytic activity | 5659 | 564 | 341.60 | 1.65 | + | 6.60E-34 |
| N6-methyladenosine-containing RNA binding | 8 | 7 | .48 | 14.50 | + | 2.26E-02 |
| ↳ RNA binding | 1086 | 338 | 65.56 | 5.16 | + | 4.40E-121 |
| ↳ nucleic acid binding | 3161 | 389 | 190.81 | 2.04 | + | 1.15E-37 |
| ↳ organic cyclic compound binding | 5175 | 660 | 312.38 | 2.11 | + | 7.59E-85 |
| ↳ binding | 13351 | 1179 | 805.92 | 1.46 | + | 2.91E-103 |
| ↳ heterocyclic compound binding | 5072 | 650 | 306.17 | 2.12 | + | 1.15E-83 |
| acyl-CoA dehydrogenase activity | 10 | 8 | .60 | 13.25 | + | 7.79E-03 |
| ↳ oxidoreductase activity, acting on the CH-CH group of donors | 56 | 16 | 3.38 | 4.73 | + | 6.11E-03 |
| ↳ oxidoreductase activity | 791 | 117 | 47.75 | 2.45 | + | 2.11E-13 |
| RNA stem-loop binding | 14 | 10 | .85 | 11.83 | + | 8.96E-04 |
| translation elongation factor activity | 17 | 11 | 1.03 | 10.72 | + | 4.70E-04 |
| ↳ translation factor activity, RNA binding | 77 | 39 | 4.65 | 8.39 | + | 7.46E-17 |
| ↳ translation regulator activity, nucleic acid binding | 95 | 45 | 5.73 | 7.85 | + | 7.47E-19 |
| ↳ translation regulator activity | 126 | 60 | 7.61 | 7.89 | + | 6.02E-26 |
| mRNA 5'-UTR binding | 24 | 14 | 1.45 | 9.66 | + | 2.57E-05 |
| ↳ mRNA binding | 272 | 120 | 16.42 | 7.31 | + | 1.08E-51 |
| miRNA binding | 30 | 17 | 1.81 | 9.39 | + | 9.28E-07 |
| ↳ regulatory RNA binding | 43 | 18 | 2.60 | 6.93 | + | 1.23E-05 |
| acid-thiol ligase activity | 27 | 15 | 1.63 | 9.20 | + | 1.25E-05 |
| ligase activity, forming carbon-sulfur bonds | 38 | 17 | 2.29 | 7.41 | + | 1.50E-05 |
| CoA-ligase activity | 24 | 12 | 1.45 | 8.28 | + | 1.06E-03 |
| aminoacyl-tRNA ligase activity | 41 | 20 | 2.47 | 8.08 | + | 2.03E-07 |
| ligase activity, forming carbon-oxygen bonds | 41 | 20 | 2.47 | 8.08 | + | 2.03E-07 |
| catalytic activity, acting on RNA | 337 | 63 | 20.34 | 3.10 | + | 5.28E-10 |
| translation initiation factor activity | 48 | 23 | 2.90 | 7.94 | + | 1.02E-08 |
| C-acyltransferase activity | 21 | 10 | 1.27 | 7.89 | + | 1.40E-02 |
| RNA helicase activity | 53 | 25 | 3.20 | 7.81 | + | 1.56E-09 |
| helicase activity | 148 | 35 | 8.93 | 3.92 | + | 6.18E-07 |
| ATPase activity, coupled | 279 | 51 | 16.84 | 3.03 | + | 2.15E-07 |
| ATPase activity | 412 | 69 | 24.87 | 2.77 | + | 4.83E-09 |
| nucleoside-triphosphatase activity | 748 | 112 | 45.15 | 2.48 | + | 4.36E-13 |
| pyrophosphatase activity | 801 | 117 | 48.35 | 2.42 | + | 6.11E-13 |
| hydrolase activity, acting on acid anhydrides, in phosphorus-containing anhydrides | 804 | 117 | 48.53 | 2.41 | + | 6.87E-13 |
| hydrolase activity, acting on acid anhydrides | 804 | 117 | 48.53 | 2.41 | + | 6.87E-13 |
| hydrolase activity | 2449 | 242 | 147.83 | 1.64 | + | 5.67E-10 |
| double-stranded RNA binding | 82 | 38 | 4.95 | 7.68 | + | 2.40E-15 |
| poly(A) binding | 24 | 11 | 1.45 | 7.59 | + | 6.32E-03 |
| poly-purine tract binding | 32 | 16 | 1.93 | 8.28 | + | 1.22E-05 |
| single-stranded RNA binding | 89 | 38 | 5.37 | 7.07 | + | 2.11E-14 |
| oxidoreductase activity, acting on the CH-NH group of donors | 27 | 12 | 1.63 | 7.36 | + | 2.83E-03 |
| mRNA 3'-UTR binding | 87 | 38 | 5.25 | 7.24 | + | 1.15E-14 |
| NAD binding | 61 | 26 | 3.68 | 7.06 | + | 3.37E-09 |
| coenzyme binding | 288 | 82 | 17.38 | 4.72 | + | 1.56E-23 |
| cofactor binding | 531 | 108 | 32.05 | 3.37 | + | 2.76E-21 |
| nucleotide binding | 2030 | 316 | 122.54 | 2.58 | + | 1.32E-47 |
| small molecule binding | 2421 | 370 | 146.14 | 2.53 | + | 8.96E-56 |
| nucleoside phosphate binding | 2030 | 316 | 122.54 | 2.58 | + | 1.32E-47 |
| ADP binding | 40 | 17 | 2.41 | 7.04 | + | 2.75E-05 |
| adenyl ribonucleotide binding | 1453 | 215 | 87.71 | 2.45 | + | 1.66E-27 |
| adenyl nucleotide binding | 1464 | 219 | 88.37 | 2.48 | + | 1.55E-28 |
| purine nucleotide binding | 1798 | 264 | 108.53 | 2.43 | + | 1.79E-34 |
| purine ribonucleotide binding | 1786 | 260 | 107.81 | 2.41 | + | 3.02E-33 |
| ribonucleotide binding | 1802 | 263 | 108.78 | 2.42 | + | 7.80E-34 |
| carbohydrate derivative binding | 2123 | 294 | 128.15 | 2.29 | + | 7.95E-35 |
| anion binding | 2703 | 391 | 163.16 | 2.40 | + | 6.47E-54 |
| ion binding | 5485 | 575 | 331.10 | 1.74 | + | 1.93E-41 |
| poly(U) RNA binding | 26 | 11 | 1.57 | 7.01 | + | 1.17E-02 |
| poly-pyrimidine tract binding | 30 | 13 | 1.81 | 7.18 | + | 1.26E-03 |
| ribosome binding | 64 | 27 | 3.86 | 6.99 | + | 1.51E-09 |
| ribonucleoprotein complex binding | 146 | 46 | 8.81 | 5.22 | + | 1.22E-13 |
| protein-containing complex binding | 1451 | 252 | 87.59 | 2.88 | + | 7.63E-44 |
| rRNA binding | 70 | 28 | 4.23 | 6.63 | + | 1.60E-09 |
| pre-mRNA binding | 33 | 13 | 1.99 | 6.53 | + | 2.99E-03 |
| oxidoreductase activity, acting on the aldehyde or oxo group of donors, NAD or NADP as acceptor | 47 | 18 | 2.84 | 6.34 | + | 3.74E-05 |
| oxidoreductase activity, acting on the aldehyde or oxo group of donors | 56 | 19 | 3.38 | 5.62 | + | 7.37E-05 |
| Ran GTPase binding | 34 | 12 | 2.05 | 5.85 | + | 1.94E-02 |
| Ras GTPase binding | 399 | 57 | 24.09 | 2.37 | + | 7.94E-05 |
| small GTPase binding | 416 | 59 | 25.11 | 2.35 | + | 4.92E-05 |
| GTPase binding | 515 | 73 | 31.09 | 2.35 | + | 9.78E-07 |
| enzyme binding | 2331 | 306 | 140.71 | 2.17 | + | 1.43E-32 |
| protein binding | 9112 | 865 | 550.03 | 1.57 | + | 1.27E-59 |
| extracellular matrix structural constituent | 143 | 48 | 8.63 | 5.56 | + | 3.10E-15 |
| structural molecule activity | 576 | 151 | 34.77 | 4.34 | + | 4.40E-42 |
| structural constituent of cytoskeleton | 64 | 21 | 3.86 | 5.44 | + | 2.24E-05 |
| hydro-lyase activity | 52 | 17 | 3.14 | 5.42 | + | 6.12E-04 |
| carbon-oxygen lyase activity | 62 | 18 | 3.74 | 4.81 | + | 1.18E-03 |
| lyase activity | 184 | 38 | 11.11 | 3.42 | + | 2.99E-06 |
| flavin adenine dinucleotide binding | 82 | 26 | 4.95 | 5.25 | + | 7.11E-07 |
| structural constituent of ribosome | 161 | 49 | 9.72 | 5.04 | + | 3.73E-14 |
| unfolded protein binding | 92 | 28 | 5.55 | 5.04 | + | 3.24E-07 |
| NADP binding | 47 | 14 | 2.84 | 4.93 | + | 1.75E-02 |
| amino acid binding | 71 | 21 | 4.29 | 4.90 | + | 1.01E-04 |
| carboxylic acid binding | 223 | 45 | 13.46 | 3.34 | + | 1.73E-07 |
| organic acid binding | 237 | 46 | 14.31 | 3.22 | + | 3.28E-07 |
| tRNA binding | 60 | 17 | 3.62 | 4.69 | + | 3.27E-03 |
| actin filament binding | 201 | 56 | 12.13 | 4.62 | + | 5.27E-15 |
| actin binding | 426 | 93 | 25.71 | 3.62 | + | 8.01E-20 |
| cytoskeletal protein binding | 976 | 163 | 58.92 | 2.77 | + | 9.40E-25 |
| pyridoxal phosphate binding | 51 | 14 | 3.08 | 4.55 | + | 3.82E-02 |
| vitamin B6 binding | 52 | 14 | 3.14 | 4.46 | + | 4.60E-02 |
| drug binding | 1644 | 242 | 99.24 | 2.44 | + | 3.15E-31 |
| vitamin binding | 132 | 27 | 7.97 | 3.39 | + | 8.84E-04 |
| modified amino acid binding | 100 | 24 | 6.04 | 3.98 | + | 3.22E-04 |
| heat shock protein binding | 140 | 33 | 8.45 | 3.90 | + | 2.16E-06 |
| chaperone binding | 101 | 22 | 6.10 | 3.61 | + | 4.36E-03 |
| single-stranded DNA binding | 111 | 23 | 6.70 | 3.43 | + | 5.34E-03 |
| oxidoreductase activity, acting on the CH-OH group of donors, NAD or NADP as acceptor | 141 | 28 | 8.51 | 3.29 | + | 9.11E-04 |
| oxidoreductase activity, acting on CH-OH group of donors | 149 | 30 | 8.99 | 3.34 | + | 2.69E-04 |
| integrin binding | 133 | 26 | 8.03 | 3.24 | + | 3.07E-03 |
| cell adhesion molecule binding | 245 | 39 | 14.79 | 2.64 | + | 1.04E-03 |
| ion channel binding | 140 | 25 | 8.45 | 2.96 | + | 2.00E-02 |
| protein C-terminus binding | 231 | 39 | 13.94 | 2.80 | + | 2.32E-04 |
| magnesium ion binding | 197 | 32 | 11.89 | 2.69 | + | 1.06E-02 |
| metal ion binding | 3499 | 299 | 211.21 | 1.42 | + | 3.71E-06 |
| cation binding | 3598 | 311 | 217.19 | 1.43 | + | 4.02E-07 |
| ubiquitin protein ligase binding | 308 | 48 | 18.59 | 2.58 | + | 1.19E-04 |
| ubiquitin-like protein ligase binding | 323 | 50 | 19.50 | 2.56 | + | 6.29E-05 |
| GTP binding | 355 | 55 | 21.43 | 2.57 | + | 1.06E-05 |
| purine ribonucleoside triphosphate binding | 1713 | 247 | 103.40 | 2.39 | + | 9.31E-31 |
| purine ribonucleoside binding | 362 | 57 | 21.85 | 2.61 | + | 3.87E-06 |
| ribonucleoside binding | 365 | 58 | 22.03 | 2.63 | + | 1.40E-06 |
| nucleoside binding | 375 | 59 | 22.64 | 2.61 | + | 1.89E-06 |
| purine nucleoside binding | 366 | 57 | 22.09 | 2.58 | + | 4.73E-06 |
| guanyl ribonucleotide binding | 379 | 58 | 22.88 | 2.54 | + | 5.21E-06 |
| guanyl nucleotide binding | 379 | 58 | 22.88 | 2.54 | + | 5.21E-06 |
| microtubule binding | 247 | 38 | 14.91 | 2.55 | + | 2.79E-03 |
| tubulin binding | 346 | 49 | 20.89 | 2.35 | + | 7.74E-04 |
| sulfur compound binding | 255 | 38 | 15.39 | 2.47 | + | 6.87E-03 |
| amide binding | 378 | 56 | 22.82 | 2.45 | + | 3.26E-05 |
| phospholipid binding | 427 | 63 | 25.78 | 2.44 | + | 3.77E-06 |
| lipid binding | 763 | 92 | 46.06 | 2.00 | + | 1.19E-05 |
| ATP binding | 1390 | 203 | 83.91 | 2.42 | + | 5.34E-25 |
| protein kinase binding | 727 | 93 | 43.88 | 2.12 | + | 7.15E-07 |
| ↳ kinase binding | 813 | 99 | 49.08 | 2.02 | + | 1.52E-06 |
| protein domain specific binding | 786 | 100 | 47.45 | 2.11 | + | 1.69E-07 |
| protein homodimerization activity | 648 | 75 | 39.12 | 1.92 | + | 1.36E-03 |
| ↳ identical protein binding | 1983 | 271 | 119.70 | 2.26 | + | 1.01E-30 |
| ↳ protein dimerization activity | 945 | 99 | 57.04 | 1.74 | + | 1.43E-03 |
| calcium ion binding | 612 | 67 | 36.94 | 1.81 | + | 2.97E-02 |
| enzyme regulator activity | 989 | 98 | 59.70 | 1.64 | + | 1.72E-02 |
| Unclassified | 2116 | 37 | 127.73 | .29 | - | 0.00E00 |
| receptor ligand activity | 488 | 6 | 29.46 | .20 | - | 1.61E-03 |
| ↳ signaling receptor activator activity | 493 | 6 | 29.76 | .20 | - | 1.12E-03 |
| ↳ receptor regulator activity | 534 | 8 | 32.23 | .25 | - | 3.28E-03 |
| DNA-binding transcription activator activity, RNA polymerase II-specific | 472 | 5 | 28.49 | .18 | - | 5.96E-04 |
| ↳ DNA-binding transcription activator activity | 475 | 5 | 28.67 | .17 | - | 6.18E-04 |
| ↳ DNA-binding transcription factor activity | 1066 | 20 | 64.35 | .31 | - | 7.44E-07 |
| ↳ transcription regulator activity | 1445 | 49 | 87.23 | .56 | - | 2.96E-02 |
| ↳ DNA-binding transcription factor activity, RNA polymerase II-specific | 814 | 13 | 49.14 | .26 | - | 6.78E-06 |
| odorant binding | 473 | 2 | 28.55 | .07 | - | 1.74E-06 |
| G protein-coupled receptor activity | 750 | 1 | 45.27 | .02 | - | 7.86E-15 |
| ↳ transmembrane signaling receptor activity | 2139 | 5 | 129.12 | .04 | - | 1.49E-45 |
| ↳ signaling receptor activity | 2339 | 13 | 141.19 | .09 | - | 2.23E-41 |
| ↳ molecular transducer activity | 2344 | 13 | 141.49 | .09 | - | 1.45E-41 |

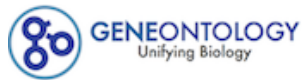

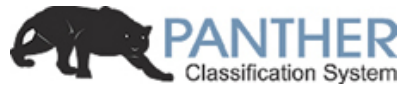

**PANTHER 15.0 released!**Analysis Summary: Please report in publication ?**Analysis Type:** PANTHER Overrepresentation Test (Released 20200407)**Annotation Version and Release Date:** GO Ontology database Released 2020-02-21**Analyzed List:** upload\_1 (Mus musculus)Change**Reference List:** Mus musculus (all genes in database)Change**Annotation Data Set:** GO cellular component complete ?**Test Type:** ☒ Fisher's Exact ☐ Binomial**Correction:** ☐ Calculate False Discovery Rate ☒ Use the Bonferroni correction for multiple testing ? ☐ No correction**Results** ?

|  | Reference list | upload_1 |
| --- | --- | --- |
| Uniquely Mapped IDs: | 22265 out of 22265 | 1344 out of 1344 |
| Unmapped IDs: | 0 | 0 |
| Multiple mapping information: | 0 | 0 |

Bonferroni count: 1452

Export Table XML with user input ids JSON with user input ids

Displaying only results for Bonferroni-corrected for P < 0.05, click here to display all results

|  | Mus musculus (REF) | upload_1 (▼ Hierarchy **NEW!** ?) |  |  |  |  |
| --- | --- | --- | --- | --- | --- | --- |
| GO cellular component complete | # | # | expected | Fold Enrichment | +/- | P value |
| eukaryotic translation initiation factor 3 complex, eIF3m | 8 | 7 | .48 | 14.50 | + | 1.19E-02 |
| ↳ eukaryotic translation initiation factor 3 complex | 15 | 12 | .91 | 13.25 | + | 1.25E-05 |
| ↳ cytoplasm | 10945 | 1140 | 660.68 | 1.73 | + | 2.25E-153 |
| ↳ cellular anatomical entity | 18638 | 1300 | 1125.06 | 1.16 | + | 3.10E-47 |
| ↳ intracellular | 13691 | 1234 | 826.44 | 1.49 | + | 4.05E-133 |
| ↳ protein-containing complex | 5309 | 630 | 320.47 | 1.97 | + | 2.03E-67 |
| chaperonin-containing T-complex | 10 | 8 | .60 | 13.25 | + | 4.11E-03 |
| ↳ chaperone complex | 22 | 14 | 1.33 | 10.54 | + | 5.93E-06 |
| ↳ cytosol | 3533 | 541 | 213.27 | 2.54 | + | 4.05E-91 |
| aminoacyl-tRNA synthetase multienzyme complex | 12 | 9 | .72 | 12.42 | + | 1.40E-03 |
| zona pellucida receptor complex | 13 | 9 | .78 | 11.47 | + | 2.25E-03 |
| endoplasmic reticulum chaperone complex | 12 | 8 | .72 | 11.04 | + | 1.07E-02 |
| ↪endoplasmic reticulum | 1639 | 187 | 98.94 | 1.89 | + | 2.08E-12 |
| ↪endomembrane system | 3884 | 380 | 234.45 | 1.62 | + | 4.47E-18 |
| ↪intracellular membrane-bounded organelle | 10291 | 974 | 621.20 | 1.57 | + | 7.37E-77 |
| ↪intracellular organelle | 11954 | 1102 | 721.59 | 1.53 | + | 7.78E-97 |
| ↪organelle | 12303 | 1111 | 742.66 | 1.50 | + | 4.55E-92 |
| ↪membrane-bounded organelle | 11048 | 1014 | 666.90 | 1.52 | + | 6.09E-76 |
| messenger ribonucleoprotein complex | 13 | 8 | .78 | 10.19 | + | 1.64E-02 |
| ↪ribonucleoprotein complex | 703 | 185 | 42.44 | 4.36 | + | 4.73E-53 |
| proton-transporting two-sector ATPase complex, catalytic domain | 14 | 8 | .85 | 9.47 | + | 2.45E-02 |
| cytoplasmic stress granule | 66 | 35 | 3.98 | 8.79 | + | 1.12E-15 |
| ↪cytoplasmic ribonucleoprotein granule | 206 | 73 | 12.43 | 5.87 | + | 8.90E-26 |
| ↪ribonucleoprotein granule | 217 | 80 | 13.10 | 6.11 | + | 2.06E-29 |
| ↪intracellular non-membrane-bounded organelle | 4162 | 477 | 251.23 | 1.90 | + | 1.72E-41 |
| ↪non-membrane-bounded organelle | 4181 | 479 | 252.38 | 1.90 | + | 1.32E-41 |
| ↪supramolecular complex | 1197 | 211 | 72.26 | 2.92 | + | 6.85E-37 |
| postsynaptic cytoskeleton | 19 | 10 | 1.15 | 8.72 | + | 3.71E-03 |
| ↪postsynapse | 727 | 120 | 43.88 | 2.73 | + | 2.57E-17 |
| ↪synapse | 1435 | 225 | 86.62 | 2.60 | + | 1.42E-32 |
| ↪cell junction | 2028 | 284 | 122.42 | 2.32 | + | 1.95E-34 |
| ↪cytoskeleton | 2135 | 241 | 128.88 | 1.87 | + | 1.70E-16 |
| smooth endoplasmic reticulum | 36 | 18 | 2.17 | 8.28 | + | 6.98E-07 |
| mitochondrial nucleoid | 47 | 22 | 2.84 | 7.75 | + | 2.27E-08 |
| ↪mitochondrial matrix | 272 | 91 | 16.42 | 5.54 | + | 4.42E-31 |
| ↪intracellular organelle lumen | 4311 | 467 | 260.23 | 1.79 | + | 2.68E-34 |
| ↪organelle lumen | 4312 | 467 | 260.29 | 1.79 | + | 3.97E-34 |
| ↪membrane-enclosed lumen | 4312 | 467 | 260.29 | 1.79 | + | 3.97E-34 |
| ↪mitochondrion | 1803 | 321 | 108.84 | 2.95 | + | 1.19E-60 |
| ↪nucleoid | 47 | 22 | 2.84 | 7.75 | + | 2.27E-08 |
| polysomal ribosome | 30 | 14 | 1.81 | 7.73 | + | 1.17E-04 |
| ↪polysome | 73 | 35 | 4.41 | 7.94 | + | 1.30E-14 |
| ↪ribosome | 234 | 70 | 14.13 | 4.96 | + | 4.65E-21 |
| postsynaptic cytosol | 26 | 12 | 1.57 | 7.65 | + | 1.09E-03 |
| ↪region of cytosol | 40 | 14 | 2.41 | 5.80 | + | 1.93E-03 |
| myelin sheath | 213 | 96 | 12.86 | 7.47 | + | 1.20E-41 |
| cytosolic small ribosomal subunit | 46 | 20 | 2.78 | 7.20 | + | 5.30E-07 |
| ↪small ribosomal subunit | 78 | 31 | 4.71 | 6.58 | + | 5.59E-11 |
| ↪ribosomal subunit | 202 | 60 | 12.19 | 4.92 | + | 1.20E-17 |
| ↪cytosolic ribosome | 117 | 37 | 7.06 | 5.24 | + | 6.87E-11 |
| proteasome accessory complex | 26 | 10 | 1.57 | 6.37 | + | 3.27E-02 |
| ↪proteasome complex | 66 | 19 | 3.98 | 4.77 | + | 3.34E-04 |
| ↪endopeptidase complex | 67 | 19 | 4.04 | 4.70 | + | 4.06E-04 |
| ↪peptidase complex | 93 | 20 | 5.61 | 3.56 | + | 7.98E-03 |
| ↳catalytic complex | 1341 | 122 | 80.95 | 1.51 | + | 2.84E-02 |
| intercalated disc | 62 | 21 | 3.74 | 5.61 | + | 7.44E-06 |
| ↳cell-cell contact zone | 85 | 23 | 5.13 | 4.48 | + | 4.86E-05 |
| ↳cell-cell junction | 497 | 58 | 30.00 | 1.93 | + | 1.47E-02 |
| ↳anchoring junction | 619 | 80 | 37.37 | 2.14 | + | 3.41E-06 |
| neuron projection cytoplasm | 39 | 13 | 2.35 | 5.52 | + | 7.14E-03 |
| ↳cytoplasmic region | 216 | 32 | 13.04 | 2.45 | + | 2.99E-02 |
| ↳plasma membrane bounded cell projection | 2281 | 252 | 137.69 | 1.83 | + | 2.37E-16 |
| ↳cell projection | 2499 | 275 | 150.85 | 1.82 | + | 5.42E-18 |
| ↳neuron projection | 1516 | 184 | 91.51 | 2.01 | + | 1.20E-14 |
| extracellular exosome | 80 | 25 | 4.83 | 5.18 | + | 1.06E-06 |
| ↳extracellular vesicle | 88 | 25 | 5.31 | 4.71 | + | 5.43E-06 |
| ↳extracellular organelle | 105 | 25 | 6.34 | 3.94 | + | 1.06E-04 |
| ↳vesicle | 2039 | 196 | 123.08 | 1.59 | + | 9.41E-07 |
| stress fiber | 85 | 26 | 5.13 | 5.07 | + | 7.14E-07 |
| ↳contractile actin filament bundle | 85 | 26 | 5.13 | 5.07 | + | 7.14E-07 |
| ↳actin filament bundle | 94 | 31 | 5.67 | 5.46 | + | 3.13E-09 |
| ↳actin cytoskeleton | 497 | 88 | 30.00 | 2.93 | + | 7.49E-14 |
| ↳actomyosin | 96 | 31 | 5.79 | 5.35 | + | 4.92E-09 |
| P-body | 72 | 22 | 4.35 | 5.06 | + | 1.54E-05 |
| catalytic step 2 spliceosome | 83 | 25 | 5.01 | 4.99 | + | 2.00E-06 |
| ↳spliceosomal complex | 197 | 49 | 11.89 | 4.12 | + | 1.62E-11 |
| ↳nucleus | 6824 | 609 | 411.92 | 1.48 | + | 1.54E-24 |
| basement membrane | 109 | 32 | 6.58 | 4.86 | + | 1.80E-08 |
| ↳collagen-containing extracellular matrix | 366 | 81 | 22.09 | 3.67 | + | 1.50E-17 |
| ↳extracellular matrix | 481 | 83 | 29.03 | 2.86 | + | 2.18E-12 |
| vesicle coat | 53 | 15 | 3.20 | 4.69 | + | 7.32E-03 |
| ↳coated vesicle membrane | 76 | 18 | 4.59 | 3.92 | + | 7.40E-03 |
| ↳vesicle membrane | 366 | 45 | 22.09 | 2.04 | + | 4.89E-02 |
| ↳organelle membrane | 2118 | 246 | 127.85 | 1.92 | + | 2.19E-18 |
| ↳cytoplasmic vesicle | 1905 | 172 | 114.99 | 1.50 | + | 7.13E-04 |
| ↳intracellular vesicle | 1910 | 172 | 115.29 | 1.49 | + | 7.39E-04 |
| ↳whole membrane | 1152 | 143 | 69.54 | 2.06 | + | 1.92E-11 |
| ↳coated vesicle | 188 | 33 | 11.35 | 2.91 | + | 5.47E-04 |
| ↳bounding membrane of organelle | 1098 | 115 | 66.28 | 1.74 | + | 1.03E-04 |
| ↳membrane coat | 95 | 20 | 5.73 | 3.49 | + | 1.05E-02 |
| ↳coated membrane | 95 | 20 | 5.73 | 3.49 | + | 1.05E-02 |
| filopodium | 93 | 25 | 5.61 | 4.45 | + | 1.39E-05 |
| ↳actin-based cell projection | 216 | 43 | 13.04 | 3.30 | + | 3.49E-07 |
| cytosolic large ribosomal subunit | 68 | 18 | 4.10 | 4.39 | + | 1.93E-03 |
| ↳large ribosomal subunit | 129 | 31 | 7.79 | 3.98 | + | 2.48E-06 |
| nuclear matrix | 88 | 22 | 5.31 | 4.14 | + | 3.10E-04 |
| ↳nuclear periphery | 113 | 27 | 6.82 | 3.96 | + | 3.03E-05 |
| ↳nuclear lumen | 3899 | 358 | 235.36 | 1.52 | + | 1.23E-12 |
| mitochondrial ribosome | 90 | 22 | 5.43 | 4.05 | + | 4.32E-04 |
| ↳organellar ribosome | 90 | 22 | 5.43 | 4.05 | + | 4.32E-04 |
| T-tubule | 70 | 17 | 4.23 | 4.02 | + | 1.01E-02 |
| ↳sarcolemma | 162 | 37 | 9.78 | 3.78 | + | 2.34E-07 |
| cortical actin cytoskeleton | 99 | 24 | 5.98 | 4.02 | + | 1.45E-04 |
| ↳cortical cytoskeleton | 130 | 31 | 7.85 | 3.95 | + | 2.90E-06 |
| ↳cell cortex | 316 | 64 | 19.07 | 3.36 | + | 7.16E-12 |
| clathrin-coated pit | 67 | 16 | 4.04 | 3.96 | + | 2.24E-02 |
| ↳plasma membrane region | 1218 | 135 | 73.52 | 1.84 | + | 2.08E-07 |
| endoplasmic reticulum-Golgi intermediate compartment | 72 | 17 | 4.35 | 3.91 | + | 1.39E-02 |
| growth cone | 205 | 48 | 12.37 | 3.88 | + | 2.07E-10 |
| ↳site of polarized growth | 213 | 49 | 12.86 | 3.81 | + | 2.04E-10 |
| ↳distal axon | 379 | 60 | 22.88 | 2.62 | + | 6.23E-07 |
| ↳axon | 715 | 103 | 43.16 | 2.39 | + | 3.64E-11 |
| focal adhesion | 156 | 36 | 9.42 | 3.82 | + | 3.22E-07 |
| ↳cell-substrate junction | 168 | 38 | 10.14 | 3.75 | + | 1.69E-07 |
| ruffle | 149 | 33 | 8.99 | 3.67 | + | 4.38E-06 |
| ↳cell leading edge | 389 | 73 | 23.48 | 3.11 | + | 2.75E-12 |
| brush border | 137 | 30 | 8.27 | 3.63 | + | 2.82E-05 |
| ↳cluster of actin-based cell projections | 195 | 35 | 11.77 | 2.97 | + | 1.53E-04 |
| microvillus | 96 | 20 | 5.79 | 3.45 | + | 1.20E-02 |
| peroxisome | 144 | 30 | 8.69 | 3.45 | + | 7.39E-05 |
| ↳microbody | 144 | 30 | 8.69 | 3.45 | + | 7.39E-05 |
| U2-type spliceosomal complex | 87 | 18 | 5.25 | 3.43 | + | 3.63E-02 |
| mitochondrial protein complex | 260 | 52 | 15.69 | 3.31 | + | 4.00E-09 |
| lamellipodium | 166 | 33 | 10.02 | 3.29 | + | 4.31E-05 |
| plasma membrane raft | 127 | 23 | 7.67 | 3.00 | + | 1.97E-02 |
| ↳membrane raft | 372 | 56 | 22.46 | 2.49 | + | 8.43E-06 |
| ↳membrane microdomain | 373 | 57 | 22.52 | 2.53 | + | 3.98E-06 |
| ↳membrane region | 386 | 60 | 23.30 | 2.58 | + | 9.13E-07 |
| postsynaptic density | 396 | 70 | 23.90 | 2.93 | + | 1.37E-10 |
| ↳asymmetric synapse | 400 | 71 | 24.15 | 2.94 | + | 7.62E-11 |
| ↳neuron to neuron synapse | 427 | 74 | 25.78 | 2.87 | + | 6.44E-11 |
| ↳postsynaptic specialization | 435 | 71 | 26.26 | 2.70 | + | 2.80E-09 |
| dendritic spine | 193 | 34 | 11.65 | 2.92 | + | 3.40E-04 |
| ↳dendrite | 705 | 106 | 42.56 | 2.49 | + | 1.50E-12 |
| ↳dendritic tree | 708 | 106 | 42.74 | 2.48 | + | 1.75E-12 |
| ↳somatodendritic compartment | 1028 | 142 | 62.05 | 2.29 | + | 1.09E-14 |
| ↳neuron spine | 199 | 35 | 12.01 | 2.91 | + | 2.35E-04 |
| perinuclear region of cytoplasm | 657 | 113 | 39.66 | 2.85 | + | 1.53E-17 |
| nuclear speck | 315 | 54 | 19.01 | 2.84 | + | 2.56E-07 |
| ↳ nuclear body | 682 | 83 | 41.17 | 2.02 | + | 2.55E-05 |
| ↳ nucleoplasm | 3324 | 302 | 200.65 | 1.51 | + | 2.01E-09 |
| endocytic vesicle | 191 | 32 | 11.53 | 2.78 | + | 2.00E-03 |
| mitochondrial inner membrane | 438 | 71 | 26.44 | 2.69 | + | 3.66E-09 |
| ↳ mitochondrial membrane | 617 | 95 | 37.24 | 2.55 | + | 1.09E-11 |
| ↳ mitochondrial envelope | 663 | 101 | 40.02 | 2.52 | + | 3.03E-12 |
| ↳ organelle envelope | 1069 | 158 | 64.53 | 2.45 | + | 2.28E-19 |
| ↳ envelope | 1070 | 158 | 64.59 | 2.45 | + | 2.42E-19 |
| ↳ organelle inner membrane | 482 | 74 | 29.10 | 2.54 | + | 1.65E-08 |
| mitochondrial outer membrane | 174 | 28 | 10.50 | 2.67 | + | 2.73E-02 |
| ↳ organelle outer membrane | 192 | 31 | 11.59 | 2.67 | + | 8.57E-03 |
| ↳ outer membrane | 192 | 31 | 11.59 | 2.67 | + | 8.57E-03 |
| sarcomere | 187 | 30 | 11.29 | 2.66 | + | 1.32E-02 |
| ↳ myofibril | 210 | 34 | 12.68 | 2.68 | + | 2.70E-03 |
| ↳ contractile fiber | 224 | 38 | 13.52 | 2.81 | + | 1.60E-04 |
| ↳ supramolecular fiber | 902 | 137 | 54.45 | 2.52 | + | 2.07E-17 |
| ↳ supramolecular polymer | 909 | 138 | 54.87 | 2.52 | + | 2.33E-17 |
| microtubule | 420 | 64 | 25.35 | 2.52 | + | 6.65E-07 |
| ↳ microtubule cytoskeleton | 1173 | 119 | 70.81 | 1.68 | + | 2.56E-04 |
| ↳ polymeric cytoskeletal fiber | 682 | 101 | 41.17 | 2.45 | + | 1.48E-11 |
| glutamatergic synapse | 508 | 73 | 30.66 | 2.38 | + | 3.70E-07 |
| neuronal cell body | 710 | 100 | 42.86 | 2.33 | + | 3.19E-10 |
| ↳ cell body | 801 | 115 | 48.35 | 2.38 | + | 1.28E-12 |
| nuclear envelope | 423 | 59 | 25.53 | 2.31 | + | 5.85E-05 |
| cell projection membrane | 295 | 41 | 17.81 | 2.30 | + | 6.10E-03 |
| apical part of cell | 429 | 51 | 25.90 | 1.97 | + | 3.25E-02 |
| nuclear outer membrane-endoplasmic reticulum membrane network | 566 | 64 | 34.17 | 1.87 | + | 1.03E-02 |
| nucleolus | 808 | 85 | 48.77 | 1.74 | + | 4.40E-03 |
| integral component of plasma membrane | 1499 | 34 | 90.49 | .38 | - | 2.35E-08 |
| ↳ intrinsic component of plasma membrane | 1576 | 37 | 95.13 | .39 | - | 2.22E-08 |
| ↳ intrinsic component of membrane | 6027 | 157 | 363.81 | .43 | - | 6.06E-38 |
| ↳ integral component of membrane | 5854 | 148 | 353.37 | .42 | - | 1.56E-38 |
| Unclassified | 1476 | 27 | 89.10 | .30 | - | 0.00E00 |

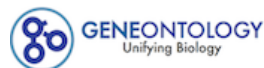

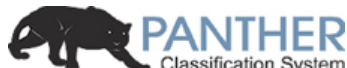

**PANTHER 15.0 released!**Analysis Summary: Please report in publication ?**Analysis Type:** PANTHER Overrepresentation Test (Released 20200407)**Annotation Version and Release Date:** GO Ontology database Released 2020-02-21**Analyzed List:** upload\_1 (Mus musculus)Change**Reference List:** Mus musculus (all genes in database)Change**Annotation Data Set:** GO biological process complete ?**Test Type:** ☒ Fisher's Exact ☐ Binomial**Correction:** ☐ Calculate False Discovery Rate ☒ Use the Bonferroni correction for multiple testing ? ☐ No correction**Results** ?

|  | Reference list | upload_1 |
| --- | --- | --- |
| Uniquely Mapped IDs: | 22265 out of 22265 | 1237 out of 1237 |
| Unmapped IDs: | 0 | 0 |
| Multiple mapping information: | 0 | 0 |

Bonferroni count: 8894

Export Table XML with user input ids JSON with user input ids

Displaying only results for Bonferroni-corrected for P < 0.05, click here to display all results

|  | Mus musculus (REF) | upload_1 (Hierarchy NEW! ?) |  |  |  |  |
| --- | --- | --- | --- | --- | --- | --- |
| GO biological process complete | # | # | expected | Fold Enrichment | +/- | P value |
| fatty acid beta-oxidation using acyl-CoA dehydrogenase | 10 | 9 | .56 | 16.20 | + | 1.53E-03 |
| ↳ fatty acid beta-oxidation | 49 | 25 | 2.72 | 9.18 | + | 2.20E-10 |
| ↳ fatty acid oxidation | 69 | 28 | 3.83 | 7.30 | + | 5.71E-10 |
| ↳ fatty acid metabolic process | 309 | 62 | 17.17 | 3.61 | + | 4.64E-12 |
| ↳ monocarboxylic acid metabolic process | 467 | 100 | 25.95 | 3.85 | + | 4.44E-23 |
| ↳ carboxylic acid metabolic process | 761 | 188 | 42.28 | 4.45 | + | 4.41E-55 |
| ↳ oxoacid metabolic process | 803 | 190 | 44.61 | 4.26 | + | 3.04E-53 |
| ↳ organic acid metabolic process | 829 | 193 | 46.06 | 4.19 | + | 3.08E-53 |
| ↳ cellular metabolic process | 6331 | 699 | 351.74 | 1.99 | + | 1.06E-84 |
| ↳ metabolic process | 7222 | 755 | 401.24 | 1.88 | + | 6.05E-85 |
| ↳ cellular process | 14059 | 1050 | 781.09 | 1.34 | + | 1.46E-57 |
| ↳ small molecule metabolic process | 1432 | 250 | 79.56 | 3.14 | + | 7.11E-50 |
| ↳ organic substance metabolic process | 6701 | 718 | 372.29 | 1.93 | + | 1.18E-82 |
| ↳ cellular lipid metabolic process | 801 | 84 | 44.50 | 1.89 | + | 1.75E-03 |
| ↳ lipid metabolic process | 1063 | 105 | 59.06 | 1.78 | + | 7.40E-04 |
| ↳ primary metabolic process | 6269 | 672 | 348.29 | 1.93 | + | 4.44E-74 |
| ↳ lipid oxidation | 75 | 28 | 4.17 | 6.72 | + | 2.99E-09 |
| ↳ lipid modification | 182 | 35 | 10.11 | 3.46 | + | 2.89E-05 |
| ↳oxidation-reduction process | 787 | 130 | 43.72 | 2.97 | + | 1.68E-21 |
| ↳fatty acid catabolic process | 73 | 28 | 4.06 | 6.90 | + | 1.75E-09 |
| ↳cellular lipid catabolic process | 164 | 32 | 9.11 | 3.51 | + | 9.99E-05 |
| ↳cellular catabolic process | 1492 | 206 | 82.89 | 2.49 | + | 8.77E-27 |
| ↳catabolic process | 1730 | 237 | 96.12 | 2.47 | + | 3.63E-31 |
| ↳lipid catabolic process | 259 | 44 | 14.39 | 3.06 | + | 9.01E-06 |
| ↳organic substance catabolic process | 1433 | 213 | 79.61 | 2.68 | + | 5.44E-32 |
| ↳monocarboxylic acid catabolic process | 93 | 35 | 5.17 | 6.77 | + | 2.75E-12 |
| ↳carboxylic acid catabolic process | 194 | 62 | 10.78 | 5.75 | + | 1.16E-20 |
| ↳organic acid catabolic process | 194 | 62 | 10.78 | 5.75 | + | 1.16E-20 |
| ↳small molecule catabolic process | 290 | 80 | 16.11 | 4.97 | + | 6.88E-24 |
| positive regulation of establishment of protein localization to telomere | 10 | 8 | .56 | 14.40 | + | 1.40E-02 |
| ↳positive regulation of establishment of protein localization | 462 | 66 | 25.67 | 2.57 | + | 6.41E-07 |
| ↳positive regulation of biological process | 6100 | 479 | 338.90 | 1.41 | + | 5.10E-13 |
| ↳regulation of biological process | 11534 | 748 | 640.81 | 1.17 | + | 2.42E-05 |
| ↳biological regulation | 12152 | 801 | 675.14 | 1.19 | + | 1.72E-08 |
| ↳regulation of establishment of protein localization | 771 | 98 | 42.84 | 2.29 | + | 9.12E-09 |
| ↳regulation of protein localization | 1075 | 136 | 59.72 | 2.28 | + | 2.67E-13 |
| ↳regulation of localization | 2886 | 256 | 160.34 | 1.60 | + | 1.91E-09 |
| ↳regulation of cellular protein localization | 560 | 87 | 31.11 | 2.80 | + | 9.66E-12 |
| ↳regulation of cellular localization | 837 | 108 | 46.50 | 2.32 | + | 2.33E-10 |
| ↳positive regulation of cellular protein localization | 308 | 51 | 17.11 | 2.98 | + | 9.52E-07 |
| ↳regulation of establishment of protein localization to telomere | 11 | 8 | .61 | 13.09 | + | 2.30E-02 |
| ↳regulation of establishment of protein localization to chromosome | 12 | 8 | .67 | 12.00 | + | 3.65E-02 |
| viral translation | 14 | 10 | .78 | 12.86 | + | 1.39E-03 |
| ↳viral gene expression | 20 | 11 | 1.11 | 9.90 | + | 2.33E-03 |
| ↳viral process | 131 | 28 | 7.28 | 3.85 | + | 1.51E-04 |
| ↳symbiotic process | 228 | 45 | 12.67 | 3.55 | + | 7.90E-08 |
| ↳interspecies interaction between organisms | 1474 | 127 | 81.89 | 1.55 | + | 2.80E-02 |
| tricarboxylic acid metabolic process | 15 | 10 | .83 | 12.00 | + | 2.21E-03 |
| dicarboxylic acid biosynthetic process | 12 | 8 | .67 | 12.00 | + | 3.65E-02 |
| ↳dicarboxylic acid metabolic process | 81 | 35 | 4.50 | 7.78 | + | 8.91E-14 |
| ↳carboxylic acid biosynthetic process | 224 | 44 | 12.45 | 3.54 | + | 1.54E-07 |
| ↳organic acid biosynthetic process | 225 | 45 | 12.50 | 3.60 | + | 5.36E-08 |
| ↳organic substance biosynthetic process | 2079 | 273 | 115.51 | 2.36 | + | 5.28E-34 |
| ↳biosynthetic process | 2161 | 283 | 120.06 | 2.36 | + | 2.65E-35 |
| ↳cellular biosynthetic process | 1985 | 261 | 110.28 | 2.37 | + | 2.62E-32 |
| ↳small molecule biosynthetic process | 446 | 70 | 24.78 | 2.82 | + | 3.00E-09 |
| regulation of RNA binding | 12 | 8 | .67 | 12.00 | + | 3.65E-02 |
| ↳regulation of molecular function | 2559 | 216 | 142.17 | 1.52 | + | 1.73E-05 |
| negative regulation of mRNA splicing, via spliceosome | 24 | 14 | 1.33 | 10.50 | + | 3.05E-05 |
| ↳regulation of mRNA splicing, via spliceosome | 110 | 46 | 6.11 | 7.53 | + | 1.88E-18 |
| ↳regulation of RNA splicing | 145 | 56 | 8.06 | 6.95 | + | 1.15E-21 |
| ↳regulation of gene expression | 3827 | 334 | 212.62 | 1.57 | + | 7.60E-13 |
| ↳regulation of macromolecule metabolic process | 5484 | 454 | 304.68 | 1.49 | + | 5.80E-16 |
| ↳regulation of metabolic process | 5976 | 501 | 332.01 | 1.51 | + | 6.07E-20 |
| ↳regulation of nucleobase-containing compound metabolic process | 3401 | 253 | 188.95 | 1.34 | + | 2.03E-02 |
| ↳regulation of primary metabolic process | 5349 | 444 | 297.18 | 1.49 | + | 1.06E-15 |
| ↳regulation of nitrogen compound metabolic process | 5198 | 426 | 288.79 | 1.48 | + | 1.19E-13 |
| ↳regulation of cellular metabolic process | 5552 | 461 | 308.46 | 1.49 | + | 1.24E-16 |
| ↳regulation of cellular process | 10826 | 698 | 601.47 | 1.16 | + | 8.92E-04 |
| ↳regulation of mRNA processing | 149 | 56 | 8.28 | 6.76 | + | 3.39E-21 |
| ↳regulation of mRNA metabolic process | 255 | 92 | 14.17 | 6.49 | + | 1.59E-35 |
| ↳negative regulation of RNA splicing | 29 | 16 | 1.61 | 9.93 | + | 4.26E-06 |
| ↳negative regulation of macromolecule metabolic process | 2494 | 238 | 138.56 | 1.72 | + | 1.13E-11 |
| ↳negative regulation of metabolic process | 2766 | 262 | 153.67 | 1.70 | + | 5.81E-13 |
| ↳negative regulation of biological process | 5169 | 436 | 287.18 | 1.52 | + | 2.00E-16 |
| ↳negative regulation of cellular metabolic process | 2469 | 225 | 137.17 | 1.64 | + | 7.46E-09 |
| ↳negative regulation of cellular process | 4632 | 383 | 257.34 | 1.49 | + | 3.62E-12 |
| ↳negative regulation of nitrogen compound metabolic process | 2272 | 204 | 126.23 | 1.62 | + | 4.72E-07 |
| ↳negative regulation of gene expression | 1643 | 166 | 91.28 | 1.82 | + | 8.36E-09 |
| ↳negative regulation of mRNA processing | 32 | 16 | 1.78 | 9.00 | + | 1.28E-05 |
| ↳negative regulation of mRNA metabolic process | 82 | 32 | 4.56 | 7.02 | + | 2.26E-11 |
| stress granule assembly | 16 | 9 | .89 | 10.12 | + | 2.54E-02 |
| ↳cellular component assembly | 1941 | 218 | 107.84 | 2.02 | + | 1.03E-17 |
| ↳cellular component organization | 5045 | 421 | 280.29 | 1.50 | + | 8.72E-15 |
| ↳cellular component organization or biogenesis | 5239 | 433 | 291.07 | 1.49 | + | 9.72E-15 |
| ↳cellular component biogenesis | 2168 | 233 | 120.45 | 1.93 | + | 5.24E-17 |
| ↳organelle organization | 3054 | 278 | 169.67 | 1.64 | + | 6.74E-12 |
| tricarboxylic acid cycle | 32 | 18 | 1.78 | 10.12 | + | 2.74E-07 |
| ↳aerobic respiration | 66 | 20 | 3.67 | 5.45 | + | 1.34E-04 |
| ↳cellular respiration | 129 | 31 | 7.17 | 4.33 | + | 2.27E-06 |
| ↳energy derivation by oxidation of organic compounds | 191 | 43 | 10.61 | 4.05 | + | 5.38E-09 |
| ↳generation of precursor metabolites and energy | 286 | 71 | 15.89 | 4.47 | + | 1.27E-18 |
| alternative mRNA splicing, via spliceosome | 18 | 10 | 1.00 | 10.00 | + | 7.69E-03 |
| ↳mRNA splicing, via spliceosome | 196 | 49 | 10.89 | 4.50 | + | 3.98E-12 |
| ↳mRNA processing | 405 | 101 | 22.50 | 4.49 | + | 6.72E-28 |
| ↳RNA processing | 752 | 136 | 41.78 | 3.26 | + | 4.58E-26 |
| ↳RNA metabolic process | 1232 | 194 | 68.45 | 2.83 | + | 1.13E-31 |
| ↳nucleic acid metabolic process | 1767 | 215 | 98.17 | 2.19 | + | 1.95E-21 |
| ↳macromolecule metabolic process | 5056 | 509 | 280.90 | 1.81 | + | 1.39E-40 |
| ↳nucleobase-containing compound metabolic process | 2191 | 296 | 121.73 | 2.43 | + | 8.29E-40 |
| ↳cellular nitrogen compound metabolic process | 2789 | 420 | 154.95 | 2.71 | + | 1.34E-74 |
| ↳nitrogen compound metabolic process | 5718 | 613 | 317.68 | 1.93 | + | 7.09E-64 |
| ↳organic cyclic compound metabolic process | 2616 | 346 | 145.34 | 2.38 | + | 2.56E-46 |
| ↳heterocycle metabolic process | 2328 | 320 | 129.34 | 2.47 | + | 3.07E-45 |
| ↳cellular aromatic compound metabolic process | 2402 | 325 | 133.45 | 2.44 | + | 8.86E-45 |
| ↳gene expression | 1616 | 270 | 89.78 | 3.01 | + | 3.51E-51 |
| ↳mRNA metabolic process | 531 | 127 | 29.50 | 4.30 | + | 2.08E-34 |
| ↳RNA splicing, via transesterification reactions with bulged adenosine as nucleophile | 196 | 49 | 10.89 | 4.50 | + | 3.98E-12 |
| ↳RNA splicing, via transesterification reactions | 196 | 49 | 10.89 | 4.50 | + | 3.98E-12 |
| ↳RNA splicing | 318 | 85 | 17.67 | 4.81 | + | 9.33E-25 |
| acyl-CoA biosynthetic process | 20 | 11 | 1.11 | 9.90 | + | 2.33E-03 |
| ↳purine ribonucleotide biosynthetic process | 115 | 26 | 6.39 | 4.07 | + | 1.83E-04 |
| ↳purine ribonucleotide metabolic process | 256 | 63 | 14.22 | 4.43 | + | 4.74E-16 |
| ↳ribonucleotide metabolic process | 266 | 64 | 14.78 | 4.33 | + | 6.41E-16 |
| ↳nucleotide metabolic process | 352 | 74 | 19.56 | 3.78 | + | 6.52E-16 |
| ↳nucleoside phosphate metabolic process | 362 | 74 | 20.11 | 3.68 | + | 2.57E-15 |
| ↳phosphate-containing compound metabolic process | 1670 | 155 | 92.78 | 1.67 | + | 1.85E-05 |
| ↳phosphorus metabolic process | 1692 | 156 | 94.00 | 1.66 | + | 2.34E-05 |
| ↳nucleobase-containing small molecule metabolic process | 426 | 85 | 23.67 | 3.59 | + | 1.99E-17 |
| ↳organophosphate metabolic process | 710 | 98 | 39.45 | 2.48 | + | 8.21E-11 |
| ↳ribose phosphate metabolic process | 277 | 66 | 15.39 | 4.29 | + | 2.60E-16 |
| ↳carbohydrate derivative metabolic process | 794 | 95 | 44.11 | 2.15 | + | 5.04E-07 |
| ↳purine nucleotide metabolic process | 273 | 66 | 15.17 | 4.35 | + | 1.35E-16 |
| ↳purine-containing compound metabolic process | 313 | 71 | 17.39 | 4.08 | + | 1.00E-16 |
| ↳organonitrogen compound metabolic process | 4270 | 452 | 237.23 | 1.91 | + | 2.09E-39 |
| ↳ribonucleotide biosynthetic process | 124 | 27 | 6.89 | 3.92 | + | 1.94E-04 |
| ↳nucleotide biosynthetic process | 171 | 36 | 9.50 | 3.79 | + | 2.05E-06 |
| ↳nucleoside phosphate biosynthetic process | 178 | 38 | 9.89 | 3.84 | + | 4.57E-07 |
| ↳organophosphate biosynthetic process | 366 | 51 | 20.33 | 2.51 | + | 2.04E-04 |
| ↳nucleobase-containing compound biosynthetic process | 653 | 68 | 36.28 | 1.87 | + | 3.29E-02 |
| ↳heterocycle biosynthetic process | 718 | 77 | 39.89 | 1.93 | + | 2.27E-03 |
| ↳organic cyclic compound biosynthetic process | 848 | 91 | 47.11 | 1.93 | + | 1.42E-04 |
| ↳aromatic compound biosynthetic process | 734 | 74 | 40.78 | 1.81 | + | 3.47E-02 |
| ↳cellular nitrogen compound biosynthetic process | 1118 | 193 | 62.11 | 3.11 | + | 2.27E-36 |
| ↳ribose phosphate biosynthetic process | 131 | 29 | 7.28 | 3.98 | + | 4.27E-05 |
| ↳purine nucleotide biosynthetic process | 125 | 28 | 6.94 | 4.03 | + | 6.28E-05 |
| ↳purine-containing compound biosynthetic process | 130 | 29 | 7.22 | 4.02 | + | 3.68E-05 |
| ↳organonitrogen compound biosynthetic process | 1070 | 192 | 59.45 | 3.23 | + | 3.01E-38 |
| ↳coenzyme biosynthetic process | 99 | 23 | 5.50 | 4.18 | + | 7.72E-04 |
| ↳coenzyme metabolic process | 204 | 53 | 11.33 | 4.68 | + | 6.13E-14 |
| ↳cofactor metabolic process | 368 | 82 | 20.45 | 4.01 | + | 2.49E-19 |
| ↳cofactor biosynthetic process | 163 | 33 | 9.06 | 3.64 | + | 2.67E-05 |
| ↳amide biosynthetic process | 421 | 135 | 23.39 | 5.77 | + | 6.14E-49 |
| ↳cellular amide metabolic process | 665 | 177 | 36.95 | 4.79 | + | 1.52E-55 |
| ↳ribonucleoside bisphosphate biosynthetic process | 30 | 12 | 1.67 | 7.20 | + | 9.68E-03 |
| ↳ribonucleoside bisphosphate metabolic process | 94 | 32 | 5.22 | 6.13 | + | 4.99E-10 |
| ↳nucleoside bisphosphate metabolic process | 94 | 32 | 5.22 | 6.13 | + | 4.99E-10 |
| ↳nucleoside bisphosphate biosynthetic process | 30 | 12 | 1.67 | 7.20 | + | 9.68E-03 |
| ↳thioester biosynthetic process | 20 | 11 | 1.11 | 9.90 | + | 2.33E-03 |
| ↳thioester metabolic process | 78 | 30 | 4.33 | 6.92 | + | 2.28E-10 |
| ↳sulfur compound metabolic process | 267 | 62 | 14.83 | 4.18 | + | 1.10E-14 |
| ↳sulfur compound biosynthetic process | 72 | 22 | 4.00 | 5.50 | + | 2.26E-05 |
| ↳purine nucleoside bisphosphate biosynthetic process | 30 | 12 | 1.67 | 7.20 | + | 9.68E-03 |
| ↳purine nucleoside bisphosphate metabolic process | 94 | 32 | 5.22 | 6.13 | + | 4.99E-10 |
| ↳acyl-CoA metabolic process | 78 | 30 | 4.33 | 6.92 | + | 2.28E-10 |
| aromatic amino acid family catabolic process | 19 | 10 | 1.06 | 9.47 | + | 1.12E-02 |
| ↳aromatic compound catabolic process | 292 | 56 | 16.22 | 3.45 | + | 6.39E-10 |
| ↳cellular amino acid catabolic process | 90 | 32 | 5.00 | 6.40 | + | 1.86E-10 |
| ↳organonitrogen compound catabolic process | 900 | 109 | 50.00 | 2.18 | + | 5.98E-09 |
| ↳cellular amino acid metabolic process | 231 | 76 | 12.83 | 5.92 | + | 1.21E-26 |
| ↳organic cyclic compound catabolic process | 323 | 64 | 17.95 | 3.57 | + | 2.63E-12 |
| ↳aromatic amino acid family metabolic process | 34 | 13 | 1.89 | 6.88 | + | 5.23E-03 |
| acetyl-CoA metabolic process | 29 | 15 | 1.61 | 9.31 | + | 2.93E-05 |
| glutamate metabolic process | 24 | 12 | 1.33 | 9.00 | + | 1.47E-03 |
| ↳glutamine family amino acid metabolic process | 57 | 20 | 3.17 | 6.32 | + | 1.69E-05 |
| ↳alpha-amino acid metabolic process | 164 | 53 | 9.11 | 5.82 | + | 1.88E-17 |
| translational elongation | 30 | 15 | 1.67 | 9.00 | + | 4.18E-05 |
| ↳cellular macromolecule biosynthetic process | 1193 | 161 | 66.28 | 2.43 | + | 5.38E-19 |
| ↳cellular macromolecule metabolic process | 3958 | 371 | 219.90 | 1.69 | + | 2.72E-20 |
| ↳macromolecule biosynthetic process | 1227 | 161 | 68.17 | 2.36 | + | 7.23E-18 |
| ↳translation | 306 | 111 | 17.00 | 6.53 | + | 9.06E-44 |
| ↳cellular protein metabolic process | 2844 | 279 | 158.01 | 1.77 | + | 4.13E-16 |
| ↳protein metabolic process | 3420 | 318 | 190.01 | 1.67 | + | 7.90E-16 |
| ↳peptide biosynthetic process | 326 | 116 | 18.11 | 6.40 | + | 3.22E-45 |
| ↳peptide metabolic process | 448 | 134 | 24.89 | 5.38 | + | 1.17E-45 |
| tRNA aminoacylation for protein translation | 40 | 20 | 2.22 | 9.00 | + | 1.14E-07 |
| ↳tRNA aminoacylation | 43 | 20 | 2.39 | 8.37 | + | 3.14E-07 |
| ↳amino acid activation | 44 | 20 | 2.44 | 8.18 | + | 4.35E-07 |
| ↳tRNA metabolic process | 165 | 28 | 9.17 | 3.05 | + | 9.79E-03 |
| ↳ncRNA metabolic process | 416 | 55 | 23.11 | 2.38 | + | 2.70E-04 |
| mRNA destabilization | 32 | 16 | 1.78 | 9.00 | + | 1.28E-05 |
| ↳RNA destabilization | 35 | 16 | 1.94 | 8.23 | + | 3.50E-05 |
| ↳regulation of RNA stability | 109 | 42 | 6.06 | 6.94 | + | 1.46E-15 |
| ↳regulation of cellular catabolic process | 698 | 122 | 38.78 | 3.15 | + | 7.56E-22 |
| ↳regulation of catabolic process | 843 | 135 | 46.84 | 2.88 | + | 2.43E-21 |
| ↳posttranscriptional regulation of gene expression | 427 | 133 | 23.72 | 5.61 | + | 5.98E-47 |
| ↳regulation of biological quality | 3959 | 386 | 219.95 | 1.75 | + | 1.36E-24 |
| ↳positive regulation of cellular metabolic process | 3270 | 258 | 181.67 | 1.42 | + | 1.32E-04 |
| ↳positive regulation of cellular process | 5424 | 424 | 301.35 | 1.41 | + | 2.66E-10 |
| ↳positive regulation of metabolic process | 3560 | 288 | 197.79 | 1.46 | + | 9.38E-07 |
| ↳positive regulation of nitrogen compound metabolic process | 3086 | 240 | 171.45 | 1.40 | + | 1.65E-03 |
| ↳positive regulation of macromolecule metabolic process | 3256 | 250 | 180.90 | 1.38 | + | 2.44E-03 |
| ↳positive regulation of cellular catabolic process | 367 | 71 | 20.39 | 3.48 | + | 1.81E-13 |
| ↳positive regulation of catabolic process | 436 | 75 | 24.22 | 3.10 | + | 6.12E-12 |
| ↳positive regulation of mRNA catabolic process | 50 | 23 | 2.78 | 8.28 | + | 1.28E-08 |
| ↳positive regulation of mRNA metabolic process | 93 | 34 | 5.17 | 6.58 | + | 1.47E-11 |
| ↳regulation of mRNA catabolic process | 121 | 48 | 6.72 | 7.14 | + | 1.37E-18 |
| ↳regulation of mRNA stability | 98 | 41 | 5.44 | 7.53 | + | 3.59E-16 |
| ↳negative regulation of translation | 130 | 46 | 7.22 | 6.37 | + | 4.39E-16 |
| ↳negative regulation of cellular protein metabolic process | 1004 | 124 | 55.78 | 2.22 | + | 4.88E-11 |
| ↳regulation of cellular protein metabolic process | 2531 | 252 | 140.62 | 1.79 | + | 1.01E-14 |
| ↳regulation of protein metabolic process | 2723 | 267 | 151.28 | 1.76 | + | 4.78E-15 |
| ↳negative regulation of protein metabolic process | 1072 | 129 | 59.56 | 2.17 | + | 6.67E-11 |
| ↳negative regulation of cellular amide metabolic process | 149 | 48 | 8.28 | 5.80 | + | 1.64E-15 |
| ↳regulation of cellular amide metabolic process | 384 | 110 | 21.33 | 5.16 | + | 2.56E-35 |
| regulation of translation | 333 | 106 | 18.50 | 5.73 | + | 2.43E-37 |
| regulation of cellular macromolecule biosynthetic process | 3333 | 251 | 185.17 | 1.36 | + | 9.09E-03 |
| regulation of macromolecule biosynthetic process | 3419 | 261 | 189.95 | 1.37 | + | 1.77E-03 |
| regulation of biosynthetic process | 3644 | 291 | 202.45 | 1.44 | + | 2.93E-06 |
| regulation of cellular biosynthetic process | 3571 | 275 | 198.40 | 1.39 | + | 3.28E-04 |
| production of miRNAs involved in gene silencing by miRNA | 27 | 13 | 1.50 | 8.67 | + | 6.31E-04 |
| gene silencing by miRNA | 40 | 17 | 2.22 | 7.65 | + | 2.75E-05 |
| posttranscriptional gene silencing by RNA | 47 | 19 | 2.61 | 7.28 | + | 6.38E-06 |
| gene silencing by RNA | 77 | 24 | 4.28 | 5.61 | + | 3.08E-06 |
| posttranscriptional gene silencing | 48 | 20 | 2.67 | 7.50 | + | 1.49E-06 |
| production of small RNA involved in gene silencing by RNA | 29 | 13 | 1.61 | 8.07 | + | 1.21E-03 |
| dsRNA processing | 29 | 13 | 1.61 | 8.07 | + | 1.21E-03 |
| translational initiation | 55 | 25 | 3.06 | 8.18 | + | 1.68E-09 |
| regulation of alternative mRNA splicing, via spliceosome | 66 | 30 | 3.67 | 8.18 | + | 6.54E-12 |
| NADH metabolic process | 27 | 11 | 1.50 | 7.33 | + | 2.36E-02 |
| actin filament capping | 32 | 13 | 1.78 | 7.31 | + | 2.99E-03 |
| negative regulation of actin filament depolymerization | 38 | 15 | 2.11 | 7.10 | + | 5.09E-04 |
| negative regulation of cytoskeleton organization | 161 | 28 | 8.94 | 3.13 | + | 6.37E-03 |
| regulation of cytoskeleton organization | 559 | 69 | 31.06 | 2.22 | + | 8.83E-05 |
| regulation of organelle organization | 1270 | 142 | 70.56 | 2.01 | + | 6.21E-10 |
| regulation of cellular component organization | 2537 | 251 | 140.95 | 1.78 | + | 2.48E-14 |
| negative regulation of cellular component organization | 734 | 80 | 40.78 | 1.96 | + | 8.08E-04 |
| negative regulation of protein depolymerization | 72 | 17 | 4.00 | 4.25 | + | 3.02E-02 |
| regulation of protein depolymerization | 89 | 20 | 4.94 | 4.04 | + | 8.29E-03 |
| regulation of protein-containing complex disassembly | 121 | 27 | 6.72 | 4.02 | + | 1.24E-04 |
| negative regulation of supramolecular fiber organization | 154 | 30 | 8.56 | 3.51 | + | 2.94E-04 |
| regulation of supramolecular fiber organization | 368 | 51 | 20.45 | 2.49 | + | 2.31E-04 |
| regulation of actin filament depolymerization | 51 | 17 | 2.83 | 6.00 | + | 5.10E-04 |
| regulation of actin polymerization or depolymerization | 185 | 31 | 10.28 | 3.02 | + | 3.38E-03 |
| regulation of actin filament length | 188 | 31 | 10.44 | 2.97 | + | 4.60E-03 |
| regulation of cellular component size | 407 | 53 | 22.61 | 2.34 | + | 1.04E-03 |
| regulation of anatomical structure size | 571 | 74 | 31.72 | 2.33 | + | 2.51E-06 |
| negative regulation of actin filament polymerization | 58 | 18 | 3.22 | 5.59 | + | 5.16E-04 |
| regulation of actin filament polymerization | 168 | 29 | 9.33 | 3.11 | + | 4.65E-03 |
| regulation of protein polymerization | 224 | 39 | 12.45 | 3.13 | + | 4.38E-05 |
| regulation of protein-containing complex assembly | 434 | 57 | 24.11 | 2.36 | + | 2.49E-04 |
| regulation of cellular component biogenesis | 964 | 103 | 53.56 | 1.92 | + | 2.15E-05 |
| negative regulation of protein polymerization | 73 | 21 | 4.06 | 5.18 | + | 1.26E-04 |
| negative regulation of protein-containing complex assembly | 136 | 25 | 7.56 | 3.31 | + | 1.02E-02 |
| alpha-amino acid catabolic process | 75 | 30 | 4.17 | 7.20 | + | 9.89E-11 |
| mRNA stabilization | 41 | 16 | 2.28 | 7.02 | + | 2.10E-04 |
| negative regulation of mRNA catabolic process | 51 | 18 | 2.83 | 6.35 | + | 1.01E-04 |
| negative regulation of RNA catabolic process | 61 | 21 | 3.39 | 6.20 | + | 8.89E-06 |
| negative regulation of cellular catabolic process | 241 | 38 | 13.39 | 2.84 | + | 6.99E-04 |
| negative regulation of catabolic process | 301 | 42 | 16.72 | 2.51 | + | 4.37E-03 |
| positive regulation of gene expression | 1995 | 162 | 110.84 | 1.46 | + | 2.68E-02 |
| RNA stabilization | 48 | 18 | 2.67 | 6.75 | + | 4.66E-05 |
| nuclear migration | 31 | 12 | 1.72 | 6.97 | + | 1.28E-02 |
| nucleus localization | 36 | 12 | 2.00 | 6.00 | + | 4.55E-02 |
| organelle localization | 474 | 66 | 26.33 | 2.51 | + | 2.14E-06 |
| cellular localization | 2053 | 249 | 114.06 | 2.18 | + | 1.98E-25 |
| localization | 4864 | 399 | 270.23 | 1.48 | + | 2.00E-12 |
| intracellular transport | 1182 | 171 | 65.67 | 2.60 | + | 1.78E-23 |
| transport | 3572 | 322 | 198.45 | 1.62 | + | 3.81E-14 |
| establishment of localization | 3711 | 332 | 206.18 | 1.61 | + | 2.99E-14 |
| establishment of organelle localization | 331 | 52 | 18.39 | 2.83 | + | 3.30E-06 |
| positive regulation of translation | 123 | 44 | 6.83 | 6.44 | + | 2.03E-15 |
| positive regulation of cellular amide metabolic process | 149 | 48 | 8.28 | 5.80 | + | 1.64E-15 |
| cortical actin cytoskeleton organization | 42 | 15 | 2.33 | 6.43 | + | 1.47E-03 |
| cortical cytoskeleton organization | 59 | 18 | 3.28 | 5.49 | + | 6.41E-04 |
| cytoskeleton organization | 1070 | 118 | 59.45 | 1.98 | + | 1.96E-07 |
| actin cytoskeleton organization | 500 | 68 | 27.78 | 2.45 | + | 2.08E-06 |
| actin filament-based process | 555 | 72 | 30.83 | 2.34 | + | 4.48E-06 |
| monosaccharide biosynthetic process | 34 | 12 | 1.89 | 6.35 | + | 2.80E-02 |
| monosaccharide metabolic process | 156 | 28 | 8.67 | 3.23 | + | 3.65E-03 |
| carbohydrate metabolic process | 405 | 60 | 22.50 | 2.67 | + | 1.26E-06 |
| cellular aldehyde metabolic process | 52 | 18 | 2.89 | 6.23 | + | 1.29E-04 |
| positive regulation of telomere maintenance via telomere lengthening | 35 | 12 | 1.94 | 6.17 | + | 3.58E-02 |
| positive regulation of organelle organization | 596 | 82 | 33.11 | 2.48 | + | 2.13E-08 |
| positive regulation of cellular component organization | 1254 | 140 | 69.67 | 2.01 | + | 7.82E-10 |
| regulation of telomere maintenance | 81 | 19 | 4.50 | 4.22 | + | 8.87E-03 |
| positive regulation of RNA splicing | 50 | 17 | 2.78 | 6.12 | + | 4.02E-04 |
| regulation of translational initiation | 59 | 20 | 3.28 | 6.10 | + | 2.76E-05 |
| alpha-amino acid biosynthetic process | 51 | 17 | 2.83 | 6.00 | + | 5.10E-04 |
| cellular amino acid biosynthetic process | 55 | 18 | 3.06 | 5.89 | + | 2.63E-04 |
| cytoplasmic translation | 64 | 20 | 3.56 | 5.62 | + | 8.69E-05 |
| ribosome assembly | 62 | 19 | 3.44 | 5.52 | + | 2.63E-04 |
| ribonucleoprotein complex biogenesis | 392 | 62 | 21.78 | 2.85 | + | 5.17E-08 |
| pyruvate metabolic process | 69 | 20 | 3.83 | 5.22 | + | 2.49E-04 |
| mitochondrial translation | 52 | 15 | 2.89 | 5.19 | + | 1.41E-02 |
| mitochondrial gene expression | 81 | 21 | 4.50 | 4.67 | + | 5.75E-04 |
| purine ribonucleoside triphosphate metabolic process | 59 | 16 | 3.28 | 4.88 | + | 1.26E-02 |
| purine nucleoside triphosphate metabolic process | 69 | 17 | 3.83 | 4.43 | + | 1.84E-02 |
| nucleoside triphosphate metabolic process | 89 | 19 | 4.94 | 3.84 | + | 2.96E-02 |
| ribonucleoside triphosphate metabolic process | 63 | 17 | 3.50 | 4.86 | + | 6.33E-03 |
| nuclear-transcribed mRNA catabolic process | 94 | 25 | 5.22 | 4.79 | + | 2.12E-05 |
| mRNA catabolic process | 119 | 30 | 6.61 | 4.54 | + | 1.65E-06 |
| RNA catabolic process | 147 | 32 | 8.17 | 3.92 | + | 9.93E-06 |
| nucleobase-containing compound catabolic process | 234 | 37 | 13.00 | 2.85 | + | 9.70E-04 |
| heterocycle catabolic process | 278 | 52 | 15.45 | 3.37 | + | 1.16E-08 |
| cellular nitrogen compound catabolic process | 272 | 51 | 15.11 | 3.37 | + | 1.76E-08 |
| cellular macromolecule catabolic process | 762 | 86 | 42.34 | 2.03 | + | 5.70E-05 |
| macromolecule catabolic process | 858 | 97 | 47.67 | 2.03 | + | 4.15E-06 |
| ribonucleoprotein complex assembly | 163 | 43 | 9.06 | 4.75 | + | 5.12E-11 |
| cellular protein-containing complex assembly | 653 | 83 | 36.28 | 2.29 | + | 6.25E-07 |
|  | 1003 | 127 | 55.72 | 2.28 | + | 2.95E-12 |
| ↳protein-containing complex assembly |  |  |  |  |  |  |
| ↳protein-containing complex subunit organization | 1144 | 142 | 63.56 | 2.23 | + | 2.28E-13 |
| ↳ribonucleoprotein complex subunit organization | 170 | 45 | 9.44 | 4.76 | + | 1.07E-11 |
| mRNA transport | 107 | 28 | 5.94 | 4.71 | + | 3.22E-06 |
| ↳RNA transport | 150 | 37 | 8.33 | 4.44 | + | 2.22E-08 |
| ↳nucleic acid transport | 150 | 37 | 8.33 | 4.44 | + | 2.22E-08 |
| ↳nucleobase-containing compound transport | 183 | 38 | 10.17 | 3.74 | + | 9.18E-07 |
| ↳organic substance transport | 1856 | 202 | 103.12 | 1.96 | + | 1.15E-14 |
| ↳nitrogen compound transport | 1550 | 176 | 86.11 | 2.04 | + | 7.41E-14 |
| ↳establishment of RNA localization | 152 | 37 | 8.44 | 4.38 | + | 3.10E-08 |
| ↳RNA localization | 170 | 42 | 9.44 | 4.45 | + | 7.05E-10 |
| ↳macromolecule localization | 2266 | 253 | 125.89 | 2.01 | + | 4.89E-21 |
| endoplasmic reticulum organization | 58 | 15 | 3.22 | 4.65 | + | 4.39E-02 |
| ↳endomembrane system organization | 367 | 57 | 20.39 | 2.80 | + | 6.96E-07 |
| antibiotic metabolic process | 91 | 23 | 5.06 | 4.55 | + | 2.05E-04 |
| ↳drug metabolic process | 412 | 83 | 22.89 | 3.63 | + | 3.54E-17 |
| maintenance of protein location in cell | 77 | 19 | 4.28 | 4.44 | + | 4.60E-03 |
| ↳maintenance of location in cell | 103 | 24 | 5.72 | 4.19 | + | 3.88E-04 |
| ↳maintenance of location | 173 | 32 | 9.61 | 3.33 | + | 3.01E-04 |
| ↳cellular protein localization | 1423 | 175 | 79.06 | 2.21 | + | 6.72E-17 |
| ↳cellular macromolecule localization | 1430 | 176 | 79.45 | 2.22 | + | 4.67E-17 |
| ↳protein localization | 1963 | 218 | 109.06 | 2.00 | + | 4.39E-17 |
| ↳maintenance of protein location | 108 | 23 | 6.00 | 3.83 | + | 2.96E-03 |
| carbohydrate catabolic process | 88 | 21 | 4.89 | 4.30 | + | 1.90E-03 |
| protein folding | 146 | 34 | 8.11 | 4.19 | + | 6.45E-07 |
| positive regulation of viral process | 82 | 19 | 4.56 | 4.17 | + | 1.04E-02 |
| ↳regulation of viral process | 183 | 32 | 10.17 | 3.15 | + | 9.43E-04 |
| ↳regulation of symbiosis, encompassing mutualism through parasitism | 200 | 36 | 11.11 | 3.24 | + | 7.96E-05 |
| protein export from nucleus | 91 | 21 | 5.06 | 4.15 | + | 3.06E-03 |
| ↳nuclear export | 106 | 21 | 5.89 | 3.57 | + | 2.59E-02 |
| ↳nucleocytoplasmic transport | 203 | 36 | 11.28 | 3.19 | + | 1.12E-04 |
| ↳nuclear transport | 203 | 36 | 11.28 | 3.19 | + | 1.12E-04 |
| ↳intracellular protein transport | 765 | 103 | 42.50 | 2.42 | + | 1.05E-10 |
| ↳protein transport | 1246 | 154 | 69.23 | 2.22 | + | 1.03E-14 |
| ↳establishment of protein localization | 1330 | 164 | 73.89 | 2.22 | + | 7.83E-16 |
| ↳peptide transport | 1276 | 155 | 70.89 | 2.19 | + | 4.03E-14 |
| ↳amide transport | 1301 | 156 | 72.28 | 2.16 | + | 8.27E-14 |
| regulation of viral genome replication | 93 | 20 | 5.17 | 3.87 | + | 1.50E-02 |
| ↳regulation of viral life cycle | 142 | 25 | 7.89 | 3.17 | + | 2.03E-02 |
| ATP metabolic process | 170 | 36 | 9.44 | 3.81 | + | 1.78E-06 |
| endoplasmic reticulum to Golgi vesicle-mediated transport | 105 | 22 | 5.83 | 3.77 | + | 6.73E-03 |
| ↳Golgi vesicle transport | 250 | 46 | 13.89 | 3.31 | + | 3.73E-07 |
| ↳vesicle-mediated transport | 1271 | 128 | 70.61 | 1.81 | + | 6.08E-06 |
| protein stabilization | 174 | 36 | 9.67 | 3.72 | + | 3.09E-06 |
| ↳regulation of protein stability | 278 | 46 | 15.45 | 2.98 | + | 7.97E-06 |
| cellular modified amino acid metabolic process | 156 | 32 | 8.67 | 3.69 | + | 3.51E-05 |
| Golgi organization | 111 | 22 | 6.17 | 3.57 | + | 1.51E-02 |
| cytosolic transport | 132 | 26 | 7.33 | 3.55 | + | 1.97E-03 |
| drug catabolic process | 148 | 29 | 8.22 | 3.53 | + | 4.44E-04 |
| regulation of nucleocytoplasmic transport | 125 | 23 | 6.94 | 3.31 | + | 2.67E-02 |
| ↳regulation of intracellular transport | 352 | 52 | 19.56 | 2.66 | + | 2.45E-05 |
| ↳regulation of transport | 1972 | 183 | 109.56 | 1.67 | + | 4.35E-07 |
| establishment of vesicle localization | 142 | 26 | 7.89 | 3.30 | + | 6.72E-03 |
| ↳establishment of localization in cell | 353 | 46 | 19.61 | 2.35 | + | 5.69E-03 |
| ↳vesicle localization | 154 | 27 | 8.56 | 3.16 | + | 8.72E-03 |
| response to calcium ion | 127 | 23 | 7.06 | 3.26 | + | 3.37E-02 |
| ↳response to metal ion | 270 | 46 | 15.00 | 3.07 | + | 3.47E-06 |
| ↳response to inorganic substance | 417 | 64 | 23.17 | 2.76 | + | 1.09E-07 |
| ↳response to chemical | 3484 | 315 | 193.56 | 1.63 | + | 7.19E-14 |
| cellular carbohydrate metabolic process | 146 | 26 | 8.11 | 3.21 | + | 1.06E-02 |
| receptor-mediated endocytosis | 138 | 24 | 7.67 | 3.13 | + | 3.88E-02 |
| regulation of mitochondrion organization | 140 | 24 | 7.78 | 3.09 | + | 4.81E-02 |
| regulation of cell shape | 158 | 27 | 8.78 | 3.08 | + | 1.35E-02 |
| ↳regulation of cell morphogenesis | 537 | 74 | 29.83 | 2.48 | + | 1.91E-07 |
| ↳regulation of anatomical structure morphogenesis | 1090 | 110 | 60.56 | 1.82 | + | 1.08E-04 |
| ↳regulation of developmental process | 2684 | 211 | 149.12 | 1.41 | + | 5.30E-03 |
| establishment of protein localization to organelle | 302 | 51 | 16.78 | 3.04 | + | 5.12E-07 |
| ↳protein localization to organelle | 622 | 89 | 34.56 | 2.58 | + | 3.00E-10 |
| protein-containing complex localization | 206 | 34 | 11.44 | 2.97 | + | 1.29E-03 |
| protein localization to nucleus | 172 | 28 | 9.56 | 2.93 | + | 2.01E-02 |
| actin filament organization | 241 | 39 | 13.39 | 2.91 | + | 2.56E-04 |
| ↳supramolecular fiber organization | 465 | 63 | 25.83 | 2.44 | + | 1.15E-05 |
| positive regulation of intracellular transport | 198 | 32 | 11.00 | 2.91 | + | 4.50E-03 |
| ↳positive regulation of transport | 1099 | 116 | 61.06 | 1.90 | + | 3.25E-06 |
| cytoskeleton-dependent intracellular transport | 186 | 29 | 10.33 | 2.81 | + | 2.85E-02 |
| positive regulation of protein-containing complex assembly | 221 | 34 | 12.28 | 2.77 | + | 5.50E-03 |
| ↳positive regulation of cellular component biogenesis | 524 | 60 | 29.11 | 2.06 | + | 6.65E-03 |
| positive regulation of cell morphogenesis involved in differentiation | 189 | 29 | 10.50 | 2.76 | + | 3.77E-02 |
| ↳regulation of cell morphogenesis involved in differentiation | 348 | 46 | 19.33 | 2.38 | + | 4.48E-03 |
| ↳regulation of cell development | 1092 | 100 | 60.67 | 1.65 | + | 3.36E-02 |
| regulation of organelle assembly | 207 | 31 | 11.50 | 2.70 | + | 2.79E-02 |
| negative regulation of apoptotic signaling pathway | 232 | 33 | 12.89 | 2.56 | + | 4.15E-02 |
| ↳negative regulation of apoptotic process | 906 | 100 | 50.34 | 1.99 | + | 7.21E-06 |
| ↳negative regulation of programmed cell death | 926 | 100 | 51.45 | 1.94 | + | 2.55E-05 |
| ↳regulation of programmed cell death | 1518 | 141 | 84.34 | 1.67 | + | 1.15E-04 |
| ↳regulation of cell death | 1672 | 159 | 92.89 | 1.71 | + | 2.34E-06 |
| ↳negative regulation of cell death | 1041 | 111 | 57.84 | 1.92 | + | 6.25E-06 |
| ↳regulation of apoptotic process | 1493 | 140 | 82.95 | 1.69 | + | 6.54E-05 |
| regulation of microtubule-based process | 241 | 34 | 13.39 | 2.54 | + | 3.37E-02 |
| positive regulation of protein transport | 443 | 58 | 24.61 | 2.36 | + | 1.89E-04 |
| ↳regulation of protein transport | 739 | 87 | 41.06 | 2.12 | + | 7.59E-06 |
| ↳regulation of peptide transport | 781 | 88 | 43.39 | 2.03 | + | 3.41E-05 |
| organic hydroxy compound metabolic process | 421 | 49 | 23.39 | 2.09 | + | 4.67E-02 |
| regulation of vesicle-mediated transport | 598 | 69 | 33.22 | 2.08 | + | 7.49E-04 |
| cellular response to nitrogen compound | 512 | 57 | 28.45 | 2.00 | + | 3.26E-02 |
| ↳response to nitrogen compound | 851 | 94 | 47.28 | 1.99 | + | 2.34E-05 |
| ↳cellular response to chemical stimulus | 2359 | 217 | 131.06 | 1.66 | + | 1.01E-08 |
| response to organic cyclic compound | 621 | 67 | 34.50 | 1.94 | + | 1.22E-02 |
| ↳ response to organic substance | 2488 | 232 | 138.23 | 1.68 | + | 3.88E-10 |
| response to organonitrogen compound | 745 | 80 | 41.39 | 1.93 | + | 1.08E-03 |
| response to cytokine | 746 | 77 | 41.45 | 1.86 | + | 8.65E-03 |
| regulation of plasma membrane bounded cell projection organization | 781 | 78 | 43.39 | 1.80 | + | 2.23E-02 |
| ↳ regulation of cell projection organization | 790 | 78 | 43.89 | 1.78 | + | 3.58E-02 |
| regulation of hydrolase activity | 994 | 97 | 55.22 | 1.76 | + | 4.06E-03 |
| ↳ regulation of catalytic activity | 1881 | 155 | 104.50 | 1.48 | + | 2.39E-02 |
| cellular homeostasis | 857 | 83 | 47.61 | 1.74 | + | 4.00E-02 |
| ↳ homeostatic process | 1644 | 141 | 91.34 | 1.54 | + | 9.59E-03 |
| cellular response to oxygen-containing compound | 912 | 88 | 50.67 | 1.74 | + | 1.90E-02 |
| ↳ response to oxygen-containing compound | 1296 | 121 | 72.00 | 1.68 | + | 9.75E-04 |
| response to endogenous stimulus | 1126 | 107 | 62.56 | 1.71 | + | 3.26E-03 |
| regulation of response to stress | 1295 | 117 | 71.95 | 1.63 | + | 9.09E-03 |
| positive regulation of molecular function | 1477 | 128 | 82.06 | 1.56 | + | 2.14E-02 |
| cellular response to stress | 1446 | 124 | 80.34 | 1.54 | + | 4.14E-02 |
| ↳ response to stress | 3164 | 237 | 175.79 | 1.35 | + | 2.84E-02 |
| cellular response to organic substance | 1822 | 156 | 101.23 | 1.54 | + | 2.58E-03 |
| regulation of multicellular organismal process | 3223 | 247 | 179.06 | 1.38 | + | 3.36E-03 |
| developmental process | 5570 | 392 | 309.46 | 1.27 | + | 2.82E-03 |
| Unclassified | 1901 | 32 | 105.62 | .30 | - | 0.00E00 |
| G protein-coupled receptor signaling pathway | 1851 | 16 | 102.84 | .16 | - | 1.16E-22 |
| ↳ signal transduction | 4827 | 177 | 268.18 | .66 | - | 1.31E-06 |
| ↳ cell communication | 5260 | 209 | 292.24 | .72 | - | 1.77E-04 |
| ↳ signaling | 5138 | 193 | 285.46 | .68 | - | 2.15E-06 |
| sensory perception of chemical stimulus | 1228 | 1 | 68.23 | .01 | - | 2.84E-24 |
| ↳ sensory perception | 1642 | 13 | 91.23 | .14 | - | 1.19E-20 |
| ↳ nervous system process | 2084 | 44 | 115.78 | .38 | - | 1.14E-10 |

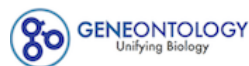

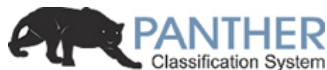

LOGIN REGISTER CONTACT US

Home About PANTHER Data PANTHER Tools PANTHER Services Workspace Downloads Help/Tutorial

PANTHER 15.0 released!

Analysis Summary: Please report in publication ?

**Analysis Type:** PANTHER Overrepresentation Test (Released 20200407)

**Annotation Version and Release Date:** GO Ontology database Released 2020-02-21

**Analyzed List:** upload\_1 (Mus musculus)

Change

**Reference List:** Mus musculus (all genes in database)

Change

**Annotation Data Set:** GO molecular function complete ?

**Test Type:** ☒ Fisher's Exact ☐ Binomial

**Correction:** ☐ Calculate False Discovery Rate ☒ Use the Bonferroni correction for multiple testing ? ☐ No correction

Results ?

|  | Reference list | upload_1 |
| --- | --- | --- |
| Uniquely Mapped IDs: | 22265 out of 22265 | 1237 out of 1237 |
| Unmapped IDs: | 0 | 0 |
| Multiple mapping information: | 0 | 0 |

Bonferroni count: 2756

Export Table XML with user input ids JSON with user input ids

Displaying only results for Bonferroni-corrected for P < 0.05, click here to display all results

|  | Mus musculus (REF) | upload_1 (Hierarchy NEW! ?) |  |  |  |  |
| --- | --- | --- | --- | --- | --- | --- |
| GO molecular function complete | # | # | expected | Fold Enrichment | +/- | P value |
| N6-methyladenosine-containing RNA binding | 8 | 7 | .44 | 15.75 | + | 1.34E-02 |
| ↳ RNA binding | 1086 | 329 | 60.34 | 5.45 | + | 1.79E-125 |
| ↳ nucleic acid binding | 3161 | 371 | 175.62 | 2.11 | + | 1.23E-39 |
| ↳ organic cyclic compound binding | 5175 | 629 | 287.51 | 2.19 | + | 8.23E-89 |
| ↳ binding | 13351 | 1089 | 741.76 | 1.47 | + | 9.87E-98 |
| ↳ heterocyclic compound binding | 5072 | 620 | 281.79 | 2.20 | + | 7.78E-88 |
| acyl-CoA dehydrogenase activity | 10 | 8 | .56 | 14.40 | + | 4.32E-03 |
| ↳ oxidoreductase activity, acting on the CH-CH group of donors | 56 | 16 | 3.11 | 5.14 | + | 2.19E-03 |
| ↳ oxidoreductase activity | 791 | 113 | 43.95 | 2.57 | + | 1.94E-14 |
| ↳ catalytic activity | 5659 | 542 | 314.40 | 1.72 | + | 1.36E-38 |
| RNA stem-loop binding | 14 | 10 | .78 | 12.86 | + | 4.32E-04 |
| translation elongation factor activity | 17 | 11 | .94 | 11.65 | + | 2.12E-04 |
| ↳ translation factor activity, RNA binding | 77 | 39 | 4.28 | 9.12 | + | 4.69E-18 |
| ↳ translation regulator activity, nucleic acid binding | 95 | 45 | 5.28 | 8.53 | + | 3.13E-20 |
| ↳ translation regulator activity | 126 | 59 | 7.00 | 8.43 | + | 5.45E-27 |
| mRNA 5'-UTR binding | 24 | 14 | 1.33 | 10.50 | + | 9.44E-06 |
| ↳ mRNA binding | 272 | 118 | 15.11 | 7.81 | + | 7.46E-54 |
| miRNA binding | 30 | 17 | 1.67 | 10.20 | + | 2.75E-07 |
| ↳ regulatory RNA binding | 43 | 18 | 2.39 | 7.53 | + | 3.55E-06 |
| aminoacyl-tRNA ligase activity | 41 | 20 | 2.28 | 8.78 | + | 4.98E-08 |
| ↳ ligase activity, forming carbon-oxygen bonds | 41 | 20 | 2.28 | 8.78 | + | 4.98E-08 |
| ↳ ligase activity | 146 | 53 | 8.11 | 6.53 | + | 7.39E-20 |
| ↳catalytic activity, acting on a tRNA | 113 | 21 | 6.28 | 3.34 | + | 1.92E-02 |
| ↳catalytic activity, acting on RNA | 337 | 63 | 18.72 | 3.36 | + | 1.50E-11 |
| translation initiation factor activity | 48 | 23 | 2.67 | 8.62 | + | 2.04E-09 |
| RNA helicase activity | 53 | 25 | 2.94 | 8.49 | + | 2.71E-10 |
| ↳helicase activity | 148 | 33 | 8.22 | 4.01 | + | 9.98E-07 |
| ↳ATPase activity, coupled | 279 | 49 | 15.50 | 3.16 | + | 1.14E-07 |
| ↳ATPase activity | 412 | 66 | 22.89 | 2.88 | + | 1.97E-09 |
| ↳nucleoside-triphosphatase activity | 748 | 109 | 41.56 | 2.62 | + | 3.18E-14 |
| ↳pyrophosphatase activity | 801 | 114 | 44.50 | 2.56 | + | 2.53E-14 |
| ↳hydrolase activity, acting on acid anhydrides, in phosphorus-containing anhydrides | 804 | 114 | 44.67 | 2.55 | + | 2.88E-14 |
| ↳hydrolase activity, acting on acid anhydrides | 804 | 114 | 44.67 | 2.55 | + | 2.88E-14 |
| ↳hydrolase activity | 2449 | 231 | 136.06 | 1.70 | + | 3.93E-11 |
| poly(A) binding | 24 | 11 | 1.33 | 8.25 | + | 2.93E-03 |
| ↳poly-purine tract binding | 32 | 16 | 1.78 | 9.00 | + | 3.96E-06 |
| ↳single-stranded RNA binding | 89 | 36 | 4.94 | 7.28 | + | 5.09E-14 |
| double-stranded RNA binding | 82 | 37 | 4.56 | 8.12 | + | 1.03E-15 |
| acid-thiol ligase activity | 27 | 12 | 1.50 | 8.00 | + | 1.23E-03 |
| ↳ligase activity, forming carbon-sulfur bonds | 38 | 14 | 2.11 | 6.63 | + | 8.64E-04 |
| mRNA 3'-UTR binding | 87 | 37 | 4.83 | 7.65 | + | 4.88E-15 |
| poly(U) RNA binding | 26 | 11 | 1.44 | 7.62 | + | 5.45E-03 |
| ↳poly-pyrimidine tract binding | 30 | 13 | 1.67 | 7.80 | + | 5.13E-04 |
| NAD binding | 61 | 25 | 3.39 | 7.38 | + | 3.25E-09 |
| ↳coenzyme binding | 288 | 79 | 16.00 | 4.94 | + | 6.31E-24 |
| ↳cofactor binding | 531 | 105 | 29.50 | 3.56 | + | 1.73E-22 |
| ↳nucleotide binding | 2030 | 301 | 112.78 | 2.67 | + | 1.41E-48 |
| ↳small molecule binding | 2421 | 354 | 134.51 | 2.63 | + | 7.77E-58 |
| ↳nucleoside phosphate binding | 2030 | 301 | 112.78 | 2.67 | + | 1.41E-48 |
| oxidoreductase activity, acting on the CH-NH group of donors | 27 | 11 | 1.50 | 7.33 | + | 7.31E-03 |
| ribosome binding | 64 | 26 | 3.56 | 7.31 | + | 1.36E-09 |
| ↳ribonucleoprotein complex binding | 146 | 45 | 8.11 | 5.55 | + | 2.79E-14 |
| ↳protein-containing complex binding | 1451 | 230 | 80.61 | 2.85 | + | 1.41E-39 |
| rRNA binding | 70 | 28 | 3.89 | 7.20 | + | 2.35E-10 |
| ADP binding | 40 | 16 | 2.22 | 7.20 | + | 4.91E-05 |
| ↳adenyl ribonucleotide binding | 1453 | 203 | 80.73 | 2.51 | + | 1.93E-27 |
| ↳adenyl nucleotide binding | 1464 | 207 | 81.34 | 2.54 | + | 9.78E-29 |
| ↳purine nucleotide binding | 1798 | 252 | 99.89 | 2.52 | + | 1.28E-35 |
| ↳purine ribonucleotide binding | 1786 | 248 | 99.23 | 2.50 | + | 2.58E-34 |
| ↳ribonucleotide binding | 1802 | 251 | 100.12 | 2.51 | + | 5.75E-35 |
| ↳carbohydrate derivative binding | 2123 | 273 | 117.95 | 2.31 | + | 5.69E-33 |
| ↳anion binding | 2703 | 368 | 150.17 | 2.45 | + | 1.79E-53 |
| ↳ion binding | 5485 | 534 | 304.74 | 1.75 | + | 7.81E-40 |
| pre-mRNA binding | 33 | 13 | 1.83 | 7.09 | + | 1.23E-03 |
| oxidoreductase activity, acting on the aldehyde or oxo group of donors, NAD or NADP as acceptor | 47 | 18 | 2.61 | 6.89 | + | 1.10E-05 |
| ↳oxidoreductase activity, acting on the aldehyde or oxo group of donors | 56 | 19 | 3.11 | 6.11 | + | 2.09E-05 |
| Ran GTPase binding | 34 | 12 | 1.89 | 6.35 | + | 8.68E-03 |
| ↳Ras GTPase binding | 399 | 54 | 22.17 | 2.44 | + | 6.84E-05 |
| ↳small GTPase binding | 416 | 55 | 23.11 | 2.38 | + | 8.35E-05 |
| ↳GTPase binding | 515 | 69 | 28.61 | 2.41 | + | 1.02E-06 |
| ↳enzyme binding | 2331 | 273 | 129.51 | 2.11 | + | 7.59E-27 |
| ↳protein binding | 9112 | 791 | 506.24 | 1.56 | + | 4.64E-53 |
| hydro-lyase activity | 52 | 17 | 2.89 | 5.88 | + | 1.99E-04 |
| ↳carbon-oxygen lyase activity | 62 | 18 | 3.44 | 5.23 | + | 3.71E-04 |
| ↳lyase activity | 184 | 38 | 10.22 | 3.72 | + | 3.26E-07 |
| flavin adenine dinucleotide binding | 82 | 25 | 4.56 | 5.49 | + | 6.06E-07 |
| unfolded protein binding | 92 | 28 | 5.11 | 5.48 | + | 5.25E-08 |
| structural constituent of ribosome | 161 | 48 | 8.94 | 5.37 | + | 6.95E-15 |
| ↳structural molecule activity | 576 | 124 | 32.00 | 3.87 | + | 3.82E-30 |
| structural constituent of cytoskeleton | 64 | 19 | 3.56 | 5.34 | + | 1.24E-04 |
| amino acid binding | 71 | 21 | 3.94 | 5.32 | + | 2.60E-05 |
| ↳carboxylic acid binding | 223 | 44 | 12.39 | 3.55 | + | 4.20E-08 |
| ↳organic acid binding | 237 | 45 | 13.17 | 3.42 | + | 7.58E-08 |
| tRNA binding | 60 | 17 | 3.33 | 5.10 | + | 1.10E-03 |
| NADP binding | 47 | 13 | 2.61 | 4.98 | + | 3.14E-02 |
| pyridoxal phosphate binding | 51 | 14 | 2.83 | 4.94 | + | 1.57E-02 |
| ↳vitamin B6 binding | 52 | 14 | 2.89 | 4.85 | + | 1.89E-02 |
| ↳drug binding | 1644 | 229 | 91.34 | 2.51 | + | 1.94E-31 |
| ↳vitamin binding | 132 | 26 | 7.33 | 3.55 | + | 6.11E-04 |
| actin filament binding | 201 | 53 | 11.17 | 4.75 | + | 1.10E-14 |
| ↳actin binding | 426 | 85 | 23.67 | 3.59 | + | 6.16E-18 |
| ↳cytoskeletal protein binding | 976 | 151 | 54.22 | 2.78 | + | 2.13E-23 |
| heat shock protein binding | 140 | 33 | 7.78 | 4.24 | + | 2.89E-07 |
| modified amino acid binding | 100 | 22 | 5.56 | 3.96 | + | 1.02E-03 |
| chaperone binding | 101 | 20 | 5.61 | 3.56 | + | 1.38E-02 |
| oxidoreductase activity, acting on the CH-OH group of donors, NAD or NADP as acceptor | 141 | 27 | 7.83 | 3.45 | + | 5.90E-04 |
| ↳oxidoreductase activity, acting on CH-OH group of donors | 149 | 29 | 8.28 | 3.50 | + | 1.56E-04 |
| single-stranded DNA binding | 111 | 21 | 6.17 | 3.41 | + | 1.51E-02 |
| extracellular matrix structural constituent | 143 | 27 | 7.94 | 3.40 | + | 7.54E-04 |
| ion channel binding | 140 | 23 | 7.78 | 2.96 | + | 4.27E-02 |
| magnesium ion binding | 197 | 32 | 10.94 | 2.92 | + | 1.26E-03 |
| ↳metal ion binding | 3499 | 271 | 194.40 | 1.39 | + | 7.88E-05 |
| ↳cation binding | 3598 | 282 | 199.90 | 1.41 | + | 1.13E-05 |
| ubiquitin protein ligase binding | 308 | 48 | 17.11 | 2.81 | + | 6.12E-06 |
| ↳ubiquitin-like protein ligase binding | 323 | 50 | 17.95 | 2.79 | + | 3.47E-06 |
| GTP binding | 355 | 54 | 19.72 | 2.74 | + | 1.89E-06 |
| ↳purine ribonucleoside triphosphate binding | 1713 | 235 | 95.17 | 2.47 | + | 2.14E-31 |
| ↳purine ribonucleoside binding | 362 | 56 | 20.11 | 2.78 | + | 3.61E-07 |
| ↳ribonucleoside binding | 365 | 57 | 20.28 | 2.81 | + | 1.79E-07 |
| ↳nucleoside binding | 375 | 58 | 20.83 | 2.78 | + | 1.70E-07 |
| ↳purine nucleoside binding | 366 | 56 | 20.33 | 2.75 | + | 7.96E-07 |
| ↳guanyl ribonucleotide binding | 379 | 57 | 21.06 | 2.71 | + | 7.69E-07 |
| ↳guanyl nucleotide binding | 379 | 57 | 21.06 | 2.71 | + | 7.69E-07 |
| protein C-terminus binding | 231 | 35 | 12.83 | 2.73 | + | 1.61E-03 |
| amide binding | 378 | 54 | 21.00 | 2.57 | + | 1.37E-05 |
| ATP binding | 1390 | 191 | 77.23 | 2.47 | + | 1.01E-24 |
| microtubule binding | 247 | 33 | 13.72 | 2.40 | + | 4.78E-02 |
| ↳tubulin binding | 346 | 44 | 19.22 | 2.29 | + | 5.12E-03 |
| phospholipid binding | 427 | 57 | 23.72 | 2.40 | + | 3.52E-05 |
| ↳lipid binding | 763 | 80 | 42.39 | 1.89 | + | 1.01E-03 |
| sulfur compound binding | 255 | 34 | 14.17 | 2.40 | + | 3.54E-02 |
| GTPase activity | 294 | 37 | 16.33 | 2.27 | + | 4.56E-02 |
| protein domain specific binding | 786 | 91 | 43.67 | 2.08 | + | 1.67E-06 |
| protein kinase binding | 727 | 81 | 40.39 | 2.01 | + | 8.75E-05 |
| ↳kinase binding | 813 | 85 | 45.17 | 1.88 | + | 4.42E-04 |
| protein homodimerization activity | 648 | 71 | 36.00 | 1.97 | + | 1.07E-03 |
| ↪ identical protein binding | 1983 | 253 | 110.17 | 2.30 | + | 1.04E-29 |
| ↪ protein dimerization activity | 945 | 89 | 52.50 | 1.70 | + | 1.24E-02 |
| enzyme regulator activity | 989 | 90 | 54.95 | 1.64 | + | 3.80E-02 |
| Unclassified | 2116 | 32 | 117.56 | .27 | - | 0.00E00 |
| DNA-binding transcription activator activity, RNA polymerase II-specific | 472 | 5 | 26.22 | .19 | - | 3.49E-03 |
| ↪ DNA-binding transcription activator activity | 475 | 5 | 26.39 | .19 | - | 3.59E-03 |
| ↪ DNA-binding transcription factor activity | 1066 | 20 | 59.22 | .34 | - | 2.18E-05 |
| ↪ DNA-binding transcription factor activity, RNA polymerase II-specific | 814 | 13 | 45.22 | .29 | - | 1.28E-04 |
| receptor ligand activity | 488 | 5 | 27.11 | .18 | - | 1.76E-03 |
| ↪ signaling receptor activator activity | 493 | 5 | 27.39 | .18 | - | 1.23E-03 |
| ↪ receptor regulator activity | 534 | 6 | 29.67 | .20 | - | 1.16E-03 |
| odorant binding | 473 | 2 | 26.28 | .08 | - | 1.19E-05 |
| G protein-coupled receptor activity | 750 | 1 | 41.67 | .02 | - | 3.38E-13 |
| ↪ transmembrane signaling receptor activity | 2139 | 3 | 118.84 | .03 | - | 2.70E-44 |
| ↪ signaling receptor activity | 2339 | 10 | 129.95 | .08 | - | 3.34E-40 |
| ↪ molecular transducer activity | 2344 | 10 | 130.23 | .08 | - | 3.47E-40 |

**PANTHER 15.0 released!**Analysis Summary: Please report in publication ?**Analysis Type:** PANTHER Overrepresentation Test (Released 20200407)**Annotation Version and Release Date:** GO Ontology database Released 2020-02-21**Analyzed List:** upload\_1 (Mus musculus)Change**Reference List:** Mus musculus (all genes in database)Change**Annotation Data Set:**  ?**Test Type:** ☒ Fisher's Exact ☐ Binomial**Correction:** ☐ Calculate False Discovery Rate ☒ Use the Bonferroni correction for multiple testing ? ☐ No correction**Results** ?

|  | Reference list | upload_1 |
| --- | --- | --- |
| Uniquely Mapped IDs: | 22265 out of 22265 | 1237 out of 1237 |
| Unmapped IDs: | 0 | 0 |
| Multiple mapping information: | 0 | 0 |

Bonferroni count: 1452

Export Table XML with user input ids JSON with user input ids

Displaying only results for Bonferroni-corrected for $P < 0.05$, click here to display all results

|  | Mus musculus (REF) | upload_1 (▼ Hierarchy NEW! ?) |  |  |  |  |
| --- | --- | --- | --- | --- | --- | --- |
| GO cellular component complete | # | # | expected | Fold Enrichment | +/- | P value |
| eukaryotic translation initiation factor 3 complex, eIF3m | 8 | 7 | .44 | 15.75 | + | 7.08E-03 |
| ↳ eukaryotic translation initiation factor 3 complex | 15 | 12 | .83 | 14.40 | + | 5.18E-06 |
| ↳ cytoplasm | 10945 | 1073 | 608.08 | 1.76 | + | 9.25E-160 |
| ↳ cellular anatomical entity | 18638 | 1197 | 1035.49 | 1.16 | + | 8.12E-44 |
| ↳ intracellular | 13691 | 1155 | 760.65 | 1.52 | + | 1.73E-139 |
| ↳ protein-containing complex | 5309 | 588 | 294.96 | 1.99 | + | 1.31E-65 |
| chaperonin-containing T-complex | 10 | 8 | .56 | 14.40 | + | 2.28E-03 |
| ↳ chaperone complex | 22 | 14 | 1.22 | 11.45 | + | 2.16E-06 |
| ↳ cytosol | 3533 | 514 | 196.29 | 2.62 | + | 1.09E-92 |
| aminoacyl-tRNA synthetase multienzyme complex | 12 | 9 | .67 | 13.50 | + | 7.23E-04 |
| zona pellucida receptor complex | 13 | 9 | .72 | 12.46 | + | 1.17E-03 |
| endoplasmic reticulum chaperone complex | 12 | 8 | .67 | 12.00 | + | 5.97E-03 |
| ↳ endoplasmic reticulum | 1639 | 175 | 91.06 | 1.92 | + | 3.21E-12 |
| ↳ endomembrane system |  |  |  |  |  |  |
|  | 3884 | 351 | 215.79 | 1.63 | + | 5.41E-17 |
| ↳ intracellular membrane-bounded organelle | 10291 | 916 | 571.75 | 1.60 | + | 3.24E-80 |
| ↳ intracellular organelle | 11954 | 1033 | 664.14 | 1.56 | + | 1.91E-100 |
| ↳ organelle | 12303 | 1037 | 683.53 | 1.52 | + | 1.86E-93 |
| ↳ membrane-bounded organelle | 11048 | 950 | 613.81 | 1.55 | + | 3.44E-78 |
| messenger ribonucleoprotein complex | 13 | 8 | .72 | 11.08 | + | 9.19E-03 |
| ↳ ribonucleoprotein complex | 703 | 182 | 39.06 | 4.66 | + | 1.60E-56 |
| proton-transporting two-sector ATPase complex, catalytic domain | 14 | 8 | .78 | 10.29 | + | 1.38E-02 |
| cytoplasmic stress granule | 66 | 35 | 3.67 | 9.55 | + | 9.26E-17 |
| ↳ cytoplasmic ribonucleoprotein granule | 206 | 73 | 11.44 | 6.38 | + | 6.16E-28 |
| ↳ ribonucleoprotein granule | 217 | 80 | 12.06 | 6.64 | + | 8.37E-32 |
| ↳ intracellular non-membrane-bounded organelle | 4162 | 444 | 231.23 | 1.92 | + | 4.51E-40 |
| ↳ non-membrane-bounded organelle | 4181 | 445 | 232.29 | 1.92 | + | 6.15E-40 |
| ↳ supramolecular complex | 1197 | 196 | 66.50 | 2.95 | + | 6.23E-35 |
| smooth endoplasmic reticulum | 36 | 17 | 2.00 | 8.50 | + | 1.27E-06 |
| mitochondrial nucleoid | 47 | 22 | 2.61 | 8.43 | + | 4.87E-09 |
| ↳ mitochondrial matrix | 272 | 86 | 15.11 | 5.69 | + | 2.79E-30 |
| ↳ intracellular organelle lumen | 4311 | 438 | 239.51 | 1.83 | + | 3.06E-34 |
| ↳ organelle lumen | 4312 | 438 | 239.57 | 1.83 | + | 3.12E-34 |
| ↳ membrane-enclosed lumen | 4312 | 438 | 239.57 | 1.83 | + | 3.12E-34 |
| ↳ mitochondrion | 1803 | 304 | 100.17 | 3.03 | + | 1.76E-60 |
| ↳ nucleoid | 47 | 22 | 2.61 | 8.43 | + | 4.87E-09 |
| polysomal ribosome | 30 | 14 | 1.67 | 8.40 | + | 4.39E-05 |
| ↳ polysome | 73 | 35 | 4.06 | 8.63 | + | 1.11E-15 |
| ↳ ribosome | 234 | 69 | 13.00 | 5.31 | + | 2.15E-22 |
| cytosolic small ribosomal subunit | 46 | 20 | 2.56 | 7.83 | + | 1.33E-07 |
| ↳ small ribosomal subunit | 78 | 30 | 4.33 | 6.92 | + | 3.72E-11 |
| ↳ ribosomal subunit | 202 | 59 | 11.22 | 5.26 | + | 1.08E-18 |
| ↳ cytosolic ribosome | 117 | 37 | 6.50 | 5.69 | + | 6.05E-12 |
| myelin sheath | 213 | 88 | 11.83 | 7.44 | + | 2.52E-38 |
| proteasome accessory complex | 26 | 10 | 1.44 | 6.92 | + | 1.65E-02 |
| ↳ proteasome complex | 66 | 19 | 3.67 | 5.18 | + | 9.87E-05 |
| ↳ endopeptidase complex | 67 | 19 | 3.72 | 5.10 | + | 1.20E-04 |
| ↳ peptidase complex | 93 | 20 | 5.17 | 3.87 | + | 2.44E-03 |
| ↳ catalytic complex | 1341 | 119 | 74.50 | 1.60 | + | 2.29E-03 |
| neuron projection cytoplasm | 39 | 12 | 2.17 | 5.54 | + | 1.46E-02 |
| ↳ cytoplasmic region | 216 | 31 | 12.00 | 2.58 | + | 1.20E-02 |
| ↳ plasma membrane bounded cell projection | 2281 | 229 | 126.73 | 1.81 | + | 3.86E-14 |
| ↳ cell projection | 2499 | 249 | 138.84 | 1.79 | + | 2.45E-15 |
| ↳ neuron projection | 1516 | 166 | 84.23 | 1.97 | + | 2.14E-12 |
| P-body | 72 | 22 | 4.00 | 5.50 | + | 3.70E-06 |
| catalytic step 2 spliceosome | 83 | 25 | 4.61 | 5.42 | + | 3.95E-07 |
| ↳ spliceosomal complex | 197 | 48 | 10.94 | 4.39 | + | 3.02E-12 |
| ↳nucleus | 6824 | 570 | 379.13 | 1.50 | + | 3.67E-25 |
| intercalated disc | 62 | 18 | 3.44 | 5.23 | + | 1.96E-04 |
| ↳cell-cell contact zone | 85 | 20 | 4.72 | 4.24 | + | 7.26E-04 |
| ↳anchoring junction | 619 | 72 | 34.39 | 2.09 | + | 4.62E-05 |
| ↳cell junction | 2028 | 251 | 112.67 | 2.23 | + | 8.60E-28 |
| stress fiber | 85 | 23 | 4.72 | 4.87 | + | 1.14E-05 |
| ↳contractile actin filament bundle | 85 | 23 | 4.72 | 4.87 | + | 1.14E-05 |
| ↳actin filament bundle | 94 | 28 | 5.22 | 5.36 | + | 4.22E-08 |
| ↳actin cytoskeleton | 497 | 81 | 27.61 | 2.93 | + | 9.99E-13 |
| ↳cytoskeleton | 2135 | 220 | 118.62 | 1.85 | + | 1.54E-14 |
| ↳actomyosin | 96 | 28 | 5.33 | 5.25 | + | 6.37E-08 |
| cytosolic large ribosomal subunit | 68 | 18 | 3.78 | 4.76 | + | 6.22E-04 |
| ↳large ribosomal subunit | 129 | 31 | 7.17 | 4.33 | + | 3.71E-07 |
| extracellular exosome | 80 | 21 | 4.44 | 4.72 | + | 7.83E-05 |
| ↳extracellular vesicle | 88 | 21 | 4.89 | 4.30 | + | 3.10E-04 |
| ↳extracellular organelle | 105 | 21 | 5.83 | 3.60 | + | 3.71E-03 |
| ↳vesicle | 2039 | 173 | 113.28 | 1.53 | + | 1.43E-04 |
| nuclear matrix | 88 | 22 | 4.89 | 4.50 | + | 7.95E-05 |
| ↳nuclear periphery | 113 | 26 | 6.28 | 4.14 | + | 2.19E-05 |
| ↳nuclear lumen | 3899 | 340 | 216.62 | 1.57 | + | 5.65E-14 |
| vesicle coat | 53 | 13 | 2.94 | 4.41 | + | 4.89E-02 |
| ↳coated vesicle membrane | 76 | 16 | 4.22 | 3.79 | + | 3.28E-02 |
| ↳organelle membrane | 2118 | 231 | 117.67 | 1.96 | + | 2.17E-18 |
| ↳cytoplasmic vesicle | 1905 | 153 | 105.84 | 1.45 | + | 1.53E-02 |
| ↳intracellular vesicle | 1910 | 153 | 106.12 | 1.44 | + | 1.96E-02 |
| ↳whole membrane | 1152 | 125 | 64.00 | 1.95 | + | 2.52E-08 |
| ↳coated vesicle | 188 | 30 | 10.44 | 2.87 | + | 2.08E-03 |
| ↳bounding membrane of organelle | 1098 | 106 | 61.00 | 1.74 | + | 2.72E-04 |
| ↳membrane coat | 95 | 18 | 5.28 | 3.41 | + | 3.58E-02 |
| ↳coated membrane | 95 | 18 | 5.28 | 3.41 | + | 3.58E-02 |
| filopodium | 93 | 22 | 5.17 | 4.26 | + | 1.83E-04 |
| ↳actin-based cell projection | 216 | 39 | 12.00 | 3.25 | + | 2.92E-06 |
| mitochondrial ribosome | 90 | 21 | 5.00 | 4.20 | + | 4.27E-04 |
| ↳organellar ribosome | 90 | 21 | 5.00 | 4.20 | + | 4.27E-04 |
| clathrin-coated pit | 67 | 15 | 3.72 | 4.03 | + | 3.15E-02 |
| ↳plasma membrane region | 1218 | 116 | 67.67 | 1.71 | + | 1.31E-04 |
| endoplasmic reticulum-Golgi intermediate compartment | 72 | 16 | 4.00 | 4.00 | + | 1.83E-02 |
| cortical actin cytoskeleton | 99 | 22 | 5.50 | 4.00 | + | 4.63E-04 |
| ↳cortical cytoskeleton | 130 | 29 | 7.22 | 4.02 | + | 6.01E-06 |
| ↳cell cortex | 316 | 62 | 17.56 | 3.53 | + | 1.88E-12 |
| T-tubule | 70 | 15 | 3.89 | 3.86 | + | 4.91E-02 |
| ↳sarcolemma | 162 | 32 | 9.00 | 3.56 | + | 1.26E-05 |
| brush border | 137 | 29 | 7.61 | 3.81 | + | 1.66E-05 |
| ↳cluster of actin-based cell projections | 195 | 33 | 10.83 | 3.05 | + | 1.92E-04 |
| focal adhesion | 156 | 33 | 8.67 | 3.81 | + | 1.68E-06 |
| ↳cell-substrate junction | 168 | 35 | 9.33 | 3.75 | + | 7.55E-07 |
| microvillus | 96 | 20 | 5.33 | 3.75 | + | 3.73E-03 |
| ruffle | 149 | 31 | 8.28 | 3.74 | + | 7.47E-06 |
| ↳cell leading edge | 389 | 69 | 21.61 | 3.19 | + | 3.93E-12 |
| sarcoplasm | 83 | 17 | 4.61 | 3.69 | + | 2.50E-02 |
| peroxisome | 144 | 29 | 8.00 | 3.62 | + | 4.31E-05 |
| ↳microbody | 144 | 29 | 8.00 | 3.62 | + | 4.31E-05 |
| mitochondrial protein complex | 260 | 51 | 14.45 | 3.53 | + | 6.49E-10 |
| U2-type spliceosomal complex | 87 | 17 | 4.83 | 3.52 | + | 4.24E-02 |
| growth cone | 205 | 40 | 11.39 | 3.51 | + | 2.48E-07 |
| ↳site of polarized growth | 213 | 41 | 11.83 | 3.46 | + | 2.11E-07 |
| ↳distal axon | 379 | 51 | 21.06 | 2.42 | + | 1.08E-04 |
| ↳axon | 715 | 90 | 39.72 | 2.27 | + | 2.43E-08 |
| lamellipodium | 166 | 32 | 9.22 | 3.47 | + | 2.10E-05 |
| dendritic spine | 193 | 32 | 10.72 | 2.98 | + | 4.44E-04 |
| ↳dendrite | 705 | 97 | 39.17 | 2.48 | + | 2.17E-11 |
| ↳dendritic tree | 708 | 97 | 39.34 | 2.47 | + | 2.67E-11 |
| ↳somatodendritic compartment | 1028 | 129 | 57.11 | 2.26 | + | 5.64E-13 |
| ↳postsynapse | 727 | 102 | 40.39 | 2.53 | + | 1.70E-12 |
| ↳synapse | 1435 | 195 | 79.73 | 2.45 | + | 4.65E-25 |
| ↳neuron spine | 199 | 33 | 11.06 | 2.98 | + | 2.91E-04 |
| plasma membrane raft | 127 | 21 | 7.06 | 2.98 | + | 4.77E-02 |
| ↳membrane raft | 372 | 47 | 20.67 | 2.27 | + | 1.73E-03 |
| ↳membrane microdomain | 373 | 48 | 20.72 | 2.32 | + | 1.01E-03 |
| ↳membrane region | 386 | 51 | 21.45 | 2.38 | + | 1.52E-04 |
| nuclear speck | 315 | 52 | 17.50 | 2.97 | + | 1.12E-07 |
| ↳nuclear body | 682 | 79 | 37.89 | 2.08 | + | 1.11E-05 |
| ↳nucleoplasm | 3324 | 288 | 184.67 | 1.56 | + | 9.35E-11 |
| mitochondrial inner membrane | 438 | 69 | 24.33 | 2.84 | + | 6.22E-10 |
| ↳mitochondrial membrane | 617 | 91 | 34.28 | 2.65 | + | 4.33E-12 |
| ↳mitochondrial envelope | 663 | 95 | 36.83 | 2.58 | + | 5.30E-12 |
| ↳organelle envelope | 1069 | 148 | 59.39 | 2.49 | + | 7.19E-19 |
| ↳envelope | 1070 | 148 | 59.45 | 2.49 | + | 7.65E-19 |
| ↳organelle inner membrane | 482 | 72 | 26.78 | 2.69 | + | 1.97E-09 |
| collagen-containing extracellular matrix | 366 | 56 | 20.33 | 2.75 | + | 4.20E-07 |
| ↳extracellular matrix | 481 | 58 | 26.72 | 2.17 | + | 3.54E-04 |
| endocytic vesicle | 191 | 29 | 10.61 | 2.73 | + | 7.39E-03 |
| postsynaptic density | 396 | 60 | 22.00 | 2.73 | + | 1.09E-07 |
| ↳asymmetric synapse | 400 | 61 | 22.22 | 2.74 | + | 6.44E-08 |
| ↳neuron to neuron synapse | 427 | 63 | 23.72 | 2.66 | + | 8.05E-08 |
| ↳postsynaptic specialization | 435 | 61 | 24.17 | 2.52 | + | 1.01E-06 |
| perinuclear region of cytoplasm | 657 | 99 | 36.50 | 2.71 | + | 5.69E-14 |
| sarcomere | 187 | 28 | 10.39 | 2.70 | + | 1.35E-02 |
| ↳ myofibril | 210 | 32 | 11.67 | 2.74 | + | 2.29E-03 |
| ↳ contractile fiber | 224 | 35 | 12.45 | 2.81 | + | 4.46E-04 |
| ↳ supramolecular fiber | 902 | 124 | 50.11 | 2.47 | + | 5.48E-15 |
| ↳ supramolecular polymer | 909 | 124 | 50.50 | 2.46 | + | 7.19E-15 |
| organelle outer membrane | 192 | 28 | 10.67 | 2.62 | + | 3.01E-02 |
| ↳ outer membrane | 192 | 28 | 10.67 | 2.62 | + | 3.01E-02 |
| microtubule | 420 | 59 | 23.33 | 2.53 | + | 1.87E-06 |
| ↳ microtubule cytoskeleton | 1173 | 110 | 65.17 | 1.69 | + | 6.30E-04 |
| ↳ polymeric cytoskeletal fiber | 682 | 91 | 37.89 | 2.40 | + | 7.47E-10 |
| nuclear envelope | 423 | 54 | 23.50 | 2.30 | + | 1.77E-04 |
| cell projection membrane | 295 | 37 | 16.39 | 2.26 | + | 2.52E-02 |
| neuronal cell body | 710 | 89 | 39.45 | 2.26 | + | 3.65E-08 |
| ↳ cell body | 801 | 104 | 44.50 | 2.34 | + | 7.08E-11 |
| glutamatergic synapse | 508 | 62 | 28.22 | 2.20 | + | 9.00E-05 |
| endoplasmic reticulum membrane | 540 | 57 | 30.00 | 1.90 | + | 2.25E-02 |
| ↳ nuclear outer membrane-endoplasmic reticulum membrane network | 566 | 62 | 31.45 | 1.97 | + | 2.75E-03 |
| nucleolus | 808 | 83 | 44.89 | 1.85 | + | 7.26E-04 |
| Unclassified | 1476 | 23 | 82.00 | .28 | - | 0.00E00 |
| integral component of plasma membrane | 1499 | 23 | 83.28 | .28 | - | 1.15E-11 |
| ↳ intrinsic component of plasma membrane | 1576 | 26 | 87.56 | .30 | - | 2.04E-11 |
| ↳ intrinsic component of membrane | 6027 | 135 | 334.85 | .40 | - | 3.17E-39 |
| ↳ integral component of membrane | 5854 | 127 | 325.24 | .39 | - | 1.17E-39 |
